# Functional capacities drive recruitment of bacteria into plant root microbiota

**DOI:** 10.1101/2024.08.22.609090

**Authors:** Gijs Selten, Florian Lamouche, Adrián Gómez-Repollés, Zuzana Blahovska, Simon Kelly, Ronnie de Jonge, Simona Radutoiu

## Abstract

Host-associated microbiota follow predictable assembly patterns but show significant variation at the bacterial isolate level depending on the host and environmental context. This variability poses challenges for studying, predicting, and engineering microbiomes. Here we examined how Arabidopsis, Barley, and Lotus plants recruit specific bacteria from highly complex synthetic communities (SynComs) composed of hundreds of bacterial isolates originating from these plants when grown in natural soil. We discovered that, despite their taxonomic diversity, bacteria enriched by these three plant species encode largely overlapping functions. A set of 266 functions common among all host-associated communities was identified at the foundation of the microbiota’s functional potential. Analysis of the differences observed between root-associated communities revealed that functions recruited by Arabidopsis and Barley were primarily driven by the SynCom composition, while Lotus selected fewer isolates but with more diverse functionalities, akin to a ‘Swiss army knife’ strategy. We analysed the variation at the functional level and found this can be explained by the combined functions of bacteria at the family level. Additionally, across major taxa, the isolates covering a broader range of their family’s functional diversity achieved higher relative abundance in the root communities. Our work sheds light on key functions and principles guiding the recruitment of bacterial isolates into root microbiota, offering valuable insights for microbiome engineering and inoculant discovery at a previously inaccessible taxonomic level.

## Main text

Eukaryotes are hosts to microbial communities that have adapted to their unique environments(*1-3*). These microbial communities play a crucial role in maintaining the health and resilience of their hosts(*4, 5*). Therefore, understanding the principles by which microbes are recruited from the environment to form these niche-specific communities is essential for treating dysbiosis, an imbalance or disruption of these microbial communities, and optimizing host health and productivity through microbiome engineering(*6-8*).

Plants recruit commensals and symbionts from established terrestrial ecosystems to facilitate the acquisition of essential nutrients, which support the growth of their photosynthetic and reproductive tissues(*5, 9-12*). In return, plants promote microbial growth by releasing a substantial and diverse portion of their photosynthates into the environment(*13, 14*). This mutual relationship led to the concept of co-evolution of plants and their associated microbes, characterized by continuous interactions and nutritional feedback loops(*15*).

Studies of healthy plants from various evolutionarily diverging taxa and diverse geographic regions have shown that members of five bacterial phyla (Actinobacteriota, Acidobacteriota, Bacteroidota, Firmicutes and Proteobacteria) are consistently enriched in their root microbiota(*16, 17*). This suggests conserved selection mechanisms indicative of a deterministic process. However, when analysing communities at the level of *16S rRNA* or single-copy gene families, a wide taxonomic diversity was uncovered across samples(*16, 18, 19*). This highlights the immense bacterial diversity in soils and in natural ecosystems(*20*) and the extensive range of bacterial functions that plants may recruit for their physiological needs and development.

Deciphering how the initial soil microbiome and various hosts influence the recruitment of plant-associated microbes to maintain homeostasis is a complex task. The diversity at the isolate level and potential ecological interactions makes host-associated microbial communities a biological system that is challenging to understand, predict and engineer(*21, 22*). The concept of selection based on specific microbial functions has often been explored through correlations between microbiome diversity and the functions of taxa identified via amplicon or metagenome sequencing(*1, 17, 18, 23*). Comprehensive and unbiased computational analysis of bacterial genomes has provided initial insights into plant-associated features encoded in bacteria that prefer plants as their niche(*24, 25*). By analysing nearly four thousand bacterial genomes, researchers identified over 3000 functions enriched in plant-associated bacteria compared to non-plant associated bacteria or soil residents(*25*). Parallel, synthetic reconstitution experiments using isolates recruited by model plants have been instrumental in identifying plant and bacterial genes that contribute to microbial community assembly, and uncovered critical principles that drive this process(*26-30*). A preference for the enrichment of endogenous bacterial populations by one host, even amidst competition from those recruited by another host from the same soil environment, has been documented(*31*). Despite significant progress, the principles guiding bacterial selection at the isolate level across different hosts and in different ecosystems remain unknown, largely due to the ecological complexity of these systems(*20*).

In this study, we aimed to identify guiding principles driving the recruitment of bacterial isolates into root microbiota. We investigated the bacterial functionalities enriched by three taxonomically diverse hosts: *Arabidopsis thaliana* (Arabidopsis), *Hordeum vulgare* (Barley), and *Lotus japonicus* (Lotus), all grown in complex microbial environments. Despite being cultivated in different soils, we uncovered a surprisingly extensive overlap in the microbial functions recruited by these plant hosts. Reconstitution experiments using synthetic microbial inocula of unprecedented diversity, combined with extensive computational analysis of root microbiomes, confirmed this overlap. These experiments also revealed ecological principles governing root–microbiota assembly at the isolate level. Out of a vast array of functions encoded by the isolates present in the microbial inocula, only 3%—represented by 266 bacterial functions—were consistently recruited across hosts. Importantly, we found that this minimal set of common bacterial functions enriched in root microbiomes operates as a communal function. This study provides a framework for identifying and characterizing the common and specific bacterial functions that enhance root competitiveness, thereby advancing our understanding and engineering of plant-associated microbiomes.

### Taxonomic and functional breadth of root-associated bacteria

We obtained a culture collection of bacteria associated with the roots of healthy Arabidopsis grown in natural Reijerscamp soil from Utrecht, the Netherlands (AtSC)(*32-34*). We also collected bacteria from Barley and Maize grown in soil from Askov, Denmark (HvSC)(*35*). To assess the extent to which different hosts impose a selection process on their root microbiota, we have Illumina-sequenced and compared the genomes of the isolates from these collections to those previously isolated from the model legume *Lotus japonicus* grown in soil from Cologne, Germany (LjSC)(*31, 35*). The hosts are taxonomically diverse, representing cereals, brassica, and nitrogen-fixing legumes. They exhibit wide variations in their root microbiome interactions: legumes form symbioses with both nitrogen-fixing bacteria and mycorrhizal fungi(*36*), cereals with mycorrhizal fungi only(*37, 38*), and brassica with neither(*31, 35*). Our first insight from this comparison was that each collection contains hundreds of isolates distributed across the most common root-associated phyla and families, maintaining taxonomic separation according to their origin, as previously described(*18, 31, 39-41*) (Figure 1a).

**Figure 1.**
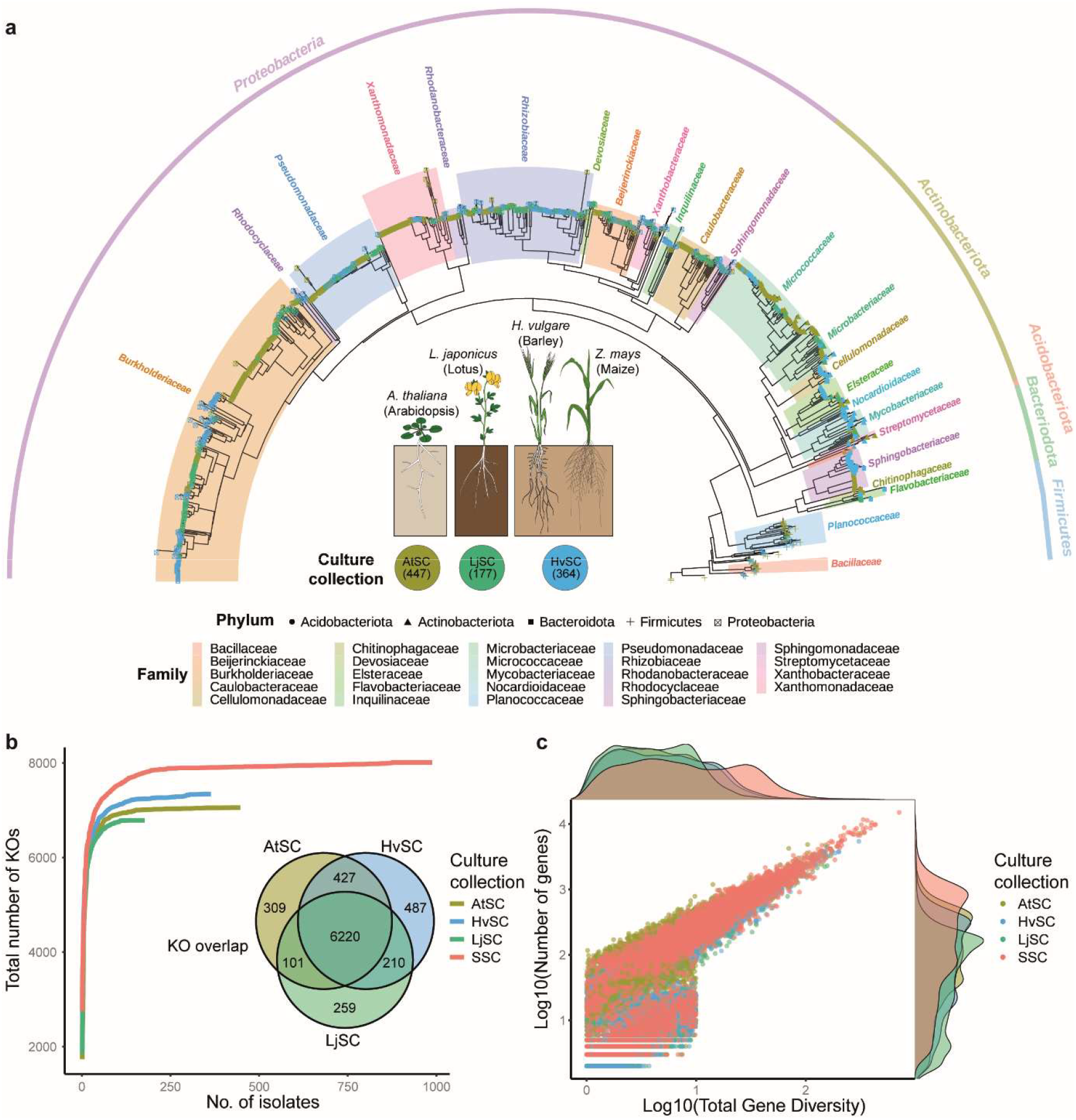
Taxonomic and functional repertoire of the analysed bacterial collections. (a) Phylogenetic tree of rhizobacteria from Arabidopsis, Lotus, Barley/Maize roots, respectively grown in natural soils from the Netherlands (NL-AtSC), Germany (DE-LjSC) and Denmark (DK-HvSC). Distances are calculated based on single-copy gene families derived from Orthofinder (v 2.5.4,(*57*)) and the taxonomy was inferred using the GTDB database (v2.3.0,(*58*)) (b) Total number of KOs identified in each culture collection. SSC (Super SynCom) denotes all three microbial collections together. The overlap of KOs across the three culture collections (AtSC, HvSC or LjSC) is shown in the Venn diagram. (c) Gene sequence variation within KOs. The distribution of number of genes per KO in each of the four microbial collections is shown on the Y-axis. The X-axis displays the gene diversity, calculated by summing all the branch lengths of the gene dendogram of each KO. A higher gene diversity indicates a larger variation in nucleotide sequence between genes in a specific KO

Next, we examined the functional diversity encoded by these genomes. We assigned functions (KO, according to KEGG ontology(*42*)) to genes encoded by each isolate and analysed the genetic variability within a specific KO to determine KO intravariability. We found that 55% of the encoded genes— approximately 3 million—were annotated with a KO, with an average of 370 genes per KO. As expected, an increase in number of isolates in a collection corresponded with an increase in KO diversity, illustrated by an increased number of KOs, gene-copy number, and KO intravariability (Figure 1b, c). The exception was HvSC, which exceeded AtSC in the number of KOs even though it consists of fewer isolates (Figure 1b). Importantly, this analysis revealed that the three collections overlapped in approximately 80% of their KOs (6,255 of 8,013 total KOs), even though they originated from different hosts grown in soils with different properties and were located up to 500 km apart (Table S1). The observed extensive overlap supports the hypothesis that hosts select isolates based on functional similarities, belonging to a diverse range of taxonomic backgrounds. This implies that with our current culture collections, pinpointing bacterial functions that provide isolates with an advantage for association with plant hosts may be achievable.

### Reconstitution experiments using complex synthetic communities

We next tested the hypothesis of functional overlap across plant hosts in controlled studies using complex synthetic communities composed of entire AtSC, LjSC, and HvSC collections, as well as a Super SynCom (SSC) comprised of all three collections together (988 unique genomes) (Figure 2a). We conducted reconstitution experiments by exposing Arabidopsis, Barley, and Lotus to these four inocula. To identify host-dependent community assemblies and minimize the influence of substrate-dependent bacterial enrichment, we used a minimal growth system, cultivating plants for three weeks in a sterile, inert substrate supplemented with low nutrients (Supplementary methods). The entire experiment was performed twice and included a replicate with nutrient-rich growth media (Figure 2a). With few exceptions, we found that the three hosts showed no significant differences in their biomass when exposed to the specific inocula (Figure S1), and that nutrient supplementation had the largest impact on their shoot weight (Figure S1 and S2). Lotus formed nitrogen-fixing nodules with all microbial communities except AtSC, indicating that HvSC isolated from cereal roots contained symbionts compatible with this legume host (Figure S3).

**Figure 2.**
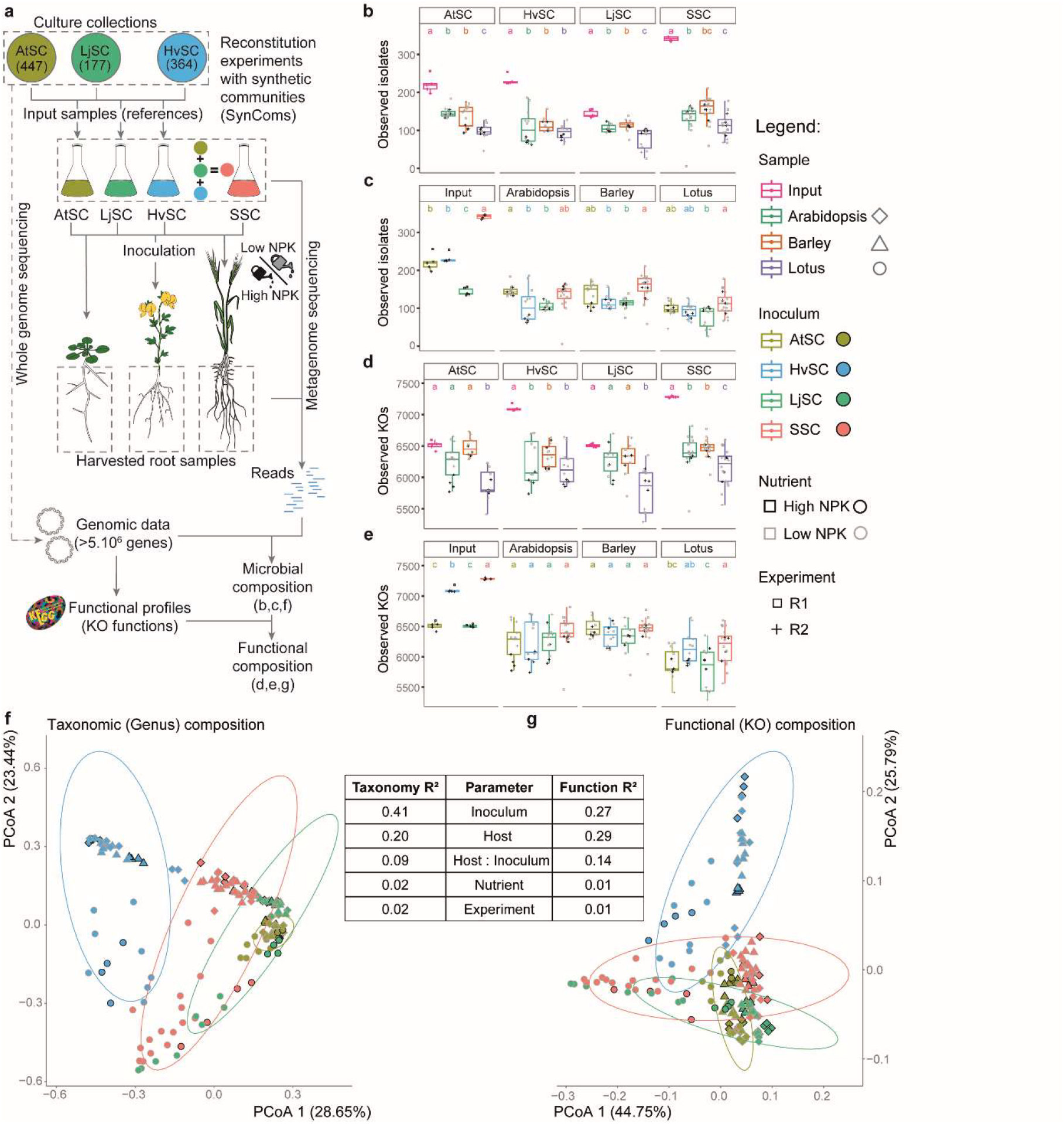
Taxonomic and functional diversities of the root-associated microbial communities from reconstitution experiments. (a) The experimental set-up: The culture collections were fully genome sequenced, with the number of unique genomes displayed between brackets. These collections were then assembled into host-specific SynComs (AtSC, HvSC, LjSC) and the Super SynCom (all three collections combined) to be inoculated with Arabidopsis, Barley, and Lotus. DNA extracted from the root and rhizoplane compartments was subjected to metagenome sequencing, from which the microbiome composition was inferred by quantification of the reads against the microbial culture collection genome sequences. Integration of bacterial genomic data and microbial composition provided the composition of bacterial functions through the KEGG Orthology database (KOs,(*42*)). (b,c,d,e) Alpha diversities: Observed isolates (b,c) and KOs (d,e) in the dataset. Statistical differences were computed using ANOVAs and post-hoc tests (p<0.05) at the SynCom (b,d) or host (c,e) levels. (f,g) Bray-Curtis PCoA on the genus composition (f) and KO composition (g) of Arabidopsis, Barley and Lotus roots (shapes) inoculated with the four SynComs (colors) under high and low nutrient conditions (peripheral colors). R^2^ values in the middle table were computed by PERMANOVA (Adonis test) and indicate the proportion of the compositional differences explained by the distinct parameters. All indicated parameters are significant (p< 0.001). To help the community explore specific KOs and isolates across the different SynComs and hosts in our data, we have created a Shiny app available at this link: https://pm-bacterial-genetics-au.shinyapps.io/SSC_community_app/.

The root-associated microbiota (microbes colonising the root endosphere and rhizoplane) in these reconstitution experiments was investigated by metagenome sequencing and read mapping to the corresponding microbial genomes. This allowed us to computationally distinguish 988 bacterial isolates (Figure S4 and S5). We found that, across hosts and SynComs, a clear selection of the isolates from the different inocula into the root microbiota of Arabidopsis, Barley, and Lotus (Figure 2b & c, S5, S6). Regardless of the initial bacterial richness (Figure 1a), we identified between 76 to 146 isolates (on average 119, each with a relative abundance >0.5%) associated with the roots (Figure 2b). The number of isolates varied significantly with the host and the applied inoculum, and in several host-inoculum combinations it also varied with the nutrient level in the substrate (Figure 2b, Supplementary Results 1, Figure S7). Among hosts, Lotus was the most restrictive, averaging 97 isolates across inocula. Barley was the most permissive, with an average of 134 isolates, while Arabidopsis was intermediate with 123 isolates. This reduction in number of isolates compared to the total number of isolates in the initial inoculum was further confirmed by a Shannon index analysis, which depicted both the richness and evenness within each community (Figure S6). Next, we investigated whether this selection at the isolate level was mirrored at the functional level, both in terms of the number (Figure 2c) and diversity of KOs (Figure 2d, S6). The number and the overall intradiversity of KOs within the selected root communities of all hosts were largely similar across the four inocula (Figure 2c and S6). Root communities of all hosts had a reduced number of KOs compared to the input (Figure S6). The largest reduction at the KO level was observed when plants were inoculated with HvSC or SSC, possibly reflecting stronger impacts of microbe–microbe interactions under these conditions. Together, these results provide evidence that plant roots have a highly selective capacity for accommodating commensal isolates as part of their root microbiota. Importantly, this selectivity remains largely stable despite the complexity of the input SynCom (Figure 2b).

### Functional similarity is recapitulated in reconstitution experiments

Leveraging the extensive diversity of the input inocula and the complex reconstitution experimental setup, we aimed to identify the main drivers of the observed root microbiota. We performed principal coordinate analysis of Bray-Curtis differences between communities at the taxonomic, genus level followed by permutational multivariate analysis of variance (PERMANOVA). This revealed a clear separation of root communities based on input inoculum (R^2^ = 0.415), with a much smaller impact from the host (R^2^ = 0.203) and an even smaller effect from the combination of input inocula and host (R^2^ = 0.090). Nutrient regime or experimental replication had significant but negligible effect on the overall variation (Figure 2-table, Supplementary Results 1, Figure S8). The observed inoculum-dependent separation of root communities aligns with findings from previous studies that analysed root microbiota of plants grown on different soils based on *16S rRNA* amplicon analysis (*43*) confirming the robustness of our experimental approach and analysis.

When performing the same analysis of Bray-Curtis differences at the functional KO level we identified a similar separation to that observed at the genus level, although the individual communities were much more clustered, indicating a reduced variation at the functional compared to taxonomic level (Figure 2g, Figure S8). Together, these results revealed a pattern consistent with the taxonomic and functional composition of the input inoculum derived from the roots of Arabidopsis, Barley and Lotus (Figure 1b), namely functional convergence across hosts and microbial communities. This was further corroborated by the PERMANOVA, which showed a lower impact of the inoculum on KO composition compared to its impact on genus or isolate composition (R^2^ 0.266 vs R^2^ 0.415 and R^2^ 0.436) (Figure 2-table and Figure S9). This analysis also revealed that, across inocula, the individual hosts had a larger impact on the overall functional composition of root communities compared to their impact on the taxonomic composition at genus or isolate level (R^2^ 0.288 vs R^2^ 0.203 and R^2^ 0.129) (Figure 2-table and Figure S9). From this, we conclude that root communities assembled primarily based on bacterial functions that were under host selection. Knowing that the number of KOs (*n* = 8,013) outweighs the number of isolates by tenfold (*n* = 988), we investigated whether this difference could impact our interpretation. To test this, we used the observation that, on average, 119 isolates were associated with the roots of the three hosts (Figure 2b). We performed simulations (*n* = 1,000) using communities of randomly chosen isolates (*n* = 119) from the combined SSC and calculated the correlations between the isolate and KO composition, revealing that observations from actual root microbiota deviate significantly from those obtained from randomly simulated communities (Figure S10). Taken together, these results provide evidence that Arabidopsis, Barley, and Lotus select isolates from the initial microbial community primarily based on the encoded functions, corroborating a deterministic role of the host in the selection of bacterial functions. The impact of host selection is often elusive when communities are analysed at low taxonomic levels, such as at the isolate or genus level using *16S rRNA* amplicons, underscoring the advantage of examining microbiota assembly at the functional level.

### Functional variation is encapsulated at the family level

PERMANOVA revealed that the level of variation detected between root communities when analysed at taxonomic (genus or isolate) level differs from when analysed at functional (KO) level (Figure 2-table). We next explored whether higher taxonomic ranks could better capture the observed variation at the functional level and found that aggregating isolates at the family level closely matched this variation (Figure 3a). The pattern was consistent across all four variables in our experiment—SynCom, host, nutrient level, and experimental replica (Figure 3a)—indicating that the functional diversity of isolates within a family can account for the variation observed in root-associated community functions.

**Figure 3.**
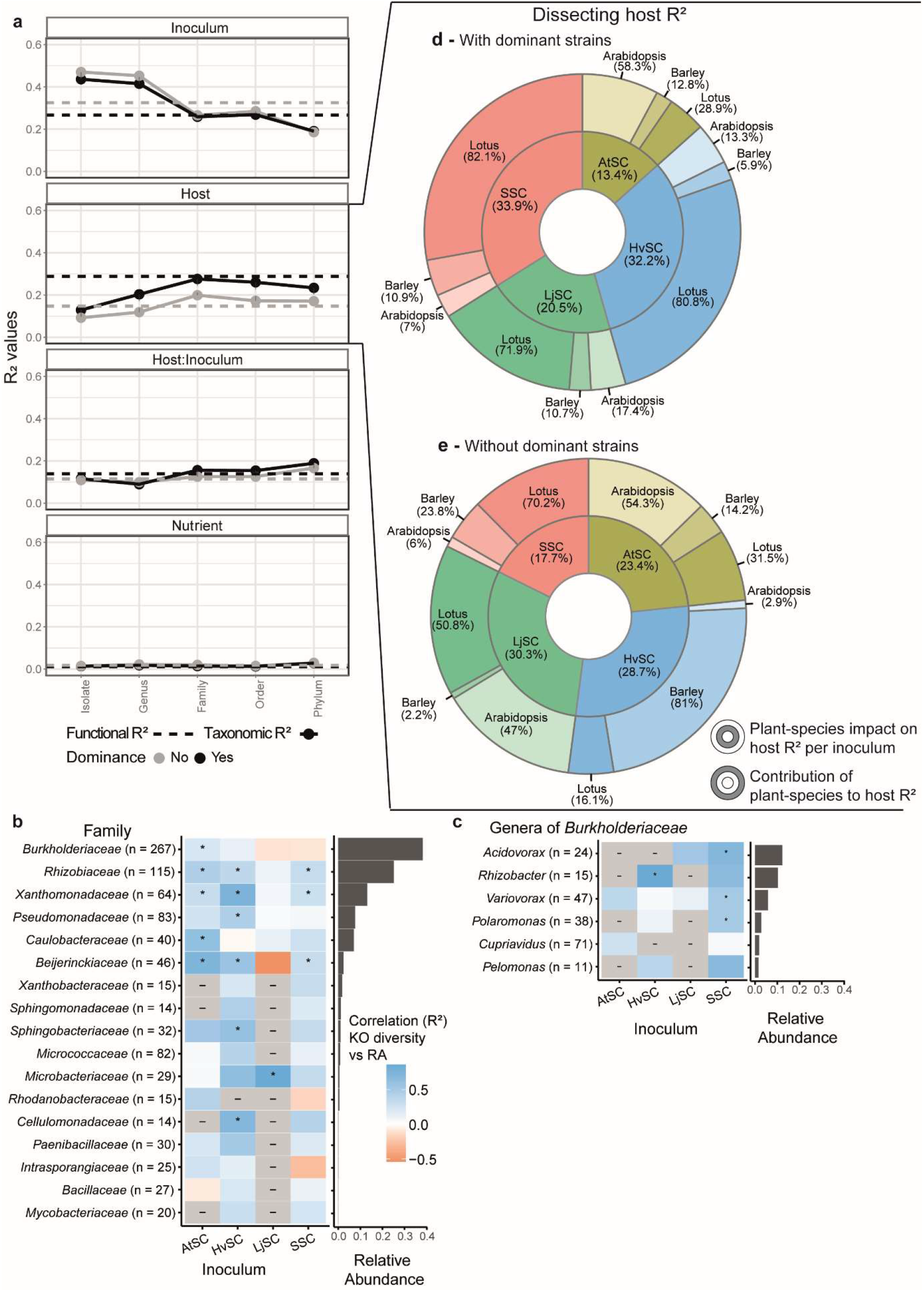
Effect of host and bacterial functional diversity on community assembly. (a) Effect of the inoculum, host, the host:inoculum interaction, and the nutrient condition (from top to bottom) on the variation in taxonomic composition (solid line) across taxonomic ranks, and on the functional composition (dashed line). Grey lines represent data when dominant isolates (the symbionts of Lotus and *Rhizobacter* sp. P2_G4) are excluded from the analyses. The R^2^ values are calculated by a PERMANOVA (Adonis test). (b,c) Correlation heatmaps between the functional diversity of bacterial isolates and their relative abundance in the root microbiome, subsetted per bacterial family. The family relative abundance (0.04-38%) in the root microbiome is presented on the right of the plot and extended to the genus level for the large *Burkholderiaceae* family in panel (c). Significant correlations (p<0.05 after multiple test correction) are displayed with a (*) symbol. Values between brackets indicate the number of isolates in the family, while (-) represents non-computable correlations due to too few isolates in the respective SynCom. (d,e) Piedonut plots dissecting host R^2^ of the original dataset (d) or without the dominant isolates: the symbionts of Lotus and *Rhizobacter* sp. P2_G4 (e). The inner circle represents the combined hosts’ impact on host R^2^ between SynComs, while the outer circle represents the contribution of each host-to-host R^2^ in each inoculum (derived from Figure S16). From panel (d) we can observe that inoculation with SSC, AtSC, HvSC or LjSC, contributed 33.9%, 13.4%, 32.2% and 20.5%, respectively, to the observed effect of the host on the functional R^2^ (0.29). Furthermore, we can conclude that for root communities observed after inoculation with the SSC (red), Lotus, Barley and Arabidopsis contributed, with 82.1%, 10.9% and 7%, respectively to the observed host effect on the functional R^2^. These values were computed based on *in silico* depletion of data from specific SynComs or hosts from the entire dataset and compared to values from the original dataset.

Bacterial families vary greatly in the number of isolates and genera, encoded functions, and cumulative abundance within a community. We therefore investigated whether these cumulative functions are shared among or preferentially present in specific isolates within a family, enabled by the complexity of the inoculum included in our study. For each isolate (*n* = 918) belonging to a total of 17 families and 6 *Burkholderiaceae* genera—together representing the majority of the root communities (90% of the total relative abundance)—we explored whether there is a correlation between the proportion of functions shared by an isolate within its family and its relative abundance in root communities (Figures 3b, c). Interestingly, in the case of nine out of the studied 17 families we found that isolates that encode larger proportions of their respective familial functions also had a larger abundance in root communities compared to the remaining isolates within the same family (Figures S11, S12). This pattern was mainly family-specific and less inoculum-specific, with the isolates from LjSC being a main deviation from the trend. Importantly, these observations were independent of the number of isolates present within the analysed families or bacterial genome completeness (Figures 3b, c, Figure S13). Together, our analyses show that selected root community-level functions are encoded at the family level, and for distinct families, a greater functional diversity within an isolate’s genome increases its likelihood of achieving higher abundance in root communities.

### The host selects bacterial functions

The three plant species included in our studies are taxonomically diverse, and the individual SynComs, although overlapping at the functional level, comprise unique isolates with unique functions (Figure 1). When analysing the root communities observed in our reconstitution experiments, we found that the host and inoculum had the largest contribution to the overall functional variation observed within (Figure 2d). Considering these experimental variables, we investigated whether the three hosts and the communities recruited from the four inocula contributed equally to the overall observed functional variation. To do so, we computationally excluded communities associated with one host at a time or those obtained when using one inoculum at a time. This analysis showed that Lotus-associated communities made the largest contribution to the overall variation, especially when exposed to inocula containing nitrogen-fixing symbionts (>70% variation with HvSC, LjSC, or SSC, compared to 28.9% for Lotus inoculated with AtSC) (Figure 3d, S14a, and S16). Bacterial symbionts, when present, were indeed highly abundant in root communities of Lotus (Figure S5). This indicates that the presence of an active nitrogen-fixing symbiosis had the largest impact on the functional variation identified in the recruited root communities (Figure 3d). By contrast, Barley or Arabidopsis had lower impact on the observed community variation (between 5,9% and 13.3%), with one exception: Arabidopsis in the presence of AtSC where the impact was larger than 50%. This suggests that Lotus and Arabidopsis have a larger impact on the recruited bacterial functions compared to Barley, particularly in the presence of communities that have host-compatible members. For Lotus, these would be the communities with a compatible symbiont, while for Arabidopsis, it is a community previously enriched by Arabidopsis itself.

Parallel to the observed host-effect on communities impacted by the recruitment and enrichment of symbionts by Lotus, our dataset also depicted an inoculum-dependent enrichment of a specific community member, *Rhizobacter* sp. isolate P2_G4 (P2_G4). P2_G4 was highly enriched in root communities of all plant hosts when inoculated with HvSC (Figure S5). This enrichment was significantly reduced when P2_G4 was a member of the SSC, indicating that HvSC provides a microbial context favouring the enrichment of this isolate in the root communities (Figure S5). The function of this bacterium is unknown and is most likely a commensal given no distinct plant phenotypes were observed in the presence of HvSC (Figure S1). Knowing that the presence of distinct members of microbiota can impact the observed variation (Figure 3d), we performed computational depletion of Lotus symbionts or this dominant *Rhizobacter* isolate to determine the effect of the experimental variables in the absence of their functions (absence of functional dominance). This analysis further confirmed the large effect of these isolates on the functional diversity identified within the dataset (Figure 3a, e, Figures S14-S19). Importantly, in their absence, we found each host had the greatest contribution to functional variation in the presence of their native SynComs (Figure 3e).

The SynComs used as inoculum in the reconstitution experiments originated from prior root-driven enrichment from hosts grown in different soils, emphasising the key role of the host in selecting bacterial functions (Figures 1 and 3e). Interestingly, Lotus and Arabidopsis had large effects on the selected functions in the presence of both AtSC and LjSC, in contrast to Barley, whose main impact on KO selection was observed mainly in the presence of HvSC (Figure 3e). Together, our analyses provide evidence that plant hosts select commensals from the microbial environment based on their function and that this selection can be significantly affected by the presence of specific members which are host-enriched (symbionts) or microbial inoculum-dependent (*Rhizobacter* sp. P2_G4). This selection has significant consequences for community composition and the overall enriched bacterial functions.

### Common root-enriched KOs and pathways

Analyses of root communities assembled in our complex reconstitution experiments revealed a clear impact of the inoculum’s composition and host species on the functional enrichment (Figure 3a). Thus, this dataset provides a unique opportunity to pinpoint general functions selected to be part of root microbiota as well as those dependent on these two major determinants. For this, we conducted differential abundance analysis of bacterial KOs and of corresponding pathways enriched in the roots of the three host species versus the input inoculum. This analysis was performed for the entire dataset and when the dominant isolates (Lotus symbionts and *Rhizobacter* sp. P2_G4) were computationally excluded to better disentangle the effect of the SynCom. First, it became clear that, overall, a significant share of the initial KOs (45.7% to 59.3%; i.e. 3500-4000 KOs/7500 total) (Figure 4a, b), was enriched in root communities, an outcome which we tie to the host-selected origin of the isolates from soil. The dominant isolates had a large impact on the overall KO enrichment (Figure S20), emphasising the clear impact of the inoculum. We next explored which pathways are defined by the root-enriched KOs and studied how prevalent these are among isolates. From this, we found that the majority of host-enriched pathways were present in >30% of the isolates, reflecting their host-selected origin (Figure 3e). Nonetheless, even though individual pathways had different levels of enrichment across various host–inoculum combinations (Figure S22), distinct pathways emerged across the combinations (Figures 4e, f). We note that these preferentially enriched pathways varied largely among isolates. Some were prevalent (present in >40% of the isolates) and included specific branches of primary metabolism such as the biosynthesis of nitrogen-donor or sulphur-containing amino acids (aspartate and glutamate), sulphur metabolism, fatty acid biosynthesis, and nicotinamide production, while others represent more rare pathways associated with biofilm and siderophore formation or the metabolism of vitamin B6 and biotin (Figure 4e, f, Supplementary Results 2 – Figures S21, S23). Next, we investigated whether the presence of these pathways confers a competitive advantage to the encoding isolates. To assess this, we compared the cumulative relative abundance of isolates with these pathways to those without, in root environments versus the input inoculum (Figure 4c). In most root communities, isolates had a significantly increased abundance if their genome encoded these pathways, suggesting a competitive advantage (Figure 4e).

**Figure 4.**
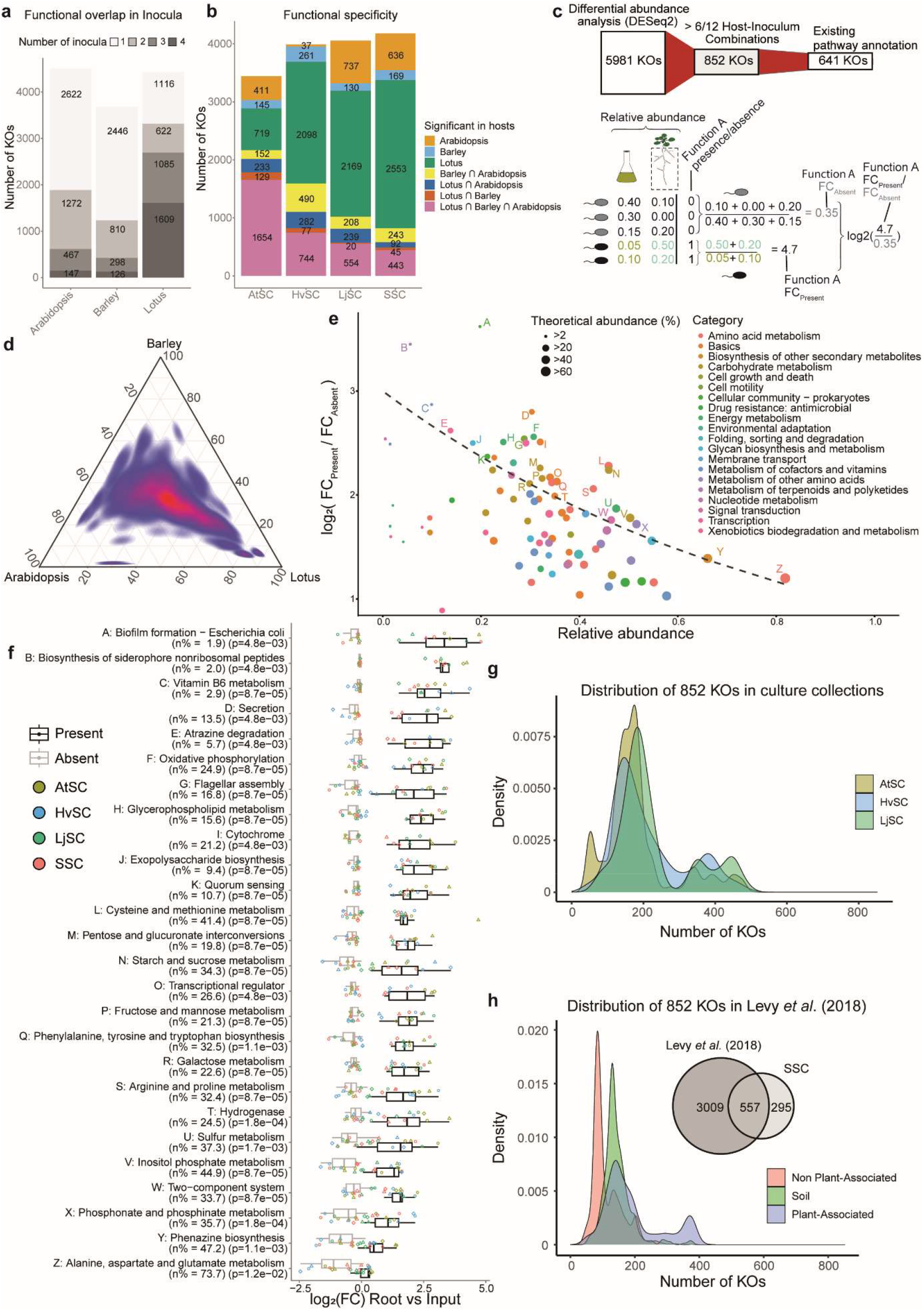
Common functions enriched in root microbiota. (a) Overlap in differentially abundant bacterial KEGG Orthologs (KOs) enriched in the roots of Arabidopsis, Barley, or Lotus compared to the inocula, the four input SynComs AtSC, HvSC, LjSC and SSC. (b) Functional overlap between host species per SynCom, shown by the number of differentially abundant KOs. (c) Sankey diagram (top) of KO enrichment and filtering steps, starting from 5,981 root enriched bacterial KOs to 852 common host-enriched KOs, of which 641 have a pathway annotation. Functional enrichment index (bottom), calculated as the log_2_(ratio) of fold changes derived from a change in cumulative relative abundance of bacterial populations with or without the pathway function between root communities and input SynCom. (d) Density distribution of common, host-enriched KOs (n=852) across the three host species based on the functional enrichment index. (e) Pathways selected by the host identified by differential abundance of the associated KOs, reflected in the cumulative relative abundance of the isolates encoding these pathways (X-axis) and the functional enrichment index combined across the three hosts (Y-axis). Notable pathways have been selected using an exponential decay function maximizing functional enrichment index together with the cumulative relative abundance of the pathways. These pathways are tagged by letters in order of their functional enrichment index (high to low). Size of the pathway dots indicate the proportion of isolates that have the respective pathway in the SSC inoculum and thus represents the theoretical abundance, and the colour indicates the category to which the KOs are assigned. (f) Boxplots of the log_2_(fold changes) derived from a change in cumulative relative abundance of bacteria with (black boxes) or without (grey boxes) the given pathway in root communities versus the respective input SynComs. (g) Density plot showing the distribution of the 852 common host-enriched KOs in the three SynCom collections used in this study. (h) Density plot showing the distribution of the 852 common host-enriched KOs in published bacterial genomes that were categorized as soil, non-plant associated or plant-associated bacteria(*25*). The Venn diagram displays the overlap between the identified plant-associated KO functions in this study and from this collection of published genomes described in Levy *et al*. (2018)(*25*).

The differential abundance analysis revealed a specific pattern where a significant proportion of KOs were enriched across multiple hosts and host-inoculum combinations (Figure 4a, b), confirming the KO overlap observed in the SynCom collections (Figure 1B). Notably, the isolates encoding root-enriched KOs exhibited enrichment across the three host species (Figure 4c), demonstrating functional convergence. Further stringent analysis identified 852 KOs enriched in more than six host-inoculum combinations (Figure 4c), with 266 KOs enriched across all 12 combinations—representing about 3% of the total number of functions encoded by the 988 isolates (Figure S23). This consistent enrichment across multiple host-SynCom combinations underscores the robustness of the functional enrichment observed in our study.

We inspected the isolates from our collections for the presence of these common KOs and found that none contained them all, few encode for approximately half, and the majority had fewer than a quarter (Figure 4g, S23g). This shows that these general functions recruited by plant roots were not isolated actions of single microbes but rather the collective outcome of interactions within the microbial communities during the establishment of the observed root microbiota. Next, we compared our findings with those from a comprehensive and unbiased meta-analysis conducted by Levy *et al*. (2018), which examined 3,837 GenBank-deposited bacterial genomes(*25*). Most of the general plant-enriched KOs identified here (771 of the 852 and 228 of the 266) were also present in genomes analysed by Levy *et al*. (2018). Specifically, 557 of 852 and 177 of 266 KOs overlapped with the >3,000 KOs identified by the authors to be plant-associated(*25*), emphasising their robust enrichment during selection of plant-associated bacteria. Additionally, we examined the isolates included in this previous study(*25*) for the prevalence of the general KOs identified here and observed a similar distribution pattern—the isolates recruited by plants contained a larger proportion of these KOs compared to soil and non-plant-associated bacteria (Figure 4h, S23h). Closer inspection of the identified common KOs revealed functions related to carbohydrate and iron transporters, quorum sensing, two-component systems, exopolysaccharide biosynthesis, and carbohydrate metabolism. These functions are known to be important for plant–microbe interactions(*44-46*) and while they are not novel *per se*, the extent of our experimental studies and the stringent and overarching selection criteria applied here suggest that these could be considered key functions for bacterial survival in complex communities associated with plant roots.

### Communities of Arabidopsis and Barley are conditionally assorted

Next, we searched for KOs that were specifically enriched by a particular host to identify possible host-preferred bacterial functions within their root communities. We found a small number of KOs to be highly enriched (Figure 5a; FC>3 in at least three out of four inocula) in communities of Arabidopsis (n=40), Barley (n=48), or Lotus with (n=355) or without the symbionts (n=61) (Figure 5b). For Arabidopsis, we found genes involved in biosynthesis of bacteriochlorophyllides by photosynthetic bacteria(*47*) to be enriched, corroborating the enrichment of isolates belonging to the *Rhodobacter* genus in this host (Supplementary Results 3 – Figure S24 – Table S9). KOs corresponding to enzymes that add pyruvate (exoV) or acetyl groups (exoZ) to the succinoglycan polysaccharide chain(*48*) were found to be enriched in Barley (Supplementary Results 3 - Figure S25 – Table S10), while those for the catabolism of erythritol(*49*), biosynthesis of Nod factors (nod), and nitrogen fixation (nif)(*50*) were enriched by Lotus (Supplementary Results 3 - Figure S26-28, Table S11 and S12). Together, this indicates a moderate but distinguishable effect of the specific hosts on the recruitment of bacteria based on specific functions.

**Figure 5:**
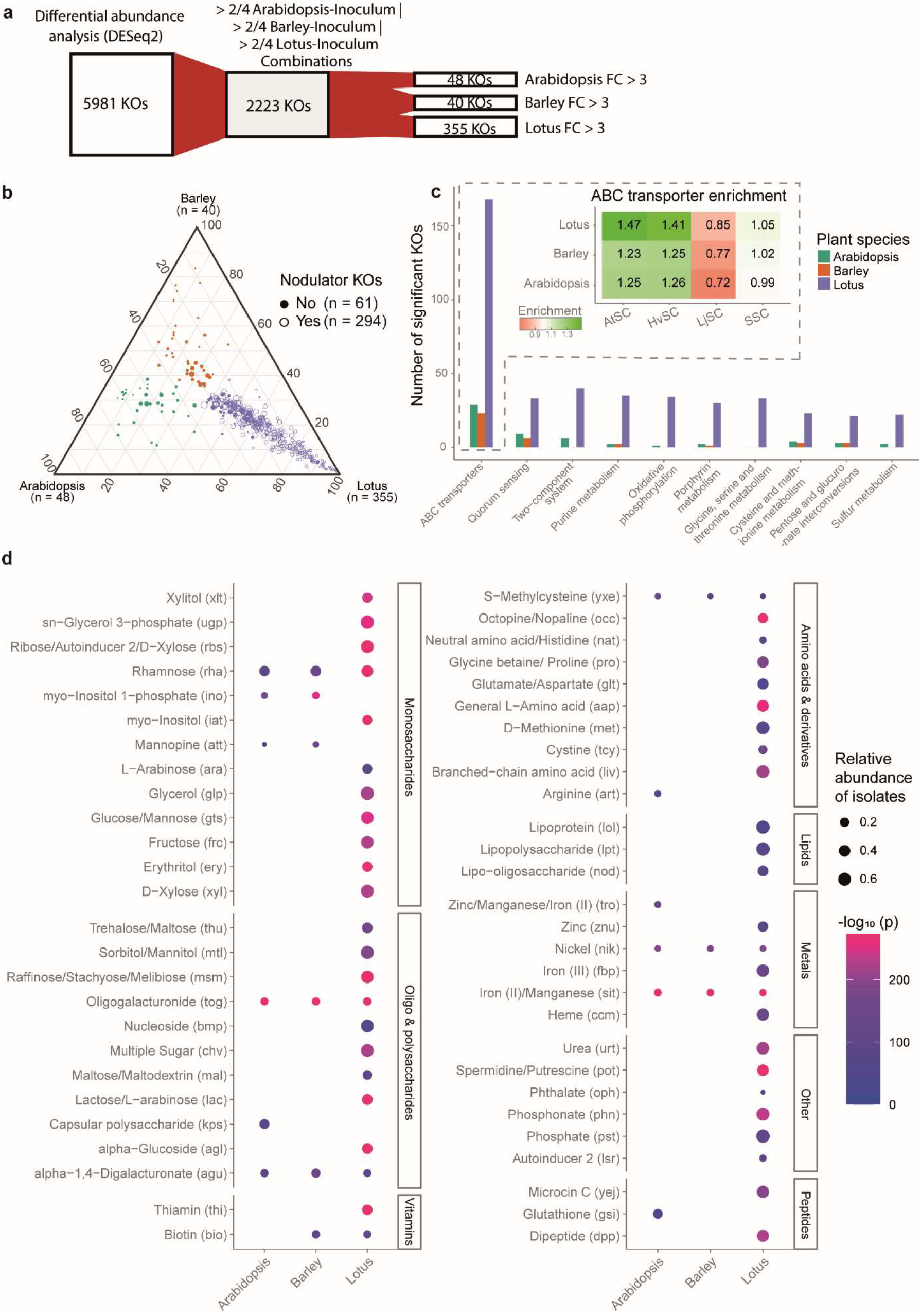
Host-specific functions are enriched by ABC transporters. (a) KOs were considered host-specific if the functional enrichment index was >3 for one host while being <3 for the other two hosts.The functional enrichment index for each host was calculated by taking the median value for each host across the four SynComs. (b) The position of the functional enrichment indices of these KOs in the ternary plane are indicative of the host-specificity. For Lotus, specifically, we performed the same analysis also after removing genes of the Lotus symbionts. Lotus-specific KOs in the dataset with the Lotus symbionts included are indicated by the closed coordinates while the open coordinates indicate KOs that are Lotus-specific when the symbionts gene were excluded. The number of host-specific KOs is added in the corners of the host-specific ternary plot. The number of Lotus-specific KOs in the dataset without the symbionts is 61 as compared to 355 when they are included. (c) Top 10 significant pathways with most host-specific KOs (X-axis) and an indication of how many KOs are Arabidopsis, Barley and Lotus-specific (Y-axis). The heatmap depicts the correlation between isolates’ number of unique ABC transporter KOs and their relative abundance on the root for each combination of host and inoculum revealing a clear enrichment of ABC transporters in abundant isolates in the root microbiome for the AtSC and HvSC condition, while LjSC displays the opposite. (d) Host-specific ABC transporters with observed host-specific enrichment, categorized by the putative substrate they transport across the cell membrane. Multiple gene cassettes that together make up an ABC transporter were combined for this analysis. Dot size represents the cumulative relative abundance of isolates carrying these ABC transporters in their genomes (average of gene cassettes) and the colour gradient is an indication of the significance of differential abundance (DESeq2 p-value) between root communities and the inoculum (averaged by Stouffer’s method(*59*) across gene cassettes).

Over 50% of KOs enriched in root communities of Arabidopsis or Barley, but significantly fewer in Lotus (25%), were found exclusively in the presence of a specific inoculum, referred to as singular KOs (Figure 4a; lightest grey bars). This suggests that the first two host species are more versatile and can select necessary functions depending on the microbial environments. Alternatively, they might exert a lower influence on the selected KOs, in which case these KOs would emerge in the root communities primarily because of microbe-microbe interactions within the individual communities. To assess the host selection’s impact on KO recruitment, we quantified the percentage of KOs that were enriched by the individual hosts in the presence of individual SynComs (HSC) and were again enriched when the same host was exposed to the SSC containing all KOs. Arabidopsis and Barley selected between 19.6-47.0% and 17.7-54.5% HSC-enriched KOs from the SSC, respectively, whereas Lotus reselected a significantly higher proportion of 69.6-91.2% (Table S8). Conversely, we examined the percentage of SSC-enriched KOs that were not enriched in the corresponding HSC. A similar pattern emerged: a significantly larger proportion of SSC-enriched KOs by Arabidopsis and Barley (on average 48.9 and 40.6%, respectively) were not selected in the HSCs, while this proportion was much lower on Lotus, averaging 18.8% (Table S8). These results suggest that functions in the root communities of Arabidopsis and Barley are less influenced by the host and are more likely to depend on other factors, such as microbe-microbe interactions during microbiota establishment.

### Lotus is a more restrictive host, but its root communities are functionally diverse

Microbial communities selected by Lotus, on the other hand, were enriched with KOs that were present across multiple, even all four inocula (1,609 out of total 4,412 KOs) (Figure 4a). Additionally, these communities contain isolates with a broader range of host-specific functions (Figure 5b). This pattern, observed even in the absence of symbionts (Lotus inoculated with AtSC) or when symbionts were excluded from the analysis (Figure S23), suggests that it is not solely related to symbiosis. Knowing that Lotus was more selective, as indicated by the fewer isolates enriched in its root microbial communities (Figure 2b), indicates that Lotus recruits isolates with a more diverse functional repertoire compared to Arabidopsis and Barley (Figure 5b and Tables S11, S12). We reasoned that if Lotus has a different selection strategy compared to the other two host species, one might expect that isolates from LjSC would be functionally better equipped to colonise the roots of the other two hosts. Indeed, we found that when all hosts were inoculated with the SSC, the combined inoculum from all three collections, the LjSC isolates were predominant in all root communities compared to those originating from HvSC or AtSC (Figure S4). The enhanced dominance of LjSC isolates might be due to the large number and diversity of bacterial transporters encoded in their genomes (Figure 5c, d, Supplementary results 4 - Figure S29-32), potentially enhancing their ability to uptake/secrete and catabolize a wide range of metabolites thus improving their survival and growth on plant roots.

## Discussion

Identifying factors that contribute to recruitment of isolates from the environment as part of a specific host microbiota remains a challenge for understanding and engineering microbial ecosystems(*7*). In this study, we identified general principles (Figure 3a, b) and delineated common and host-specific bacterial functions (Figures 4, 5) that facilitate the recruitment of isolates into microbiota of taxonomically diverse plant species. Notably, we found that plants analysed here repeatedly recruited isolates which, despite being taxonomically distinct, had a large overlap in their encoded functions, indicating that these functions enable bacteria to adapt to a plant-host-associated lifestyle. A similar pattern of functional convergence has been found in mouse and human gut microbiota, emphasizing the concept of hosts selecting bacteria based on their functions, rather than taxonomic identity(*51*). We examined large collections of bacterial isolates that were isolated from three different plant species grown on three different soils (Figure 1a). These isolates were further studied in complex community reconstitution experiments, allowing us to disentangle the impact of the host and the inoculum’s composition on functional diversity, independent of the confounding effects of the soil. This revealed guiding principles for the assembly of bacterial functions within root microbiota (Figure 3). Irrespective of the inoculum’s composition or the host, the functional variation within root communities aligns with family-level taxonomic variation. Previous theoretical studies(*52*) have also shown convergence of community composition at the family level and that is strongly influenced by the nature of limiting nutrients. In our study, the inocula contained families with up to 267 distinct isolates allowing us to demonstrate that the impact of the host or inoculum on root communities is reflected in the collective functions of the enriched isolates within a family. This provides empirical support for the concept that a family’s metabolic abilities shape the overall community composition. The extensive diversity captured within the experimental design of this study revealed that isolates covering a broad functional spectrum within their family-level diversity generally had a better capacity for colonisation. Our results are consistent with recent findings on pathogen colonisation resistance in the mouse gut(*53*), where it was observed that a pathogen could succeed in colonisation if it could fill metabolic functions missing in the gut community. Building on these findings and our own, we envision that root commensals within a family will likely occupy available metabolic niches, thereby achieving an optimal balance between the necessary functions at the community level and the number of isolates capable of covering these functions. Interestingly, isolates from LjSC encode a relatively larger number of plant-associated functions, including numerous and diverse ABC transporters (Figure 5), and these isolates also demonstrated increased competitiveness compared to those from AtSC and HvSC. Given the origin of the three collections (three hosts grown on three soils), our study cannot distinguish between the effect of the original soil or host on the selection of isolates within LjSC. Overall, our study revealed that functional versatility within an isolate ensured better survival across different hosts, a key insight for developing biological-based inoculants or designing SynComs for microbiota studies.

We used here an experimental strategy of unprecedented scale, where the three plant hosts were exposed to functionally complex and taxonomically diverse inocula across two nutrient regimes. These experiments revealed assembly of communities with dominant isolates that were either host-driven or inoculum composition-driven. Irrespective of the functions encoded by these dominant isolates, our study identified a minimal set of functions consistently enriched in the isolates that collectively form root communities (Figure 5). We found these functions to be prevalent among plant-associated bacteria and, importantly, they were community-driven functions rather than based on selection of a few elite isolates.

The studies reported here were performed using a sterile, inert substrate, where the only nutrients for bacteria were those provided by the plant host via root exudation and those present in the provided nutrient solution. In these conditions, we found that irrespective of the taxonomic richness present in the initial inoculum, plant roots selected a reduced number of isolates (Figure 2). While acknowledging that increasing the sequencing depth, low abundant or rare colonisers could be captured, these results reflect high competition among bacteria for the available resources. We found that nutrient availability had a significant impact on functions recruited in particular host–inoculum combinations (Figure 2g, Figure S7), but not on the overall observed variation (Figure 3a, S8), indicating that recruitment of basic commensal functions into the root and rhizoplane niches based on root-provided metabolites is robust. Notably, in natural environments, the soil will provide additional resources and properties that support the survival and growth of bacteria in the rhizosphere(*20*). Consequently, we envision that the interactions observed here will be influenced by physico-chemical properties of the soil, nutrient availabilities, pH, and water, as well as the remaining soil-adapted microbes. All these elements can modulate in various ways the survival of isolates with a preferred plant-associated lifestyle(*20*). Nonetheless, based on the well-documented and consistent enrichment of isolates from the soil into root microbiota(*25, 43*) we envision that those functions identified here using nutrient-limited experimental conditions are traits of isolates well-adapted to the root niches that provide an advantage for bacterial enrichment in the rhizoplane and root communities.

The three hosts included in our study were found to have different strategies for assembling their root communities. Arabidopsis and Barley had a lower impact compared to Lotus, which was much more restrictive in terms of how many isolates could be accommodated—but, importantly, those isolates, even if fewer, covered a diverse panel of functions. In other words, Arabidopsis and Barley were more permissive hosts, while Lotus followed a strategy of selecting versatile ‘Swiss army knife’ isolates. Plants secrete up to 20% of their photosynthates into the soil (*14*), which vary according to plant taxonomy, genotype, and physiology(*13, 54, 55*). We envision that the differences we observed here between host species could be driven by exudate profiles and their root surface morphology. Lotus is the only host in our study with nitrogen-fixing symbiotic capacity; thus, we can also envision a possibility whereby the host is genetically wired to support the enrichment of the symbiotic bacteria, and thus the remaining members of the community need to be highly competitive to be Lotus root competent(*56*).

The plant-root microbiota is incredibly complex and, at a low taxonomic level, shows large variability that is host and environment specific. This work revealed that we can identify strategies used by the different plant hosts to structure their root microbiota and minimal sets of bacterial functions, providing soil bacteria with increased root colonisation competency advancing our future strategies for inoculant selection, microbiome modelling, engineering and microbiome-based plant breeding(*7, 8, 30*).

## Acknowledgments and funding

The authors want to thank members of the Plant-Microbe Interactions lab (Utrecht, The Netherlands) and the Molecular Biology and Genetics lab (Aarhus, Denmark) for helpful discussions and to Jens Stougaard, and Corné M.J. Pieterse for critical feedback on the manuscript. This study was supported by the Novo Nordisk Foundation Grant no. NNF19SA0059362 (GS, FL, AG-R, RdJ, SR) and SK was funded by Bill and Melinda Gates Foundation and the UK’s Foreign, Commonwealth and Development Office (FCDO) through the Engineering Nitrogen Symbiosis for Africa project (ENSA; OPP11772165). We acknowledge the Utrecht Sequencing Facility (USEQ) for providing sequencing service and data. USEQ is subsidized by the University Medical Center Utrecht and The Netherlands X-omics Initiative (NWO project 184.034.019).

## Author contributions

Conceptualization: SR, RdJ,

Methodology: SR, RdJ, GS, FL, AG-R, SK

Investigation: GS, FL, AG-R, SK, ZB

Visualization: GS, FL, AG-R

Funding acquisition: SR, RdJ,

Project administration: SR, RdJ,

Supervision: SR, RdJ, SK

Writing – original draft: SR, RdJ, GS, FL,

Writing – review & editing: SR, RdJ, GS, FL, AG-R, SK

