## Supplemental Information for "Functional capacities drive recruitment of bacteria into plant root microbiota"

### Supplemental information - overview

S1 – Experimental photos of inoculations
S2 – Shoot weights
S3 – *Lotus japonicus* nodule counts
S4 – Pseudoalignment of reads to bacterial genomes – inoculum
S5 – Pseudoalignment of reads to bacterial genomes – bacterial taxonomy
S6 – Alpha diversity plot – rearrangement of samples
Supplemental Results (SR)1 – Nutrient condition effect on diversity analyses
S7 – Alpha diversity plot – nutrient condition subset
S8 – Beta diversity plot – distance to centroids
S9 – Beta diversity plot – isolate
S10 – Computational simulation on taxonomy-functional PERMANOVA
S11 – Correlation functional diversity and bacterial root abundance – family
S12 – Correlation functional diversity and bacterial root abundance – genus
S13 – Genome completeness of bacterial isolates
S14 – PERMANOVA $R^2$ on multiple taxonomic ranks – without nodulators or *Rhizobacter* sp. P2_G4
S15 – Piedonut plot on host PERMANOVA $R^2$ – without nodulators or *Rhizobacter* sp. P2_G4
S16 – Heatmap host PERMANOVA $R^2$ – host drop-out effect
S17 – PERMANOVA $R^2$ on multiple taxonomic ranks – host drop-out effect – without nodulators and *Rhizobacter* sp. P2_G4
S18 – PERMANOVA $R^2$ on multiple taxonomic ranks – host drop-out effect – without nodulators
S19 – PERMANOVA $R^2$ on multiple taxonomic ranks – host drop-out effect – without *Rhizobacter* sp. P2_G4
S20 – DESeq2 number of significant KOs – overlap across hosts and inocula
S21 – DESeq2 – fold change of KOs root vs input
S22 – Heatmap on general pathways
SR2 – General host-selected functions – the 266 KO set
S23 – General selected KOs – selection strictness, host-specificity distribution, distribution across isolates

SR3 – Host-specific functions

S24 – Arabidopsis-specific functions – *bch/chl*

S25 – Barley-specific functions – *exo*

S26 – Lotus-specific nodulator functions – *nif*

S27 – Lotus-specific nodulator functions – *nod*

S28 – Lotus-specific functions – *ery*

SR4 – Host-specific functions – ABC transporters

S29 – Ternary plots of most frequent host-specific pathways

S30 – Correlation ABC transporter diversity and bacterial root abundance – isolates

S31 – Heatmap ABC transporter diversity and bacterial root abundance – family

S32 – Correlation ABC transporter diversity and bacterial root abundance – family

### Supplemental Data and Tables - overview

#### Data

S1 – KO profiles of bacterial isolates
S2 – SSC isolate taxonomy
S3 – Overview of SSC samples and sequencing output
S4 – Phenotypic data – shoot weights and *Lotus japonicus* nodule numbers
S5 – SSC isolate genome lengths
S6 – SSC microbiome – isolate total abundance
S7 – SSC microbiome – KO total abundance
S8 – SSC metadata
S9 – KO annotations, pathways, and pathway annotations
S10 – Updated KO annotations, pathways, and pathway annotations
S11 – KO intravariability in all SSC KOs
S12 – CheckM output on SSC genome assemblies
S13 – DESeq2 output – root vs input
S14 – Pathway abundances
S15 – Isolates used in Levy *et al.* (2018)
S16 – KOs that were significantly plant-associated in Levy *et al.* (2018)
S17 – Pathway fold change root vs input
S18 – KO fold change root vs input
S19 – KO fold change root vs input – host subset
S20 – KO fold change root vs input – host subset and non-nodulator dataset
S21 – Gene cassettes
S22 – Host-specificity of ABC transporters
S23 – Shiny app – original dataset
S24 – Shiny app – dataset without dominators

#### Tables

S1 – Physicochemical composition of soils from which the bacterial isolates derive
S2 – Nutrient media used in SSC experiment
S3 – Growth speed and media – SSC isolates
S4 – Abundance distribution of computationally simulated microbiome samples
S5 – Overview of the 852 and 266 KOs
S6 – Overview of the pathway-annotated KOs among 852 and 266 KOs
S7 – Overview of the 852 and 266 KOs in Levy *et al.* (2018)
S8 – Number and percentages of double selected KOs between inocula
S9 – Arabidopsis-specific KOs
S10 – Barley-specific KOs
S11 – Lotus-specific KOs
S12 – Lotus-specific KOs – no nodulator dataset
S13 – Host-specific pathways - frequency

### Glossary

Culture collection – Collections of pure bacterial cultures, most commonly preserved in glycerol stocks.

Family representativeness – An indication on how well the isolate represents the family in terms of functionality, reflected by the family KO proportion: the percentage of unique KOs that a bacterial isolate has from the total amount of unique KOs that can be found in the family (pangenome).

Functional convergence/divergence – the process in which the composition of bacterial functions in the root microbiome between hosts becomes more similar (convergence) or dissimilar (divergence).

Functional diversity – An indication of a bacterial isolate's functional capacity and versatility. A high number of unique functions indicates a high functional diversity

Genome completeness – An overview of the presence and absence of conserved bacterial markers in the genome, indicating how complete the genome is assembled from the short reads.

KOs – functional gene annotations from the Kyoto Encyclopedia of Genes and Genomes (KEGG) database.

KO diversity – The number of unique KOs that an inoculum or bacterial isolate has, which is indicative of the functional capabilities.

KO intravariability/gene diversity – The diversity that exists within a KO. A high KO intravariability indicates the occurrence of multiple genes that have the same KO annotation but diverging nucleotide sequences, which could indicate multiple different functions within the KO. A low KO intravariability might indicate fewer genes in a KO or the presence of (near-)identical gene copies, suggesting a similar functional.

KO/gene branch length – The cumulative branch lengths in a dendogram of all genes within a KO built from nucleotide sequence identity. A high branch length indicates a high KO intravariability and thus a high number of distinct genes within the KO.

Microbial context – an observation in the microbial context, means it can only be observed in this specific microbiome (inoculum) composition and not in others.

PERMANOVA $R^2$ value – The proportion of compositional differences explained by a metadata variable. When investigating the microbiome or functional composition of samples, i.e. within-sample counts of bacterial isolates or bacterial functions, we investigate how similar or distinct samples are by looking at the compositional differences. When samples derive from different hosts or are inoculated with different inocula, for example, we can calculate how much the composition is affected by these variables (the hosts or inocula), by calculating the $R^2$ value for each variable.

SynCom reconstitution experiment – Experiment in which bacterial isolates are individually cultivated and assembled into a (synthetic) community in similar proportions. The SynCom or inoculum members are most commonly representing the natural community. Inoculum and SynCom are synonyms.

### Methods

*Culture collection establishment and genome assembly - AtSC*

The *Arabidopsis thaliana* (hereafter Arabidopsis) culture collection was isolated from Arabidopsis Col-0 plants and *myb72* mutant plants grown in iron-deficient natural soil (Reijerscamp soil, the Netherlands) (Berendsen *et al.*, 2018; Selten *et al.*, 2024; Stringlis *et al.*, 2018) (Table S1). The rhizospheres of the Arabidopsis plants were harvested at three days and seven days post-transplantation to the soil system by removing the roots from the soil and shaking vigorously to only keep tightly root-associated bacterial isolates. The roots were subsequently washed in 10 mM $MgSO_4$, diluted $10^6$ times and plated on 1/10th tryptic soy agar, Reasoner's 2A agar, yeast extract medium, minimal medium with 2 mM coumarin, potato dextrose agar and King's B medium (Table S2). Single colonies were picked after three, five, seven, ten, fourteen and twenty-one days after plating and individually cultivated in liquid tryptic soy broth.

Bacterial isolates that were harvested at three days post-transplantation were subjected to high throughput two-step barcoded amplicon sequencing on the V3-V4 region of the *16S rRNA* gene. In both PCRs, the V3-V4 region of the *16S rRNA* gene was amplified using the degenerate i7 (5′ CAAGCAGAAGACGGCATACGAGAT - [i7] - GTCTCGTGGGCTCGG 3') and i5 primers (5′ AATGATACGGCGACCACCGAGATCTACAC - [i5] - TCGTCGGCAGCGTC 3'). The barcoded V3-V4 *16S rRNA* amplicons were purified after the first and second PCR using Agencourt AMPure XP beads. Finally, the V3-V4 *16S rRNA* amplicons were pooled and sequenced on an Illumina MiSeq platform. The number of bacterial isolates was narrowed down according to uniqueness in the V3-V4 *16S rRNA* sequence using the DADA2 plugin of Qiime2 (Qiime2 version 2019.1) (Bolyen *et al.*, 2019). Additional bacterial isolates that were isolated from plants at seven days post-transplantation were inspected for colony morphological characteristics such as size, color, shape, opacity, elevation, stickiness and shininess, and morphologically novel ones were added to those from three-day soil-grown plants to build up the Arabidopsis culture collection.

Subsequently, DNA was extracted from pure bacterial cultures using the MagAttract Microbial DNA kit and genomic fragments were amplified using the degenerate Nextera barcoded i5 (5' AATGATACGGCGACCACCGAGATCTACAC - [i5] - TCGTCGGCAGCGTC 3') and i7 primers (5' CAAGCAGAAGACGGCATACGAGAT - [i7] - GTCTCGTGGGCTCGG 3') following the Hackflex protocol (Gaio *et al.*, 2022). Amplified genomic fragments were purified using the AMPure XP beads, pooled and sequenced on an Illumina NovaSeq platform. Genomic reads were demultiplexed and cleaned with cutadapt (version 2.8) (Martin, 2011) and assembled into genomes using A5 (A5-miseq version 20160825) (Coil *et al.*, 2015). Genome completeness, contamination, and heterogeneity (collectively, purity) was checked with CheckM (version 1.1.3) (Parks *et al.*, 2015) and any genomes with multiple single copy gene occurrences were subjected to a metagenomic binning procedure (MaxBin version 2.2.7) (Wu *et al.*, 2014) to disentangle genomes from these apparent mixed bacterial cultures. Non-bacterial contigs in the genome assemblies were removed using MMSeqs2 (version 13.45111) (Steinegger & Söding, 2017). Next, dRep (version 3.4.1) (Olm *et al.*, 2017) was run on the 563 genomes to dereplicate the genomes to a non-redundant set of 447 unique genomes at 99.99% sequence identity. Open reading frames were identified and annotated by PROKKA (version 1.14.6) (Seemann, 2014) and EggNOG (version 2.1.4-2) (Cantalapiedra *et al.*, 2021), respectively (Supplemental Data S1).

The isolates' taxonomy was inferred using the Genome Taxonomy DataBase (GTDB, version v2.3.0) (Chaumeil *et al.*, 2020) (Supplemental Data S2).

*Culture collection establishment and genome assembly - HvSC*

The *Hordeum vulgare* (hereafter Barley) culture collection is a cereal culture collection isolated from both Barley (Golden Promise) and *Zea mays* (hereafter Maize) (W22) roots grown in Askov soil (Denmark) (Overgaard *et al.*, 2022) (Table S1). Bacteria associated with Barley and Maize roots were isolated at seven weeks post germination. The roots were harvested and excess soil particles were removed, and the remaining roots were shaken vigorously in sterile water. The bacteria-soil suspension was diluted $2x10^6$ in $1/10^{th}$ tryptic soy broth or in yeast extract medium and plated in 96-well plates (Table S2). Pure bacterial cultures were isolated at seven- and fourteen-days post-harvest.

From these bacterial suspensions, the DNA was extracted and subjected to high-throughput two-step barcoded amplicon sequencing on the V5-V7 region of the *16S rRNA* gene. High throughput DNA extraction was performed by adding a basic buffer to the bacterial suspension (100 mM NaOH and 100 mM EDTA at pH = 12) followed by a thermal lysis of the bacterial cells and the addition of a pH buffer (40 mM Tris-HCl at pH = 7.46). The V5-V7 region of the *16S rRNA* gene was amplified in both PCRs by degenerate 799F (5' AATGATACGGCGACCACCGAGATCTACAC - [P5] - GACTGCGACTGGCG - AACMGGATTAGATACCCKG 3') and 1192R (5' CAAGCAGAAGACGGCATACGAGAT - [P7] - CAGCCATTTAGTGTC - ACGTCATCCCCACCTTCC 3') primers, cleaned by Agencourt AMPure XP beads after the first and second PCR, pooled, and sequenced on an Illumina HiSeq platform. The number of bacterial isolates was narrowed down according to uniqueness in the V5-V7 *16S rRNA* sequence using the USearch plugin of Qiime1 (Qiime1 version 1.9.1) (Caporaso *et al.*, 2010).

Thereafter, the bacterial isolates were recultivated on tryptic soy agar and yeast extract medium plates to isolate pure bacterial colonies and remove any contamination. Next, the DNA was extracted from these liquid cultures for whole-genome sequencing according to Gonzalez-y-Merchand *et al.* (1996). DNA library preparation, whole genome sequencing, genomic read filtering and clean-up, genome assembly, genome assembly quality check, functional annotation and taxonomy inference were performed as described for the Arabidopsis culture collection (Supplemental Data S1 and S2). The final version of the Barley culture collection consists of 392 isolates dereplicated to 364 unique genomes at 99.99% sequence identity.

*Culture collection establishment and genome assembly - LjSC*

The *Lotus japonicus* (hereafter Lotus) culture collection derives from Lotus Gifu plants cultivated in Cologne Agricultural Soil (CAS) (Germany) (Bulgarelli *et al.*, 2012) (Table S1). An extensive description of the isolation, sequencing and genome assembly of the Lotus culture collection can be found in Wippel *et al.* (2021). The Lotus culture collection is publicly available under the accession number PRJEB37696. Additionally, we used MMSeqs2 (version 13.45111) (Steinegger & Söding, 2017) to clean the assemblies from any non-bacterial contigs, similarly to the Arabidopsis and Barley culture collections, and used dRep (version 3.4.1) (Olm *et al.*, 2017) to dereplicate the isolate set of 290 isolates to 177 unique genomes at 99.99% sequence identity. We also re-inferred the taxonomy of the bacterial isolates using GTDB (version v2.3.0) (Chaumeil *et al.*, 2020) (Supplemental Data S2).

*SynCom preparation*

All bacterial isolates from the Arabidopsis (563), Lotus (290) and Barley (392) culture collections were cultivated at nine, seven and five days prior to inoculation in 1/10th tryptic soy broth or yeast extract medium broth depending on their growth speed (Table S2 and S3). Deep-well cultivation plates (2 mL) were filled with 1 mL of bacterial growth media and 10 µL bacterial suspension, sealed with breathable membranes, and incubated on a shaker at 21°C.

On the day of the inoculation, bacterial isolates were washed twice by spinning the deep-well plates at 4,700 rpm for 8 min, removing the supernatant, adding 600 µL of low nutrient Long Ashton medium (SLA) (Table S2) and resuspending the bacterial culture. The bacterial cultures were subjected to OD600 measurement and pooled per host-specific inoculum in similar proportions at an OD600 of 0.1. The pooled AtSC, HvSC and LjSC were subsequently subjected to another OD600 measurement, extraction of a small aliquot to facilitate analysis of the input samples, pooling, and finally dilution to an OD600 of 0.02 using high nutrient SLA, low nutrient SLA or Hoagland. The AtSC, HvSC and LjSC were pooled in equal volume and concentration ratios to create the SSC.

*Seed surface-sterilization and germination*

Arabidopsis seeds were surface sterilized in 70% ethanol for two minutes, washed four times in sterile water, submerged in sterile water and stored in 4 °C in the dark for three days to induce seed imbibition and stratification. Arabidopsis germination was induced by placing the stratified seeds on a Murashige & Skoog agar (MS) plate (Table S2), which was then placed vertically at 20°C with a 16/8 hour light/dark cycle nine days prior to inoculation.

Barley seeds were surface sterilized in 1:20 bleach (NaClO) solution for 15 minutes, followed by five washes with sterile water and an hour incubation in sterile water on a moving tray for seed imbibition. Subsequently, Barley seeds were placed in between sterile filter paper sheets that were soaked with sterile water previously and stored in sealed-off petri dishes horizontally at 20°C with a 16/8 hour light/dark cycle five days prior to inoculation.

Lotus seeds were scarified and surface sterilized by sandpaper scrubbing and submerging them in 1:20 bleach (NaClO) solution for 20 minutes respectively. Next, Lotus seeds were washed with sterile water and imbibed in sterile water at 4°C overnight. Lotus seeds were then placed on sterile filter paper sheets that were soaked with sterile water into sealed-off petri dishes and placed vertically at 20°C with a 16/8 hour light/dark cycle six days prior to inoculation.

*Mycorrhizal germination*

Mycorrhizal spores (Symbiom, MycoGrow™ batch from 2020, Czech Republic) were incubated at a concentration of 500 spores per mL at 28°C in low nutrient SLA media three days before inoculation in a dark environment (Table S2). The SLA was supplemented with strigolactone-like hormone GR24 to a concentration of 100 nM to induce germination. On the day of the inoculation, the fungal spores were washed three times with long nutrient SLA before addition to the experimental system.

*Experimental set-up*

UV-light sterilized black plastic vase-shaped square pots with a height of 15 cm and 7 x 7 cm width and length at the top were filled with filter paper and three different particle layers respectively: a 3-4 cm leca (lightweight expanded clay aggregate) grain layer (particle size between 0.5 and 1 cm), a 10 cm layer containing a mix of leca grains (leca particle size < 0.5 cm), vermiculite, gray sand and sterile water mix layer in a 3:3:1:1 ratio and a 1-2 cm vermiculite layer. These substrate layers were autoclaved for one hour at 120°C before filling the pots.

Two days prior to the inoculation, pots were water-saturated by the addition of 25 mL sterile water, and five Arabidopsis, one Barley or ten Lotus seedlings were transplanted into the pots with their roots facing downward. Each treatment consisted of a different combination of inoculum, host, and nutrient condition and included four replicates/pots (Figure 2a). The following control treatments were used; 1. hosts grown without inoculum and mycorrhizae, 2. substrate-soil control with inoculum and mycorrhizae, 3. substrate-soil control without inoculum but with mycorrhizae and 4. substrate-soil control without inoculum and mycorrhizae. A sample overview can be found in Supplemental Data S3.

*SynCom inoculation*

The inocula were added to the pots by pouring 25 mL of inoculum (OD600 - 0.02) diligently from the top into the pots. Additionally, 1 mL of SLA-washed mycorrhizal spores were added from the top into the pots. Controls without inoculum received 25 mL of sterile nutrient solution.

*Growth and harvest*

Plants were grown in open pods as indicated. The pots were placed individually in boxes for maintaining individual watering. These were, in turn, placed into larger boxes to separate host-inoculum combinations. The large boxes were covered with plastic to maintain humidity in the pots during plant establishment. These plastic covers were removed for Lotus and Barley at one week post inoculation, and were kept on the boxes for Arabidopsis until the harvest.

The experiment was conducted twice, with small alterations in the growth conditions between the two experiments. In the first experiment (R1), the plants were only watered with low nutrient SLA media and grown under three different conditions when inoculated with the SSC or grown under one host-specific condition when inoculated with the host-specific inocula:

1. Arabidopsis when inoculated with any inoculum as well as Barley and Lotus when inoculated with the SSC were cultivated in short-day conditions at elevated temperatures (10 h/14 h light/dark, 75%/75% humidity, 22/18 °C, 200/0 lux).
2. Barley when inoculated with any inoculum as well as Arabidopsis and Lotus when inoculated with the SSC were cultivated in long-day conditions at lower temperatures (16 h/8 h light/dark, 80%/80% humidity, 18/16 °C, 200/0 lux).
3. Lotus when inoculated with any inoculum as well as Arabidopsis and Barley when inoculated with the SSC were cultivated in long-day conditions at elevated temperatures (16 h/8 h light/dark, 80%/80% humidity, 22/18 °C, 200/0 lux).

In the second experiment (R2), the three plant species were grown in only host-specific growth conditions irrespective of inoculum.

1. Arabidopsis plants (day/night) were grown in short-day conditions at elevated temperatures (10 h/14 h light/dark, 75%/75% humidity, 22/18 °C, 200/0 lux).
2. Lotus and Barley plants (day/night) were grown in long-day conditions at elevated temperatures (16 h/8 h light/dark, 75%/75% humidity, 22/18 °C, 200/0 lux).

Plants were regularly bottom-watered with low nutrient SLA (R1 and R2), high nutrient SLA (R2) or Hoagland (R2 – only Arabidopsis), depending on the nutrient condition that these plants were inoculated with. Three weeks post-inoculation, plants were removed from the pots, shoots were cut off from the roots and the roots were isolated from the middle clay, vermiculite, and gray sand layer using sterile tweezers. The shoots were weighted (Supplemental Data S4) and representative pictures of each host and treatment combination (inoculum and nutrient condition) were taken (before removal from the pots) (Figure S1). Excess soil was removed from the roots and roots were washed five times in sterile water by vigorous shaking. The wash-off of the first wash was spun down for 15 minutes at 4,700 rpm, excess water was removed, and the pellet's resuspension was taken as the rhizosphere fraction. Finally, the firmly washed roots were dried on filter paper and, together with the rhizosphere fraction, stored at -80 °C. For Lotus, nodules were separated from the roots, counted, and separately stored at -80°C (Supplemental Data S4). The bulk soil samples consisted of 300 mg of substrate taken from the middle clay grain, vermiculite, gray sand layer and stored at -80°C.

*DNA library preparation and community profiling*

DNA was extracted from input, nodule, rhizosphere, root (endosphere and rhizoplane), and soil samples using the FastDNA SPIN Kit for Soil according to manufacturer's instructions. Metagenomic DNA was diluted to 5 ng/µL and amplified using the Hackflex protocol (Gaio *et al.*, 2022). Amplified metagenomic fragments were purified using the AMPure XP beads, pooled and sequenced on an Illumina NovaSeq 6000 platform (PE150, 2x150bp).

An overview of the sequencing data is presented in Supplemental Data S3. Metagenomic samples were cleaned from host reads, depending on the host used in each case, using Bowtie2 (version 2.2.5) (Langmead & Salzberg, 2012). Once the samples were cleaned, the metagenomic reads were pseudoaligned against the relevant reference index composed of the genome sequences of the inoculated strains using Salmon (version 1.9.0), except for analysis of within- and between-sample diversity in which case we used the same reference index for all samples based on the genome sequences of all SSC isolates (Patro *et al.*, 2017). Default settings were used for Salmon except for a minimum score for pseudoaligning reads (--minScoreFraction) set at 0.95, allowing for the distinction of bacterial isolates with high genomic similarity. The Salmon output was assembled into a microbiome count table and normalized for the bacterial isolates' genome lengths (Supplemental Data S5 and S6). An overview of the proportion of pseudoaligned reads to the plant genome and SynCom indices is shown in Supplemental Data S6. The PICRUSt2 (version 2.5.1) algorithm was used to extrapolate the bacterial abundances to KO abundances to investigate the functional or KO composition of the samples (Douglas *et al.*, 2020) (Supplemental Data S7). The KO profiles from Supplemental Data S1 were used for this extrapolation step. For further analysis, the isolate and KO abundances were normalized to produce relative abundances. The metadata can be found in Supplemental Data S8.

*Data analysis – phylogenetic placement*

Phylogenetic distances between bacterial isolates were calculated with Orthofinder using STAG support values and STRIDE was used to root the phylogenetic tree of the isolates (Figure 1a) (Emms & Kelly, 2019).

*Data analysis – KO intravariability*

KO diversity, as presented in Figure 1c, was assessed by establishing a kmer-based phylogenetic tree of each KO in each inoculum using the gene sequences as nodes with the ape package in R (version 5.7-1) (Paradis & Schliep, 2019). The sum of all edge lengths is used to obtain the total branch length or total gene diversity (Figure 1c; x-axis) (Supplemental Data S11). The number of nodes in the phylogenetic tree represents the total number of genes (Figure 1c; y-axis).

*Data analysis – phenotypic data*

Phenotypic data such as shoot weight and the number of nodules were statistically assessed by a Kruskal Wallis test followed by a Dunn post hoc test (Figures S2 and S3).

*Data analysis – diversity analyses*

The alpha diversity analyses (Figures S6 and S7) were performed using for each samples the isolates with relative abundance values >0.05% or KOs with relative abundance >1.5e-5% (0.05% divided by the average number of KOs per genome (3,295)). To calculate the observed isolates, observed KOs, and the Shannon diversity of the isolate and KO compositions we used the phyloseq package in R (version 1.44.0) (McMurdie & Holmes, 2013). Statistical differences between the inocula and/or hosts were assessed by an ANOVA with post-hoc Tukey HSD test (Figures 2b, c, d, e, S6, and S7). Beta diversity analyses to calculate the Bray-Curtis distances between samples (Figures 2f, g, S8, and S9) were conducted on the original microbiome and functional abundance tables using the vegan package in R (version 2.6-4, Oksanen *et al.*, 2013). Statistical significance of compositional differences by different variables was inferred by an Adonis test/PERMANOVA from the vegan package in R. The $R^2$ of the Adonis test was calculated to investigate what proportion of the total compositional variance is explained by different variables such as inoculum, host, or nutrient condition, calculated on multiple taxonomic levels and the functional level in Figure 3a and S14. In these figures, the nitrogen-fixing symbionts of Lotus or *Rhizobacter* sp. P2_G4, representing the dominant isolates in the dataset, were computationally excluded to quantify their effect on the microbiome composition. To investigate variation within samples of the same treatment (unique combination of host, inoculum, and nutrient condition), we determined the centroid of the treatment in the PCoA plane and calculated the distance from each sample to the centroid (Figure S8).

*Data analysis – simulation of within and between community diversity*

To investigate how Adonis/PERMANOVA $R^2$ values of the microbiome composition correlate to the $R^2$ values of the functional composition, we conducted a simulation with 1,000 iterations (Figure S10). In each iteration, we generated a randomized microbiome dataset consisting of 54 samples (equal to those present in our dataset): nine groups of six replicates. We determined that the average number of isolates that are recruited across the root samples was 119, and thus selected this number of random isolates for every host-specific inoculum thrice, thus totaling nine groups. The random abundance values resembled the abundance distribution in the actual SSC dataset (Table S4). The microbiome dataset was then extrapolated to a functional dataset using the PICRUSt2 (version 2.5.1) algorithm (Douglas *et al.*, 2020) and the Bray-Curtis distances between the samples on both the genus and KO composition were calculates as before. We categorized the dataset in three, six, or nine groups by K-means clustering before calculating the Adonis $R^2$ values that separate these three, six, or nine groups in both the isolate and functional dataset. The isolate and functional $R^2$ values were subsequently plotted to each other, and a Pearson correlation was used to calculate the significance and slope of the correlation.

*Data analysis – family representativeness*

The family and genus representativeness of isolates (Figures 3d, 3e, S11, and S12) was assessed by calculating the representativeness in KO composition (Formula A) and the representativeness in abundances in the root communities (Formula B). Family representativeness was calculated for the host-specific inocula that they derive from (AtSC, HvSC and LjSC). Genome completeness as assessed previously is added to the family representativeness in Figure S13 (Supplemental Data S12), demonstrating that poor assembly quality does not lead to a lower KO diversity.

$$A)\ KO\ proportion\ = \frac{Number\ of\ unique\ KOs\ in\ isolate}{Number\ of\ unique\ KOs\ in\ family\ pangenome}$$

$$B)\ Relative\ abundance\ proportion\ = \frac{Relative\ abundance\ isolate\ in\ dataset}{Relative\ abundance\ family\ in\ dataset}$$

Figure S31 is visualized in a similar manner, in which the KO diversity is subsetted to ABC transporter KO diversity.

*Data analysis – host effect on host $R^2$*

To investigate the effect of each host on the microbiome and functional composition, we subsetted the data per inoculum and excluded one host at a time to recalculate the $R^2$ value and compare it with that obtained $R^2$ value for the entire data set. An overview on how the $R^2$ value changes at each taxonomic level and functional level in the different datasets (with and without dominators or without nodulators or *Rhizobacter* sp. P2_G4) can be found in Figures S17, S18, and S19. The summary of this analysis on the functional composition can be found in Figure S16.

The values presented in Figure S16 were used to calculate the contribution of each host to the host $R^2$ presented in Figures 3b, c, and S15. For each host, the difference between the host $R^2$ value with all three hosts and the host $R^2$ value of the dataset with the other two hosts is calculated. The resulting value divided by the sum of all these values leads to the host contribution (Formula C) (outer ring in Figures 3c, 3d, and S15). To investigate the degree to which the hosts affect the host $R^2$ in each inoculum, the contributions of the three hosts in each inoculum are summed and divided by the summed contributions of the three hosts across all inocula (Formula D) (inner ring in Figures 3c, 3d, and S15).

$$C1)\ Arabidopsis\ contribution = 100\% * \frac{Host\ R^2 - Host\ R^2_{Barley\ \cap Lotus}}{\sum Host\ R^2 - Host\ R^2_{Arabidopsis\ \cap Barley} + Host\ R^2 - Host\ R^2_{Arabidopsis\ \cap Lotus} + Host\ R^2 - Host\ R^2_{Barley\ \cap Lotus}}$$

$$C2)\ Barley\ contribution = 100\% * \frac{Host\ R^2 - Host\ R^2_{Arabidopsis\ \cap Lotus}}{\sum Host\ R^2 - Host\ R^2_{Arabidopsis\ \cap Barley} + Host\ R^2 - Host\ R^2_{Arabidopsis\ \cap Lotus} + Host\ R^2 - Host\ R^2_{Barley\ \cap Lotus}}$$

$$C3)\ Lotus\ contribution = 100\% * \frac{Host\ R^2 - Host\ R^2_{Arabidopsis\ \cap Barley}}{\sum Host\ R^2 - Host\ R^2_{Arabidopsis\ \cap Barley} + Host\ R^2 - Host\ R^2_{Arabidopsis\ \cap Lotus} + Host\ R^2 - Host\ R^2_{Barley\ \cap Lotus}}$$

$$D1)\ Host\ effect\ in\ AtSC\ = 100\% * \frac{\sum Arabidopsis\ contribution\ +\ Barley\ contribution +\ Lotus\ contribution\ in\ AtSC}{\sum AtSC(Arabidopsis\ contribution +\ Barley\ contribution +\ Lotus\ contribution\ in\ AtSC) + \sum HvSC + \sum LjSC + \sum SSC}$$

$$D2)\ Host\ effect\ in\ HvSC\ = 100\% * \frac{\sum Arabidopsis\ contribution\ +\ Barley\ contribution +\ Lotus\ contribution\ in\ HvSC}{\sum AtSC(Arabidopsis\ contribution +\ Barley\ contribution +\ Lotus\ contribution\ in\ AtSC) + \sum HvSC + \sum LjSC + \sum SSC}$$

$$D3)\ Host\ effect\ in\ LjSC\ = 100\% * \frac{\sum Arabidopsis\ contribution\ +\ Barley\ contribution +\ Lotus\ contribution\ in\ LjSC}{\sum AtSC(Arabidopsis\ contribution +\ Barley\ contribution +\ Lotus\ contribution\ in\ AtSC) + \sum HvSC + \sum LjSC + \sum SSC}$$

$$D4)\ Host\ effect\ in\ SSC\ = 100\% * \frac{\sum Arabidopsis\ contribution\ +\ Barley\ contribution +\ Lotus\ contribution\ in\ SSC}{\sum AtSC(Arabidopsis\ contribution +\ Barley\ contribution +\ Lotus\ contribution\ in\ AtSC) + \sum HvSC + \sum LjSC + \sum SSC}$$

*Data analysis – differential abundance analysis*

Differential abundance testing was using DESeq2 (version 1.40.0) (Love *et al.*, 2014) (Supplemental Data S13). DESeq2 has an internal normalization step that normalizes KO abundances by the geometric mean of the KO, followed by a local regression and a log2foldchange shrinkage using the apeglm shrinkage estimator. Significance was tested by a Wald test and p-values were adjusted for multiple testing using the Benjamini and Hochberg method (significance $p < 0.05$). DESeq2 was run to find the enriched KOs in the root microbiome as compared to the initial inoculum and compare this between hosts and inocula (Figures 4a, b, S20, S23a, b, and Table S8).

To create the heatmap presented in Figure S22, significant KOs by DESeq2 analysis were annotated into pathways and categories that were derived from the KEGG database (Supplemental Data S9). KOs without pathway annotation (3443 KOs) were given an annotation based on their KO annotation (Supplemental Data S9), e.g. transporter, transcriptional regulator, or secretion, to generate a new KO pathway annotation table (Supplemental Data S10).

The relative abundances of all KOs in a pathway that had at least one significant KO were collapsed into pathway relative abundances for every host-inoculum combination. These pathway abundances were averaged across replicates of the same host-inoculum combination and filtered to only include pathways with a higher average relative abundance in the root samples as compared to respective inocula. The results of this analysis were scaled by row and plotted in a heatmap using tidyheatmap in R (version 0.1.0) (Supplemental Data S14).

*Data analysis – differential abundance analysis – general KOs/pathway subsets*

To produce Figures 4d, e, f, g, and h, S21a, and S23d, e, f, g, and h, general KOs were selected from the DESeq2 output (Table S5). In the main figure the threshold was lenient, only allowing KOs that were significant in more than six out of the 12 host-inoculum combinations (852 KOs – Figures 4c, d, g, h, and S21a). For Figures 4e and f, only the KOs were plotted with pathway annotations (641 out of the 852 KOs were assigned to pathways) (Table S6).

In the Supplemental Figures, we used a stricter threshold by only allowing KOs that were significant in all 12 host-inoculum combinations after removing KOs of the Lotus symbionts and *Rhizobacter* sp. P2_G4 (266 KOs – Figures S21b, S23d, g, and h) (Table S5). For Figures S23e and f, only the KOs with pathway annotations from these 266 KOs were plotted (203 out of the 266 KOs were assigned to pathways) (Table S6).

*Data analysis – differential abundance analysis – fold change calculation*

For KOs and pathways, the fold change of isolates with the KO or pathway was calculated between the root samples and the input samples in Figures 4e, f, S23e, f, S24, S25, S26, S27, S28, and S29. For each host-inoculum combination, the cumulative relative abundances of isolates with the KO or pathway were averaged across sample replicates and divided by the average cumulative relative abundances of isolates with the KO or pathway in the respective input inocula (Formula E1) (Supplemental Data S17 and S18). For pathways, only significant KOs were taken into consideration for the calculation of the fold change. In this analysis, inocula for which the KO/pathway was not significant for any host were excluded. The KO/pathway average was then calculated for the remaining inocula in which it is significant.

$$E1)\ FC\ present\ \frac{\frac{\sum_{root\ sample\ replicates}(\sum_{isolates\ with\ KO\ |\ pathway} relative\ abundance\ in\ root)}{number\ of\ root\ sample\ replicates}}{\frac{\sum_{input\ sample\ replicates}(\sum_{isolates\ with\ KO\ |\ pathway} relative\ abundance\ in\ initial\ inoculum)}{number\ of\ initial\ inoculum\ sample\ replicates}}$$

The median fold change across inocula was taken to produce Figures 4d and S23d (Table S6). The respective fold change for KOs and pathways across all host-inoculum combinations were averaged to produce Figures 4e, f, S21, S23e and f.

In Figures 4e, f, S21, S23e, and f, the fold change differences on the y-axes are calculated between the isolates with the KO or pathway as compared to the isolates without the KO (Formula E2 and E3).

$$E2)\ FC\ Absent\ \frac{\frac{\sum_{root\ sample\ replicates}(\sum_{isolates\ without\ KO\ |\ pathway} relative\ abundance\ in\ root)}{number\ of\ root\ sample\ replicates}}{\frac{\sum_{input\ sample\ replicates}(\sum_{isolates\ without\ KO\ |\ pathw} relative\ abundance\ in\ initial\ inoculum)}{number\ of\ initial\ inoculum\ sample\ replicates}}$$

$$E3)\ \text{Enrichment} = \log2(\frac{\text{FC present}}{\text{FC absent}})$$

For the pathway figures (Figure 4e, f, S23e, and f), a Wilcoxon test was conducted to compare the fold changes of the isolates with and without the pathway and only significant differences (Bonferroni p-value adjustment) were kept for Figures 4e, f, S23e, and S23f.

*Data analysis – comparison to Levy et al. (2018)*

To investigate the universality of the KOs identified in this study for root competence, we investigated the distribution of the 852 and 266 KOs from the different selections across the SSC isolates (Figures 4g and S23g). In addition, we also investigated whether the 852 KOs and 266 KOs are plant-associated in the data of Levy *et al.* (2018). The KO profiles of the 3,837 bacteria investigated by Levy et al. (2018) were downloaded (from http://labs.bio.unc.edu/Dangl/Resources/gfobap_website/index.html), and the presence of the 852 or 266 KOs in the KO profiles of all groups of bacteria (non plant-associated, plant-associated, and soil bacteria) was recorded (Figures 4h and S23h) (Supplemental Data S15). In addition, a Venn diagram in Figure 4h and S23h shows the overlaps for the 852 or 266 KOs with the plant-associated KOs found by Levy *et al.* (2018) (Supplemental Data S16 and Table S7).

*Data analysis – Differential abundance analysis – Host-specific KOs/pathways*

To select host-specific KOs, the DESeq2 output was filtered to only include KOs that were significant in the presence of at least three inocula for one host in the original dataset. Similar to the general KOs, the fold change between the relative abundance of isolates with the KO in the root samples and the input samples was calculated for each host individually and the median value from the four inocula was taken (Formula E1) (Supplemental Data S19). From this dataset, KOs were considered host-specific when the fold change for one host was higher than three (>3) and lower than three (<3) for the other two hosts (Tables S9, S10, and S11). These host-specific sets were used to produce Figures 5b, c, and d. In consideration that many of these KOs could be derived from genomes of the nitrogen-fixing symbionts, we repeated the same analysis for Lotus with the computational exclusion of these from the dataset (Supplemental Data S20 and Table S12).

From the host-specific lists of KOs, we selected Arabidopsis, Barley, and Lotus-specific gene clusters for an in-depth exploration (Figures S24, S25, S26, S27, S28 and Supplementary results SR3). The gene abundances, isolate abundances, isolate fold changes between root and input and the presence of these genes across the bacterial phylogeny of these gene clusters is shown in these figures, with an explanation of the pathway in which they function.

To investigate the host-specificity of genes in the most frequent host-specific pathways (Table S13), we collapsed genes into gene cassettes by averaging their fold change for each host and create a non-redundant dataset (Supplemental Data S21). This dataset was used to visualize the isolate fold change per host and inoculum in Figure S29. The most frequent host-specific pathway, ABC transporters, was investigated in-depth with regard to the class of compounds they are predicted to transport (Figure 5d and Supplemental Data S22), and their distribution and diversity across the SSC isolates (Figures S30, S31, and S32). For Figure 5d, the p-values were averaged across gene cassettes that together make up the ABC transporter (Supplemental Data S21) using Stouffer's method.

Aside from the in-depth analysis that was conducted in this manuscript, we generated datasets that can produce Figures 4, e, f, S21, and the individual plots from Figures S24, S25, S26, S27, and S28, meaning the isolate abundances, gene abundances, ternary plots, and the family piedonut plots, for any KO or pathway (pathways only 4e and f) of interest.

*Data availability*

Data deposition: The data reported in this article have been deposited in the National Center for Biotechnology Information Short Read Archive BioProject database. The relevant accession numbers can be found in the separate, specific sections below.

*Arabidopsis culture collection*

Genomic reads from the bacterial cultures are deposited under accession number PRJNA1131834, while the genomic sequences are deposited under accession numbers PRJNA1138681 (isolated at three days post transplantation) and PRJNA1139421 (isolated at seven days post transplantation). Genomes, PROKKA gene predictions, and EggNOG annotations are also available at https://zenodo.org/records/10992416 (Selten *et al.*, 2024).

*Barley culture collection*

Genomic reads from the bacterial cultures are deposited under accession number PRJNA1131819, while the genomic sequences are deposited under accession number PRJNA1139693.

*Lotus culture collection*

Genomic reads as well as genome assemblies from the bacterial cultures have been deposited previously from Wippel *et al.* (2021) under accession number PRJEB37696. Genomic sequences can also be found at https://www.at-sphere.com/. Bacterial isolates LjRoot204, LjRoot204_2, LjRoot205, LjRoot206, and LjRoot208 were not included under this accession number and can be found under accession numbers PRJNA1142758 (genomic reads) and PRJNA1142862 (genomes).

*Metagenome sequencing data*

Shotgun metagenome-sequenced reads from the root microbiomes of Arabidopsis, Barley, and Lotus generated in this study have been deposited under accession number PRJNA1131994.

*Shiny app*

To assist the community in investigating specific KOs in our data, we have built a shiny app under the following link: https://pm-bacterial-genetics-au.shinyapps.io/SSC_community_app/. This app will allow users to investigate KO abundances across hosts and inocula, abundances of isolates with a specific KO/pathway across hosts and inocula, and the functional enrichment of KOs for each host that is similarly calculated in this manuscript. The user can specify the original dataset as well as the dataset in which the dominator isolates, the Lotus symbionts and the HvSC-dominant *Rhizobacter* isolate P2_G4, are computationally excluded. The data to generate the figures in the shiny app are included in Supplemental Data S23 (original dataset) and S24 (exclusion of nodulators and *Rhizobacter* sp. P2_G4 dataset).

*Scripts*

Rscripts developed for the analyses conducted in this study and to generate the figures can be found under the following Zenodo link: https://doi.org/10.5281/zenodo.13322963.

*Methods - references*

Berendsen, R. L., Vismans, G., Yu, K., Song, Y., de Jonge, R., Burgman, W. P., Burmølle, M., Herschend, J., Bakker, P. A. H. M., & Pieterse, C. M. J. (2018). Disease-induced assemblage of a plant-beneficial bacterial consortium. *The ISME Journal*, *12*(6), 1496–1507.

Bolyen, E., Rideout, J. R., Dillon, M. R., Bokulich, N. A., Abnet, C. C., Al-Ghalith, G. A., Alexander, H., Alm, E. J., Arumugam, M., Asnicar, F., Bai, Y., Bisanz, J. E., Bittinger, K., Brejnrod, A., Brislawn, C. J., Brown, C. T., Callahan, B. J., Caraballo-Rodríguez, A. M., Chase, J., … Caporaso, J. G. (2019). Reproducible, interactive, scalable and extensible microbiome data science using QIIME 2. *Nature Biotechnology*, *37*(8), 852–857.

Bulgarelli, D., Rott, M., Schlaeppi, K., Ver Loren van Themaat, E., Ahmadinejad, N., Assenza, F., Rauf, P., Huettel, B., Reinhardt, R., Schmelzer, E., Peplies, J., Gloeckner, F. O., Amann, R., Eickhorst, T., & Schulze-Lefert, P. (2012). Revealing structure and assembly cues for *Arabidopsis* root-inhabiting bacterial microbiota. *Nature*, *488*(7409), 91–95.

Cantalapiedra, C. P., Hernández-Plaza, A., Letunic, I., Bork, P., & Huerta-Cepas, J. (2021). eggNOG-mapper v2: functional annotation, orthology assignments, and domain prediction at the metagenomic scale. *Molecular Biology and Evolution*, *38*(12), 5825–5829.

Caporaso, J. G., Kuczynski, J., Stombaugh, J., Bittinger, K., Bushman, F. D., Costello, E. K., Fierer, N., Peña, A. G., Goodrich, J. K., Gordon, J. I., Huttley, G. A., Kelley, S. T., Knights, D., Koenig, J. E., Ley, R. E., Lozupone, C. A., McDonald, D., Muegge, B. D., Pirrung, M., … Knight, R. (2010). QIIME allows analysis of high-throughput community sequencing data. *Nature Methods*, *7*(5), 335–336.

Chaumeil, P.-A., Mussig, A. J., Hugenholtz, P., & Parks, D. H. (2020). GTDB-Tk: a toolkit to classify genomes with the Genome Taxonomy Database. *Bioinformatics*, *36*(6), 1925–1927.

Coil, D., Jospin, G., & Darling, A. E. (2015). A5-miseq: an updated pipeline to assemble microbial genomes from Illumina MiSeq data. *Bioinformatics*, *31*(4), 587–589.

Douglas, G. M., Maffei, V. J., Zaneveld, J. R., Yurgel, S. N., Brown, J. R., Taylor, C. M., Huttenhower, C., & Langille, M. G. I. (2020). PICRUSt2 for prediction of metagenome functions. *Nature Biotechnology*, *38*(6), 685–688.

Emms, D. M., & Kelly, S. (2019). OrthoFinder: phylogenetic orthology inference for comparative genomics. *Genome Biology*, *20*(1), 238.

Gaio, D., Anantanawat, K., To, J., Liu, M., Monahan, L., & Darling, A. E. (2022). Hackflex: low-cost, high-throughput, Illumina Nextera Flex library construction. *Microbial Genomics*, *8*(1).

Gonzalez-y-Merchand, J. A., Estrada-Garcia, I., Colston, M. J., & Cox, R. A. (1996). A novel method for the isolation of mycobacterial DNA. *FEMS Microbiology Letters*, *135*(1), 71–77.

Langmead, B., & Salzberg, S. L. (2012). Fast gapped-read alignment with Bowtie 2. *Nature Methods*, *9*(4), 357–359.

Levy, A., Salas Gonzalez, I., Mittelviefhaus, M., Clingenpeel, S., Herrera Paredes, S., Miao, J., Wang, K., Devescovi, G., Stillman, K., Monteiro, F., Rangel Alvarez, B., Lundberg, D. S., Lu, T.-Y., Lebeis, S., Jin, Z., McDonald, M., Klein, A. P., Feltcher, M. E., Rio, T. G., … Dangl, J. L. (2018). Genomic features of bacterial adaptation to plants. *Nature Genetics*, *50*(1), 138–150.

Love, M. I., Huber, W., & Anders, S. (2014). Moderated estimation of fold change and dispersion for RNA-seq data with DESeq2. *Genome Biology*, *15*(12), 550.

Martin, M. (2011). Cutadapt removes adapter sequences from high-throughput sequencing reads. *EMBnet Journal*, *17*(1), 10.

McMurdie, P. J., & Holmes, S. (2013). phyloseq: an R package for reproducible interactive analysis and graphics of microbiome census data. *PLOS ONE*, *8*(4), e61217.

Oksanen, J., Blanchet, F. G., Kindt, R., Legendre, P., Minchin, P. R., O'hara, R. B., Simpson, G. L., Solymos, P., Stevens, M. H. H., Wagner, H., & others. (2013). Package 'vegan.' *Community Ecology Package*, *2*(9), 1–295.

Olm, M. R., Brown, C. T., Brooks, B., & Banfield, J. F. (2017). dRep: a tool for fast and accurate genomic comparisons that enables improved genome recovery from metagenomes through de-replication. *The ISME Journal*, *11*(12), 2864–2868.

Overgaard, C. K., Tao, K., Zhang, S., Christensen, B. T., Blahovska, Z., Radutoiu, S., Kelly, S., & Dueholm, M. K. D. (2022). Application of ecosystem-specific reference databases for increased taxonomic resolution in soil microbial profiling. *Frontiers in Microbiology*, *13*.

Paradis, E., & Schliep, K. (2019). ape 5.0: an environment for modern phylogenetics and evolutionary analyses in R. *Bioinformatics*, *35*(3), 526–528.

Parks, D. H., Imelfort, M., Skennerton, C. T., Hugenholtz, P., & Tyson, G. W. (2015). CheckM: assessing the quality of microbial genomes recovered from isolates, single cells, and metagenomes. *Genome Research*, *25*(7), 1043–1055.

Patro, R., Duggal, G., Love, M. I., Irizarry, R. A., & Kingsford, C. (2017). Salmon provides fast and bias-aware quantification of transcript expression. *Nature Methods*, *14*(4), 417–419.

Seemann, T. (2014). Prokka: rapid prokaryotic genome annotation. *Bioinformatics*, *30*(14), 2068–2069.

Selten G., Stassen M.J.J., De Rooij P., Berendsen R.L., Stringlis I. & de Jonge R. (2024). Draft genome sequences of *Arabidopsis thaliana*-associated micro-organisms from Reijerscamp soil, the Netherlands [Data set].

Steinegger, M., & Söding, J. (2017). MMseqs2 enables sensitive protein sequence searching for the analysis of massive data sets. *Nature Biotechnology*, *35*(11), 1026–1028. https://doi.org/10.1038/nbt.3988

Stringlis, I. A., Yu, K., Feussner, K., de Jonge, R., Van Bentum, S., Van Verk, M. C., Berendsen, R. L., Bakker, P. A. H. M., Feussner, I., & Pieterse, C. M. J. (2018). MYB72-dependent coumarin exudation shapes root microbiome assembly to promote plant health. *Proceedings of the National Academy of Sciences*, *115*(22).

Wippel, K., Tao, K., Niu, Y., Zgadzaj, R., Kiel, N., Guan, R., Dahms, E., Zhang, P., Jensen, D. B., Logemann, E., Radutoiu, S., Schulze-Lefert, P., & Garrido-Oter, R. (2021). Host preference and invasiveness of commensal bacteria in the *Lotus* and *Arabidopsis* root microbiota. *Nature Microbiology*, *6*(9), 1150–1162.

Wu, Y.-W., Tang, Y.-H., Tringe, S. G., Simmons, B. A., & Singer, S. W. (2014). MaxBin: an automated binning method to recover individual genomes from metagenomes using an expectation-maximization algorithm. *Microbiome*, *2*(1), 26.

### Supplemental Figures and Results

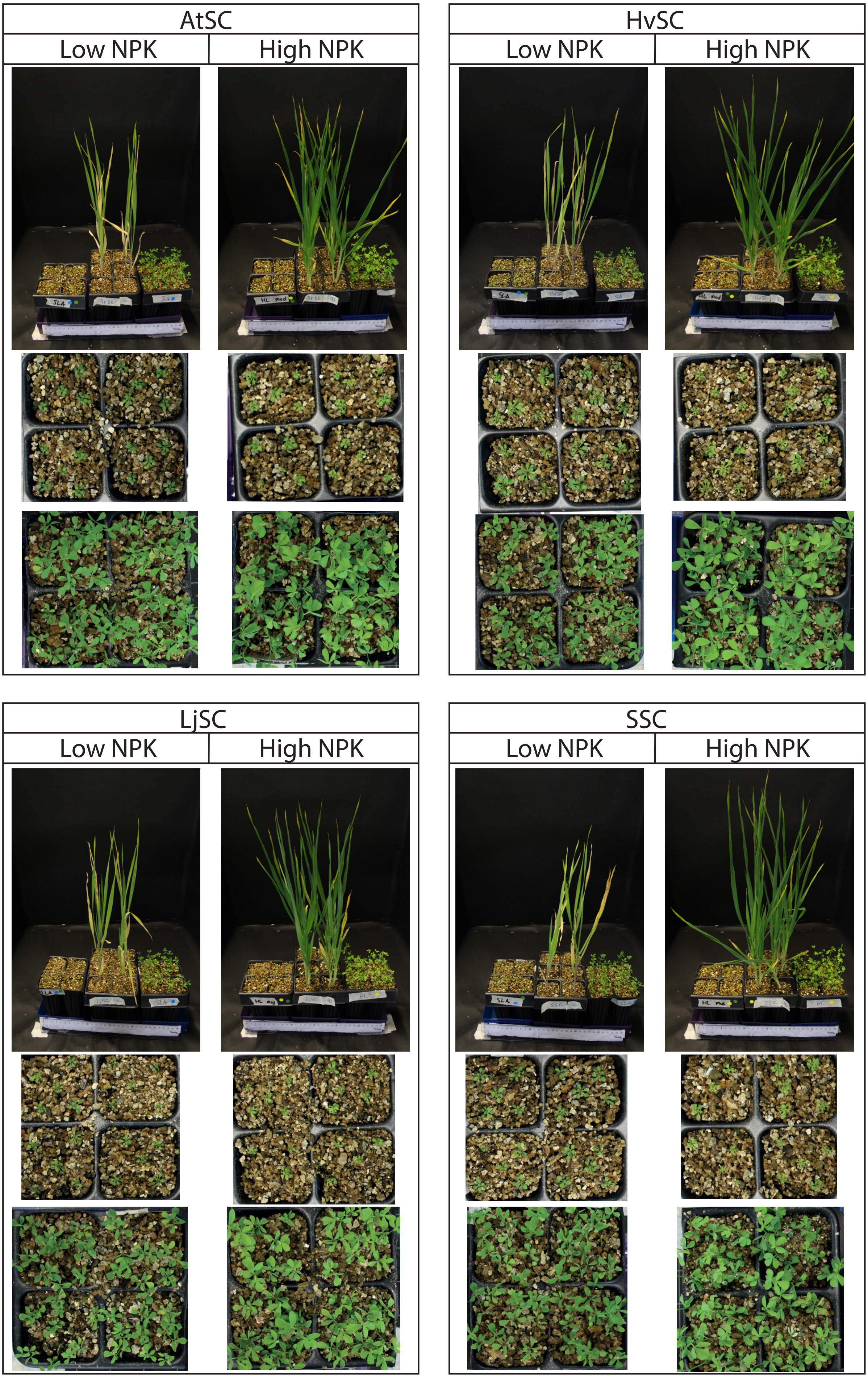

**Figure S1. Experimental photos Arabidopsis, Barley and Lotus shoots.** Phenotypes Arabidopsis, Barley, and Lotus three weeks after inoculation with microbial communities and grown either in low or high nutrient conditions (NPK). All the plants were photographed at the same scale. A close-up top view is added for Arabidopsis and Lotus plants.

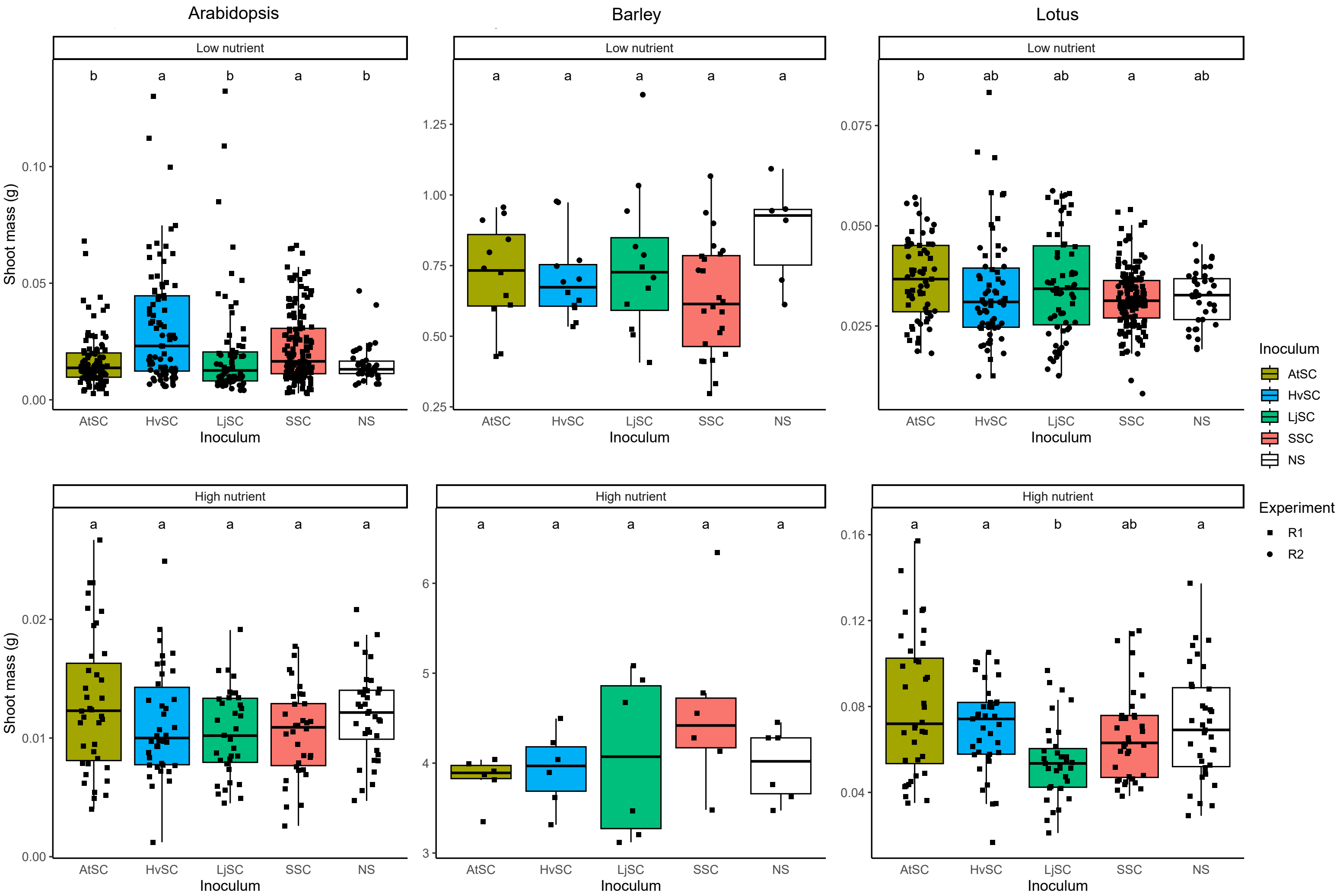

**Figure S2. Fresh shoot biomass of Arabidopsis, Barley and Lotus following microbial community inoculations.** Fresh shoot biomass for Arabidopsis (left), Barley (middle) and Lotus (right) plants when grown under low nutrient conditions (top) and high nutrient conditions (bottom) in the presence or absence (NS) of host-derived inocula (AtSC, HvSC and LjSC) or a combined SuperSynCom (SSC) inoculum. Shapes indicate whether the shoots were from the first (R1) or second (R2) SuperSynCom experiment. Kruskal-Wallis followed by a Dunn's post-hoc test was conducted to investigate statistical significance between inocula subsetted by host and nutrient condition.

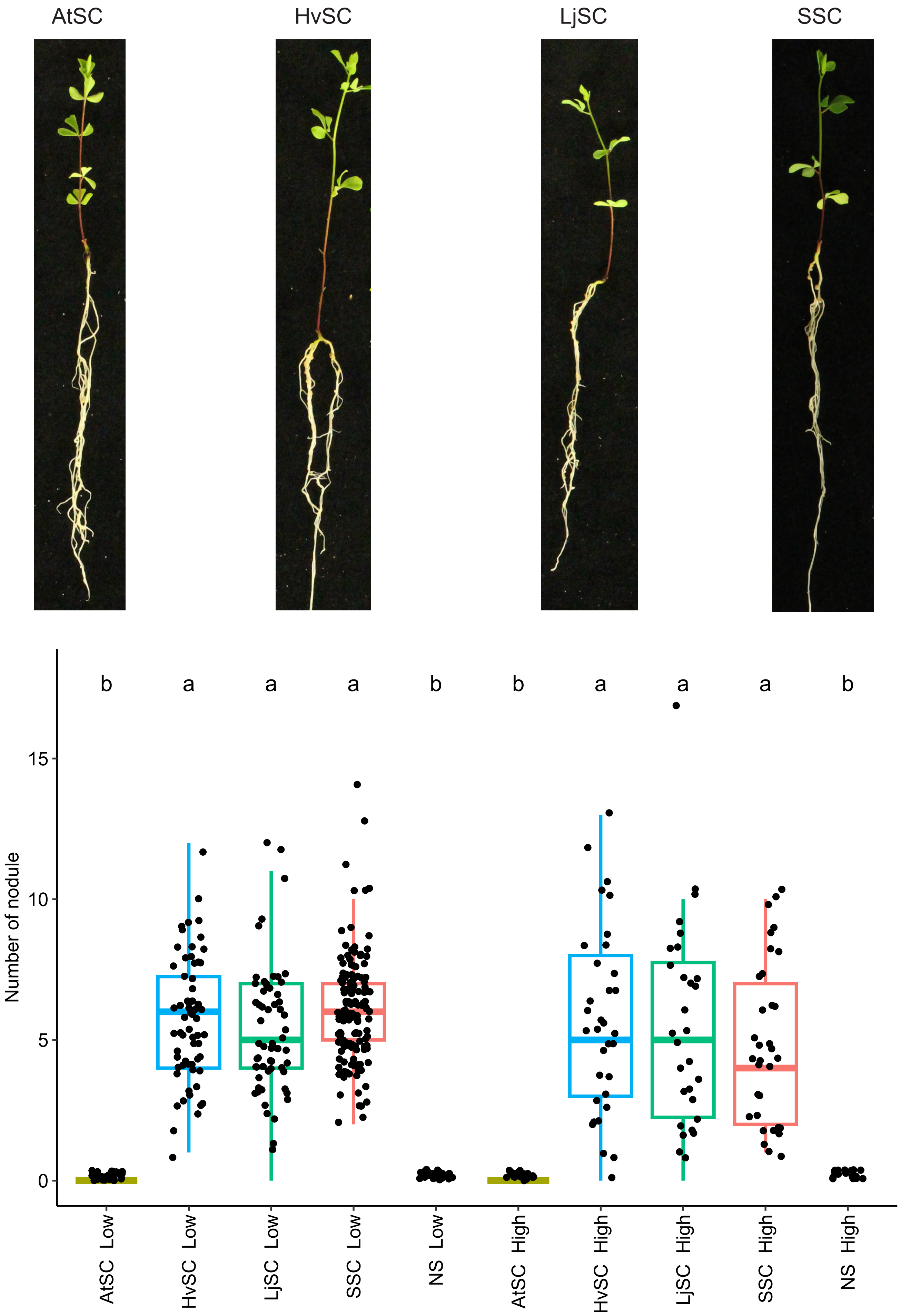

**Figure S3. Number of nodules on Lotus roots following microbial community inoculations.** Representative pictures of Lotus plants inoculated with the four inocula under low nutrient conditions (top) and nodule number of Lotus when inoculated with the four inocula under low and high nutrient conditions (bottom). A Kruskal wallis test followed by a Dunn post hoc test was conducted to analyze statistical significance between groups.

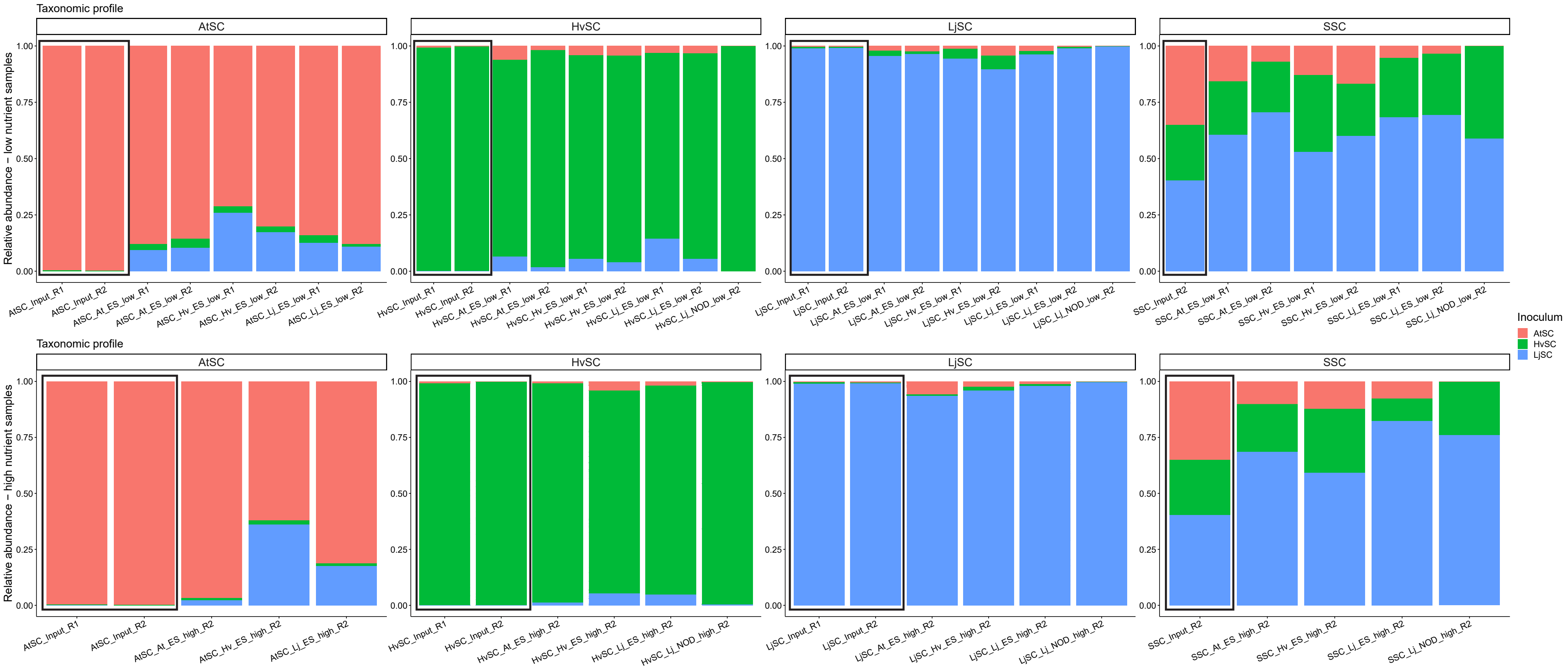

**Figure S4. Read proportion correctly pseudoaligning inoculum index.** Taxonomic profile of the input inocula (bars in black boxes) and the Arabidopsis, Barley and Lotus root microbiome when inoculated under low nutrient conditions (top) and high nutrient conditions (bottom) with the AtSC, HvSC, LjSC and SSC (left to right). The relative abundance bars are colored by the inoculum that the isolates derive from. The X-axis provides information on the sample's nature (inoculum, host, compartment, nutrient condition, and experiment), for which the meaning can be found in supplemental dataset 3.

Pseudoalignment of reads to the Salmon indices can lead to read misalignment, with, e.g., reads from AtSC samples pseudoaligning to HvSC or LjSC indices. This does not necessarily mean LjSC or HvSC isolates are contaminating the AtSC samples, as seed endophyte-derived reads could have a high genomic similarity to LjSC or HvSC isolates, which would be reflected in these contamination levels. Despite mispseudoalignment of a number of reads, most evident in AtSC samples, we demonstrate that our data contains clean input inocula, and marginal contamination levels in the host-specific inocula.

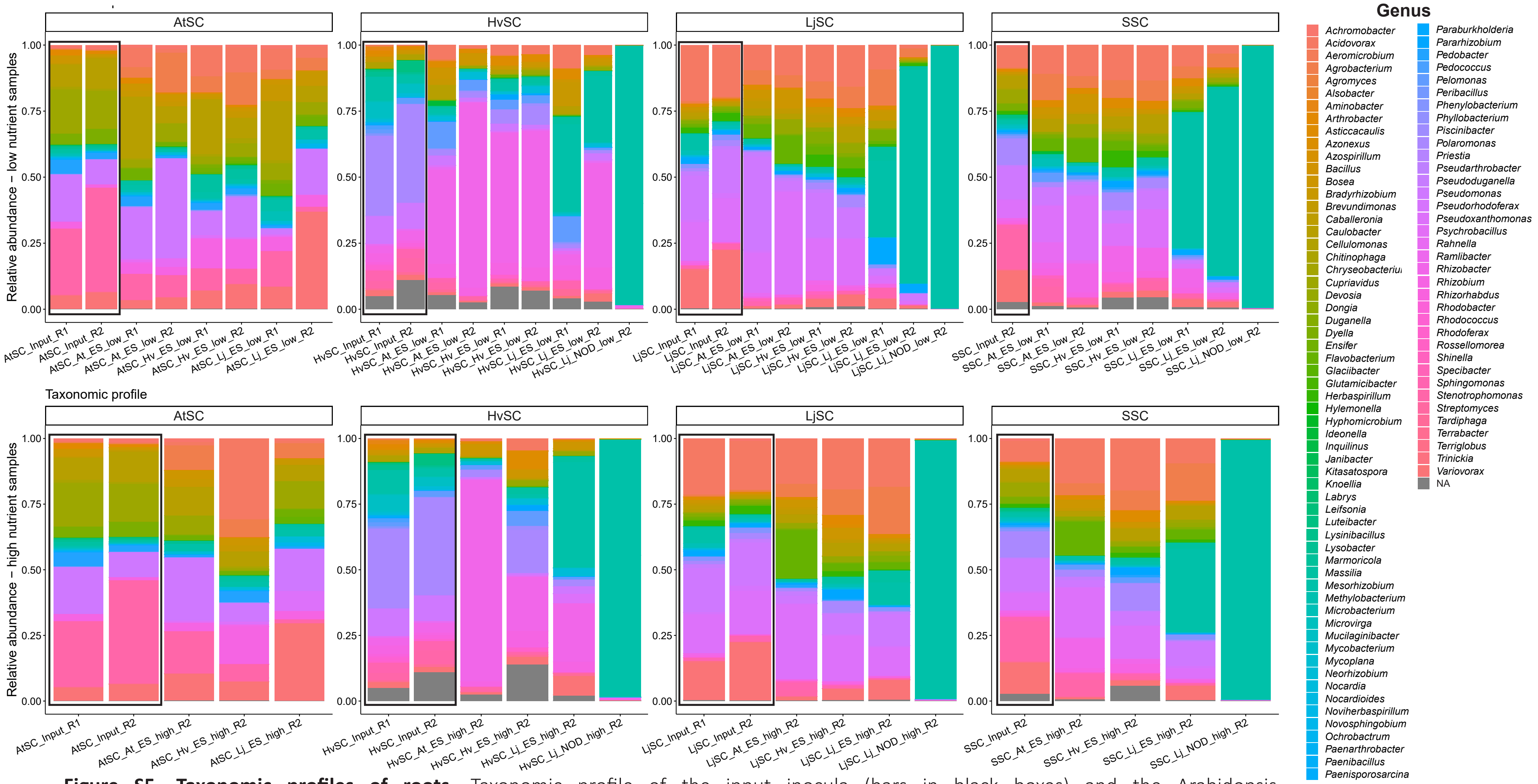

**Figure S5. Taxonomic profiles of roots.** Taxonomic profile of the input inocula (bars in black boxes) and the Arabidopsis, Barley and Lotus root microbiome when inoculated under low nutrient conditions (top) and high nutrient conditions (bottom) with the AtSC, HvSC, LjSC and SSC (left to right). The relative abundance bars are colored by the genus composition. The X-axis provides information on the sample’s nature (inoculum, host, compartment, nutrient condition, and experiment), for which the meaning can be found in supplemental dataset 3.

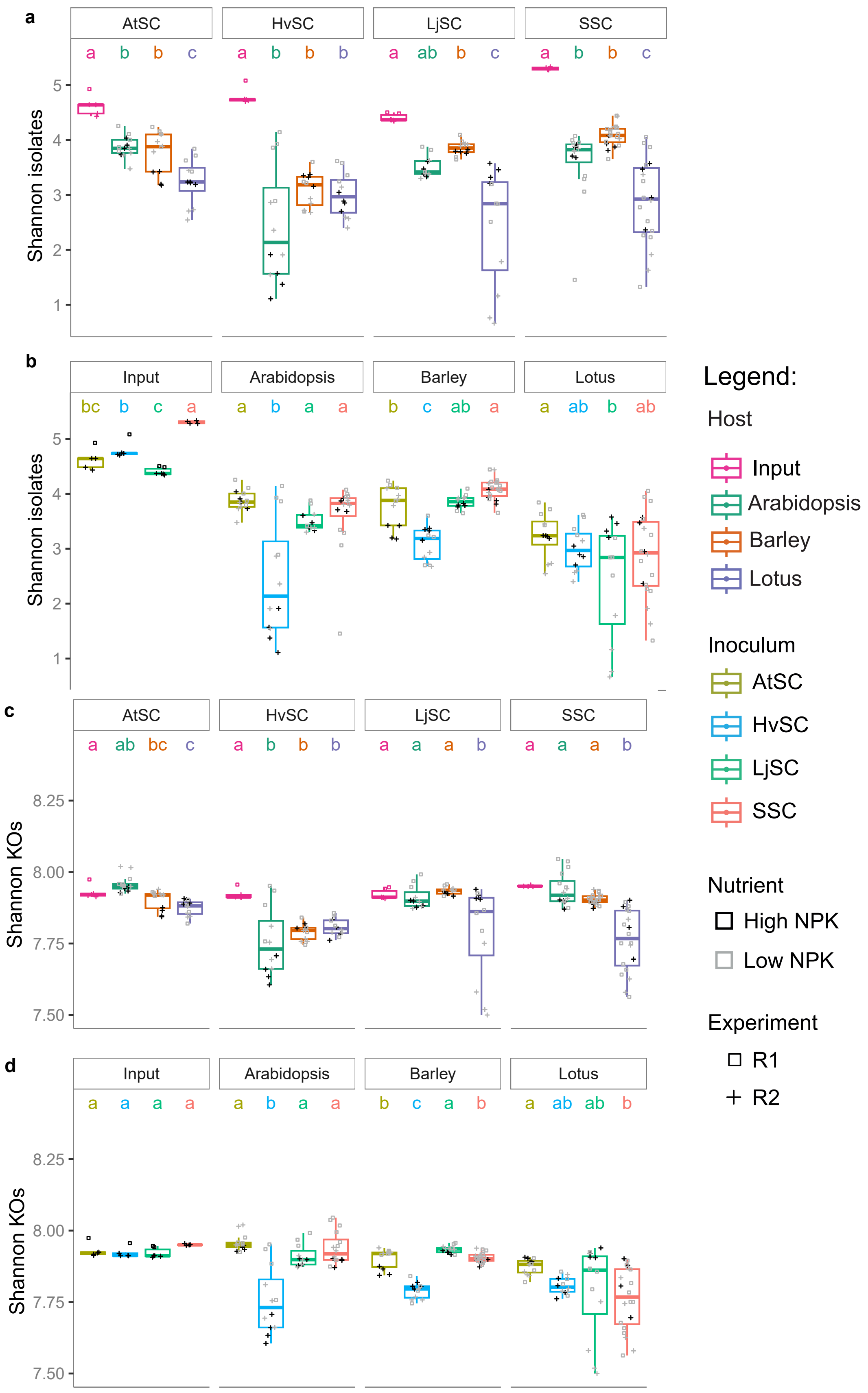

**Figure S6. Shannon diversity plots.** Shannon diversity on the isolates (a and b) and on the KOs (c and d) for each host and inoculum. Plots a and b are the same but the groups are grouped either by inoculum (a) or host (b) allowing statistical comparison between groups. This is similar for plots c and d. The alpha diversity values were statistically assessed with an ANOVA with post-hoc Tukey HSD test.

#### Supplemental Results 1 - Nutrient condition effect on diversity analyses

The reduction in number of isolates and KOs that the roots of Arabidopsis, Barley, and Lotus accommodate as compared to the initial inoculum is substantial (Figures 2b and c). The effect of the nutrient condition on root microbiome assembly in this analysis was found to be reduced and appears to differentially affect this reduction in specific host and inoculum contexts (Figure S7). Frequently, a lower nutrient condition seems to positively affect the number of isolates and KOs, most evident in the Arabidopsis and Lotus contexts. In the context of a host or inoculum-dependent enrichment of specific community members, illustrated by the enrichment of Lotus symbionts in the low nutrient Lotus-context and the enrichment of *Rhizobacter* isolate P2_G4 in the high nutrient HvSC-context, the communities are more uneven in terms of isolates and functions (Figure S7).

These findings suggest that a low nutrient environment can enrich for increased taxonomic and functional diversity, which could be related to an increased nutritional dependence of the plant on its microbiota (Singh *et al.*, 2022). In addition, oligotrophs may be prevented from dominating the microbiome in low-nutrient, scarce conditions, leading to more diverse communities (Wang *et al.*, 2018; Zeng *et al.*, 2016). However, in contrast, long-term nutrient enrichment has also been shown to increase bacterial diversity (Bledsoe *et al.*, 2020). The only exception to the correlation between nutrient concentration and bacterial diversity is when Lotus is grown in low-nutrient conditions, inevitably leading to the recruitment of symbionts to engage in nitrogen-fixing symbiosis, greatly affecting the microbiome composition (Zgadzaj *et al.*, 2016).

In line with the findings from Figures 2f and g, we demonstrate that the variation in microbiome composition of samples with the same treatment (host, inoculum, and nutrient condition) is lower for the functional composition as compared to the genus composition (Figure S8). This indicates that functional convergence not only occurs between hosts and inocula, but also within treatments. The variation in microbiome composition between nutrient conditions seems to be largely similar between high and low nutrient conditions, with slightly higher variation in the low nutrient condition (Figure S8). This supports the argument that the host's interactions with its microbiota are more stochastic when nutrients are limited, thus leading to more variation and diversity, though this difference between high and low nutrient conditions is only marginal. Interestingly, across inocula and nutrient conditions, more variation exists in the Arabidopsis and Lotus root microbiome as compared to Barley root microbiome. Surprisingly, the Barley root microbiome harbors the highest taxonomic and functional diversity (Figure 2b and d), which is relatively stable across nutrient conditions (Figure S7 and S8), while simultaneously displaying greater phenotypic disparity (Figure S1).

One potential driver of this difference in microbiome assembly might be related to domestication processes that cereal crops, such as Barley, have been subjected to. Breeding practices negatively affect the root microbiota's diversity in crops, as the process nullifies a million-year long process of evolution (Edwards *et al.*, 2015; Peiffer *et al.*, 2013). On the other hand, studies on domesticated grass species demonstrate that the microbial diversity is unaffected by domestication (Naylor *et al.*, 2017; Ling *et al.*, 2022).

#### *References*

Bledsoe, R. B., Goodwillie, C., & Peralta, A. L. (2020). Long-term nutrient enrichment of an oligotroph-dominated wetland increases Bacterial Diversity in bulk soils and plant rhizospheres. *MSphere*, *5*(3).

Edwards, J., Johnson, C., Santos-Medellín, C., Lurie, E., Podishetty, N. K., Bhatnagar, S., Eisen, J. A., & Sundaresan, V. (2015). Structure, variation, and assembly of the root-associated microbiomes of rice. *Proceedings of the National Academy of Sciences*, *112*(8).

Ling, N., Wang, T., & Kuzyakov, Y. (2022). Rhizosphere bacteriome structure and functions. *Nature Communications*, *13*(1), 836.

Naylor, D., DeGraaf, S., Purdom, E., & Coleman-Derr, D. (2017). Drought and host selection influence bacterial community dynamics in the grass root microbiome. *The ISME Journal*, *11*(12), 2691–2704.

Peiffer, J. A., Spor, A., Koren, O., Jin, Z., Tringe, S. G., Dangl, J. L., Buckler, E. S., & Ley, R. E. (2013). Diversity and heritability of the maize rhizosphere microbiome under field conditions. *Proceedings of the National Academy of Sciences*, *110*(16), 6548–6553.

Singh, S. K., Wu, X., Shao, C., & Zhang, H. (2022). Microbial enhancement of plant nutrient acquisition. *Stress Biology*, *2*(1), 3.

Wang, C., Liu, D., & Bai, E. (2018). Decreasing soil microbial diversity is associated with decreasing microbial biomass under nitrogen addition. Soil Biology and Biochemistry, 120, 126-133.

Zeng, J., Liu, X., Song, L., Lin, X., Zhang, H., Shen, C., & Chu, H. (2016). Nitrogen fertilization directly affects soil bacterial diversity and indirectly affects bacterial community composition. Soil Biology and Biochemistry, 92, 41-49.

Zgadzaj, R., Garrido-Oter, R., Jensen, D. B., Koprivova, A., Schulze-Lefert, P., & Radutoiu, S. (2016). Root nodule symbiosis in *Lotus japonicus* drives the establishment of distinctive rhizosphere, root, and nodule bacterial communities. *Proceedings of the National Academy of Sciences*, *113*(49).

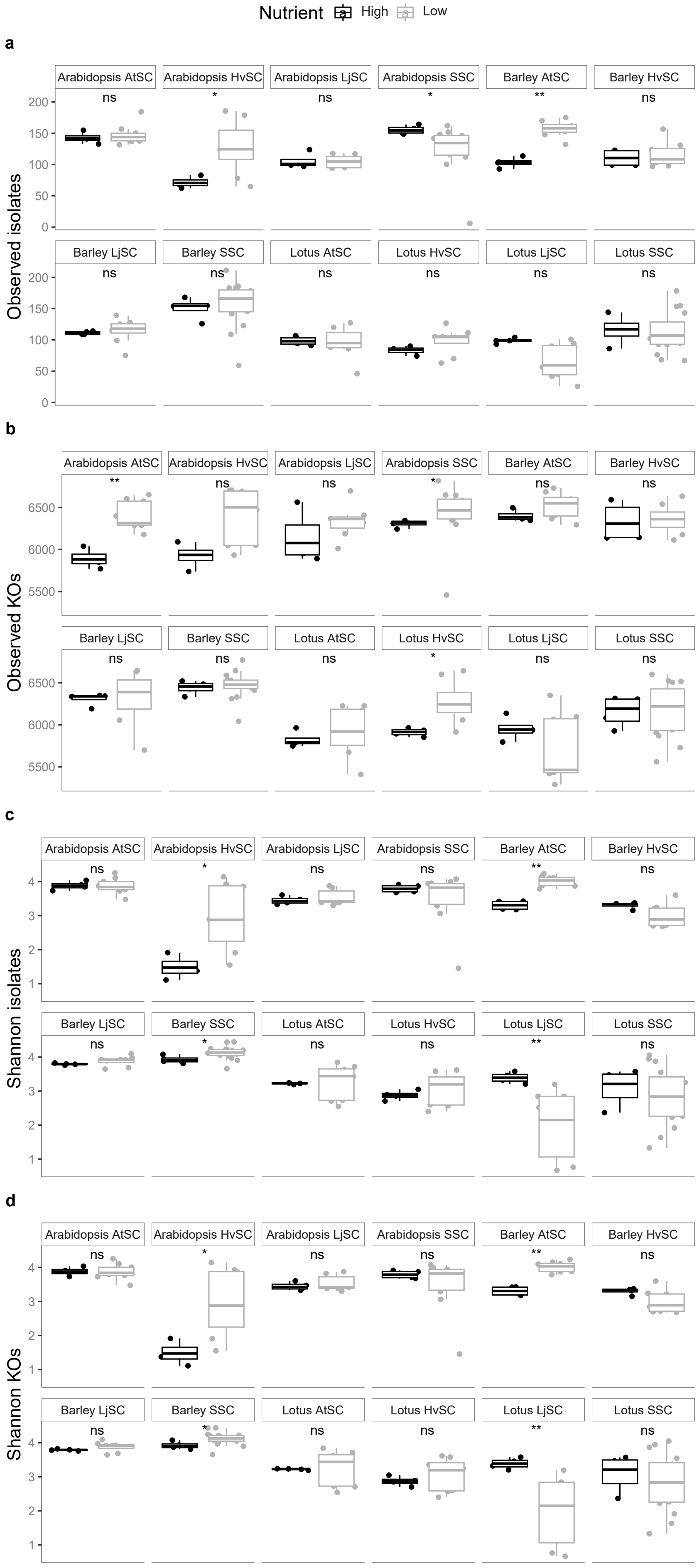

**Figure S7. Nutrient condition effect on alpha diversity plots.** The data is subsetted to show the observed isolates (a), observed KOs (b), shannon diversity for isolates (c) and shannon diversity for KOs (d) between low and high nutrient conditions. Statistical differences are indicated by * assessed by a Wilcoxon test (p-value < 0.05).

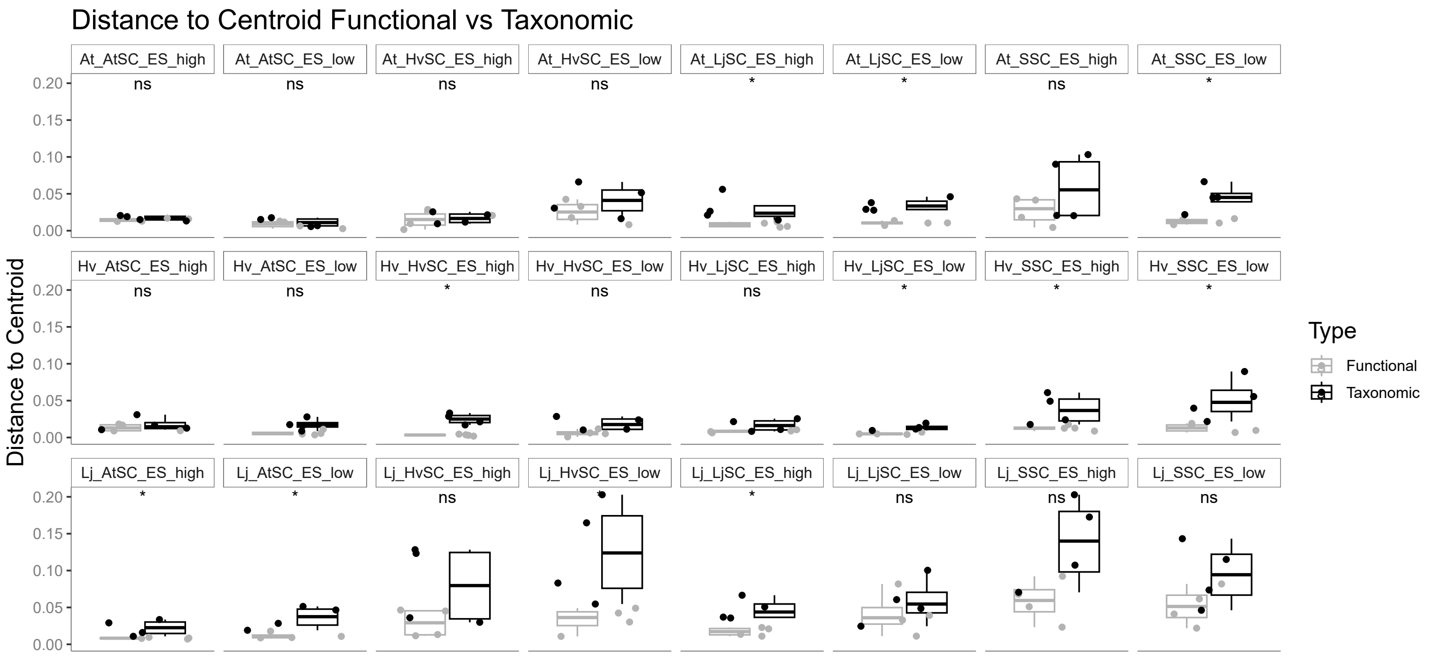

**Figure S8. Distance to centroid for genera and functions.** Each boxplot panel indicates the distance between the individual samples and the centroid of the whole group of samples for both the genus and KO composition shown in figure 2f and g (only for the SSC experiment R2). A high distance indicates a greater compositional variation between replicates of the same treatment. Statistical differences are indicated by * assessed by a Wilcoxon test (p-value < 0.05). The panel titles provide information on the sample's nature (host, inoculum, compartment, and nutrient condition), for which the meaning can be found in supplemental dataset 3.

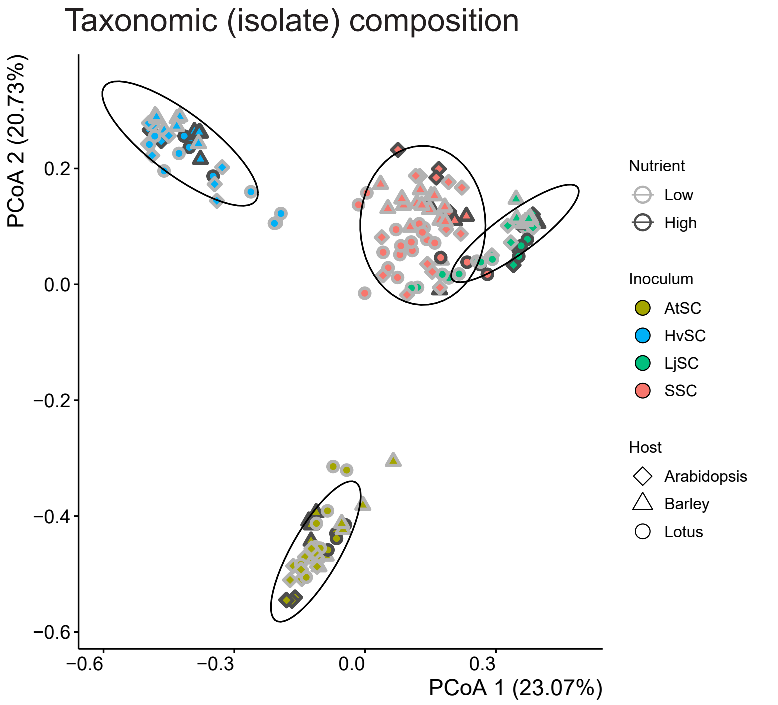

| Parameter | Taxonomy R² |
|---|---|
| Inoculum | 0.44 |
| Host | 0.13 |
| Inoculum:Host | 0.12 |
| Nutrient | 0.01 |
| Experiment | 0.02 |

**Figure S9. Beta diversity analysis on isolate composition.** Bray-Curtis PCoA on the isolate composition of Arabidopsis, Barley and Lotus roots (shapes) inoculated with the four inocula (colors) under high and low nutrient conditions (peripheral colors). The right panel shows the adonis $R^2$ values, indicating what proportion of the compositional differences is explained by the different metadata variables. All the indicated parameters are significant (p<0.001)

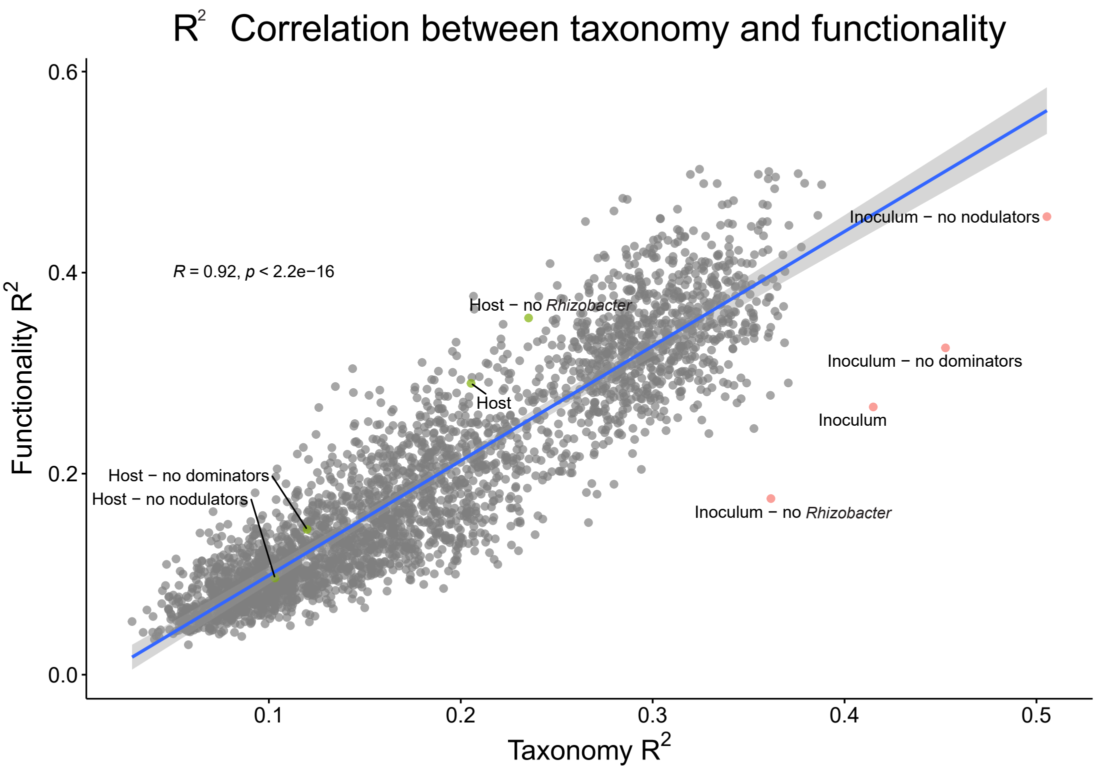

**Figure S10. Computational simulation of randomly assembled root microbiomes to assess taxonomic versus functional $R^2$.** The correlation between the $R^2$ on genus composition (X-axis) and the $R^2$ on the KO composition (Y-axis) based on 1000 microbiome simulations. The $R^2$ values of figure 2f and g are added to indicate deviations of the correlation and thus what the host's selection effect is; the inoculum $R^2$ values (red) and the host $R^2$ values (green) of the dataset with and without the Lotus symbionts and/or the HvSC dominant *Rhizobacter* sp. P2_G4.

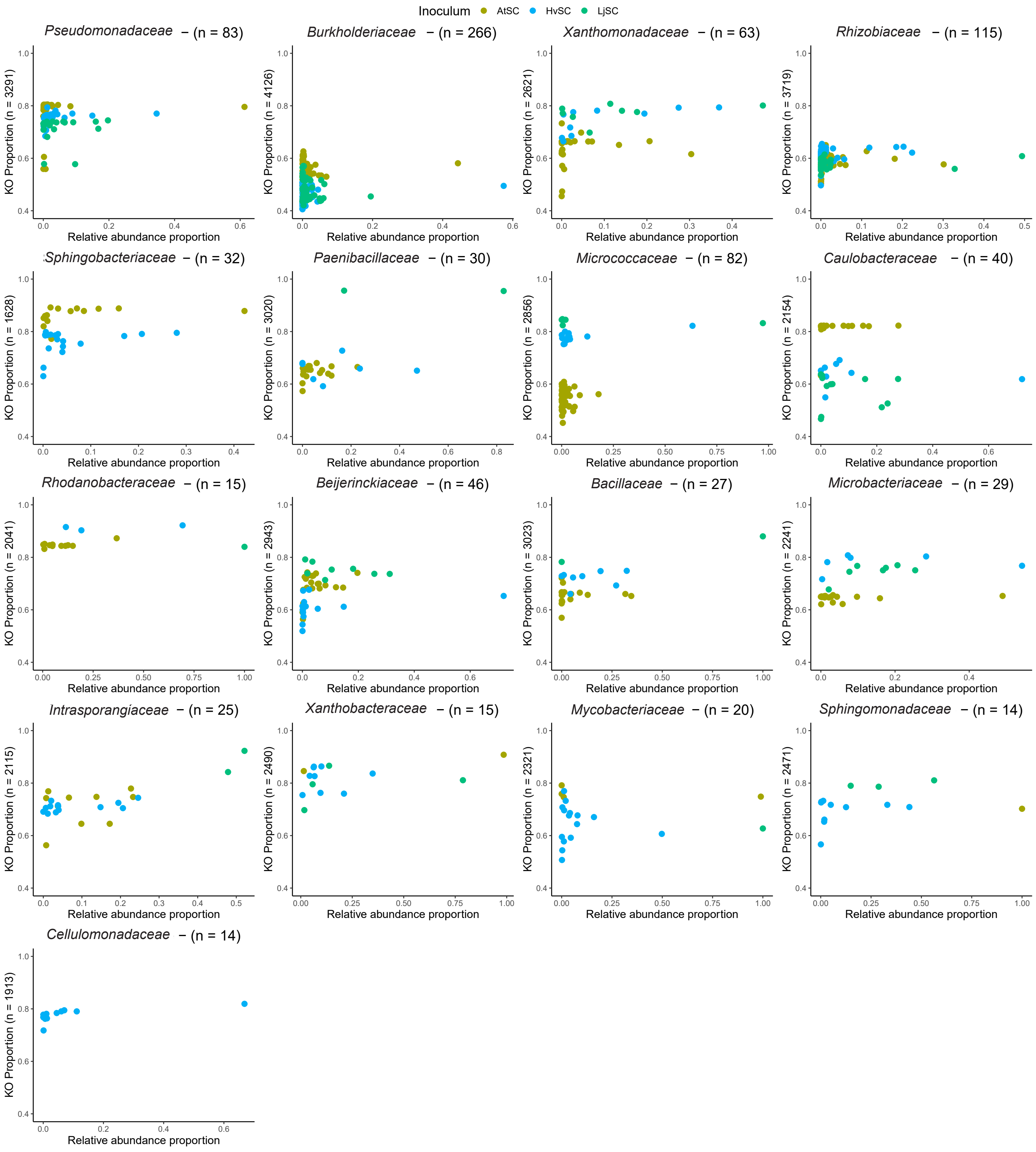

**Figure S11. Correlation KO diversity and relative abundance per family.** The relation between genomic diversity of isolates (Y-axis) and the relative abundance (X-axis) in the dataset subsetted by family. The KO proportion indicates the number of KOs an isolate has as compared to the number of KOs in the family pangenome. The relative abundance proportion indicates the isolate's relative abundance as compared to the family's relative abundance. The numbers in the brackets indicate the number of isolates in the family (plot titles) and number of KOs in the family pangenome (Y-axes). Colors indicate from which inoculum the isolates originate. Correlations calculated in each these plots are displayed in the heatmap of figure 3b.

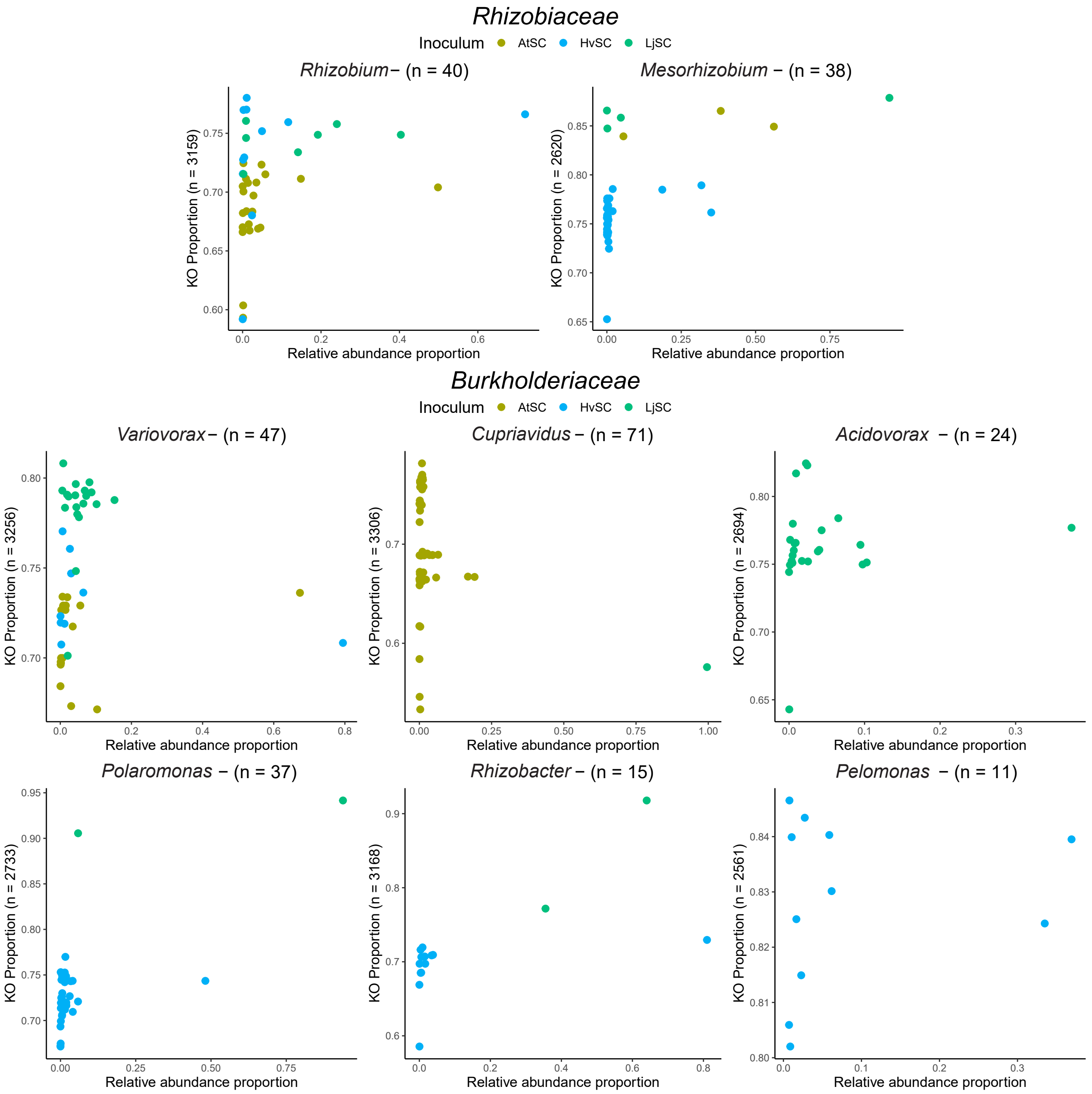

**Figure S12. Correlation KO diversity and relative abundance per genus in the two largest families.** The relation between genomic diversity of isolates (Y-axis) and the relative abundance (X-axis) in the dataset subsetted per genus in the *Rhizobiaceae* (top) and *Burkholderiaceae* (bottom). The KO proportion indicates the number of KOs in an isolate compared to the number of KOs in the genus pangenome. The relative abundance proportion indicates the isolate's relative abundance as compared to the genus' relative abundance. The colors indicate from which inoculum the isolate derives from.

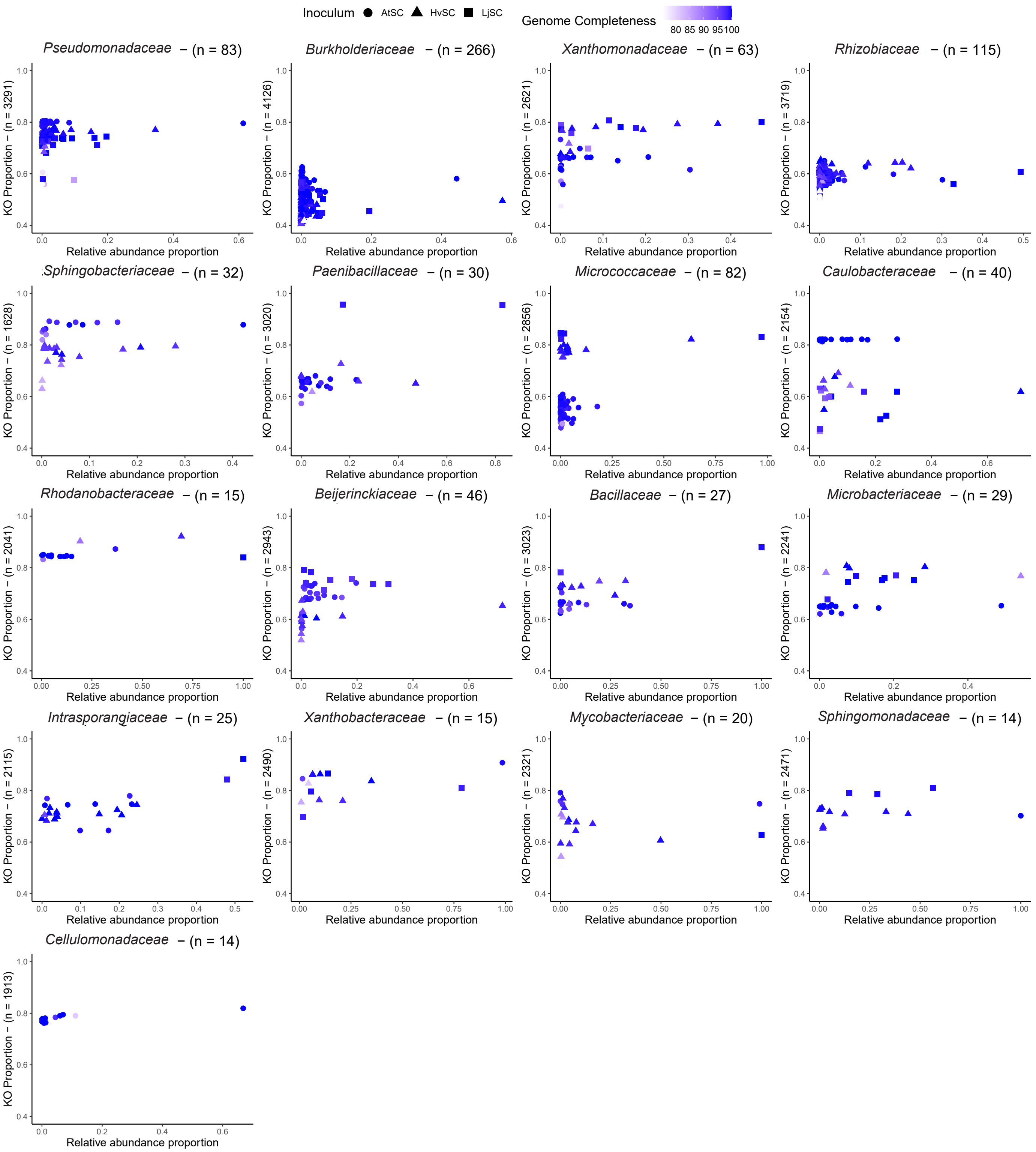

**Figure S13. Functional diversity of isolates as compared to genome completeness.** The relation between genomic diversity of isolates (Y-axis) and the relative abundance (X-axis) in the dataset subsetted per family. The KO proportion indicates the number of KOs in an isolate compared to the number of KOs in the family pangenome. The relative abundance proportion indicates the isolate's relative abundance as compared to the family's relative abundance. The colors indicate how complete the genome was to account for potentially missing parts of the genome that incidentally lead to a lower KO proportion and the shapes indicate from which inoculum the isolate is derived from.

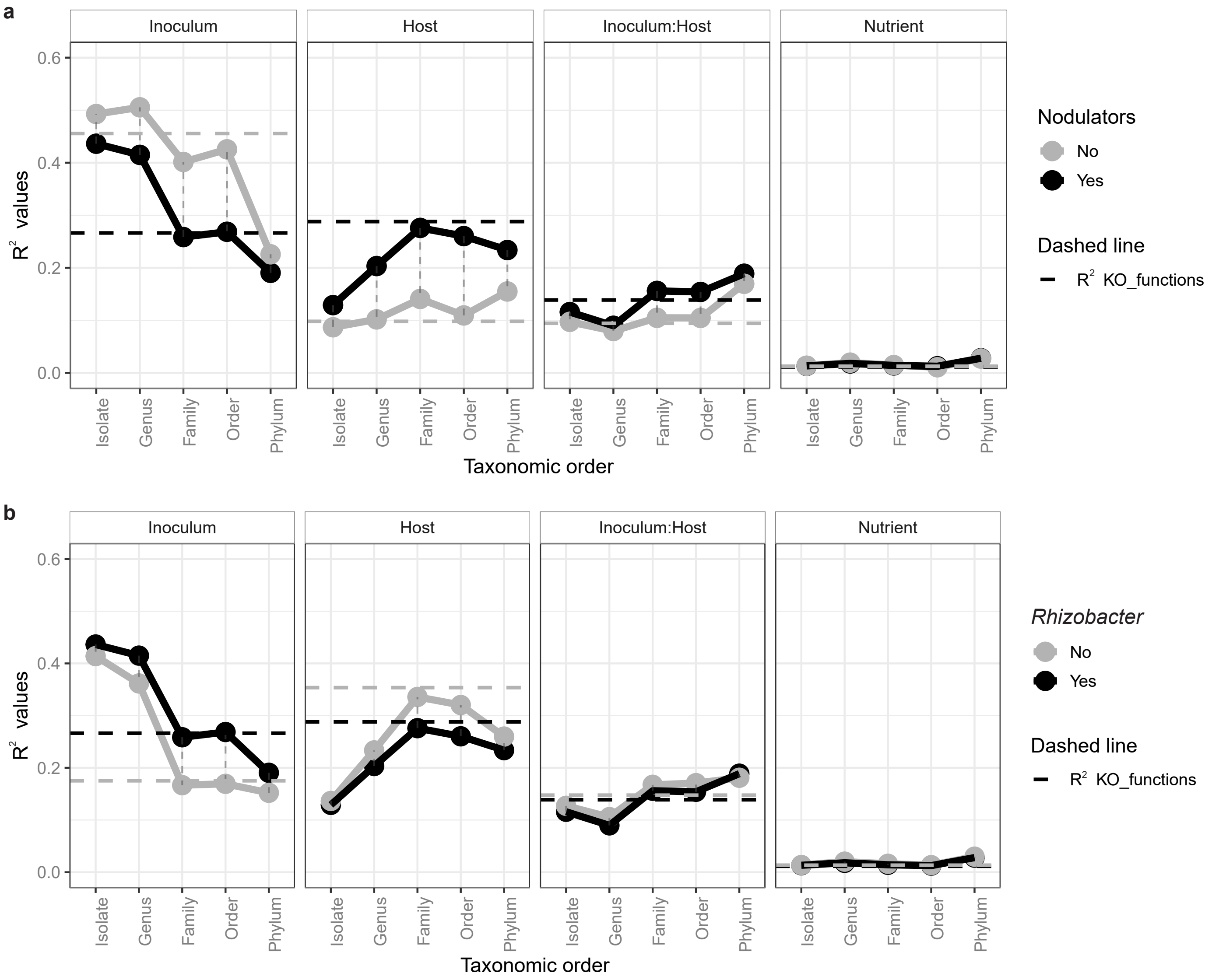

**Figure S14. Compositional variation on functional and taxonomic diversity with and without dominant isolates.** The effect of the inoculum, the host, the inoculum:host interaction, and the nutrient condition (from left to right) on the taxonomic composition on different levels as well as the functional composition. The more transparent lines indicate the data when the Lotus symbionts and their genes are excluded (a) or when the HvSC *Rhizobacter* sp. P2_G4 and its genes were excluded (b). The $R^2$ values are calculated by an Adonis test.

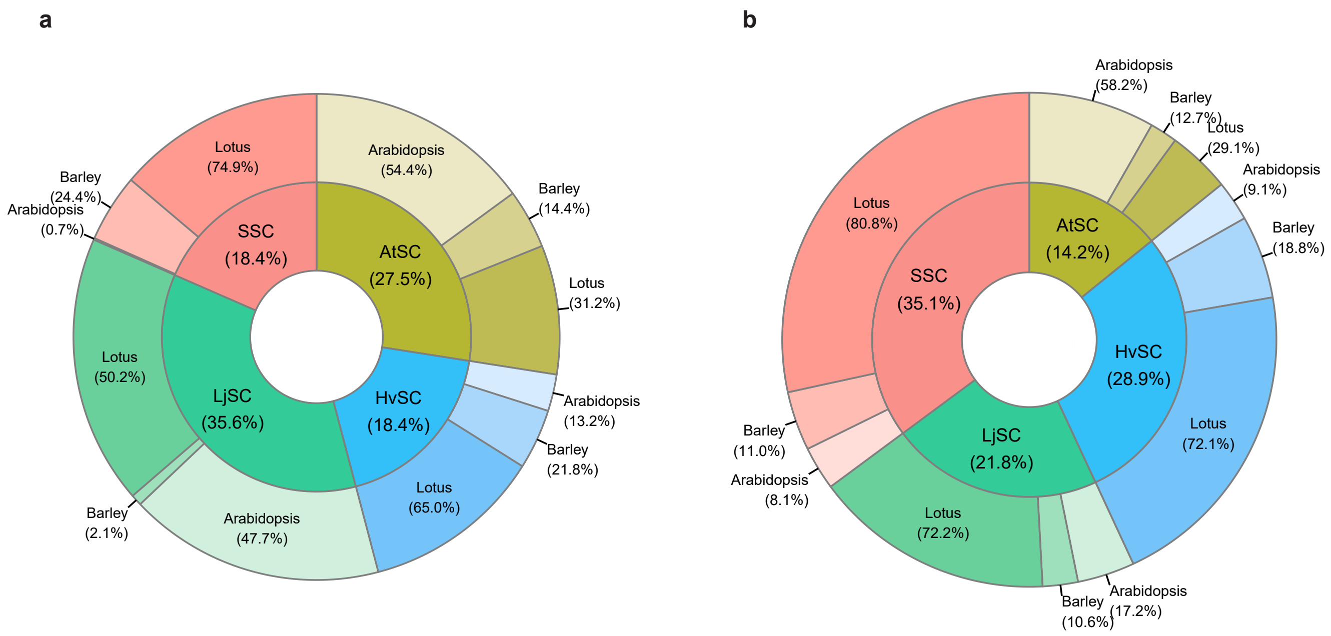

**Figure S15. Host's effect on compositional differences without Lotus symbionts or *Rhizobacter* sp. P2_G4.** Piedonut plot in which the pie chart indicates how much the hosts altogether affect the functional compositional differences in each inoculum. The percentages indicate in which inoculum the host compositional differences were the biggest or smallest. The donut plot indicates per inoculum the effect of each host to the compositional differences. (a) The piedonut plot that excludes the Lotus symbionts. (b) The piedonut plot that excludes the HvSC *Rhizobacter* sp. P2_G4. The original $R^2$ values that these percentages reflect can be found in figure S14a and b.

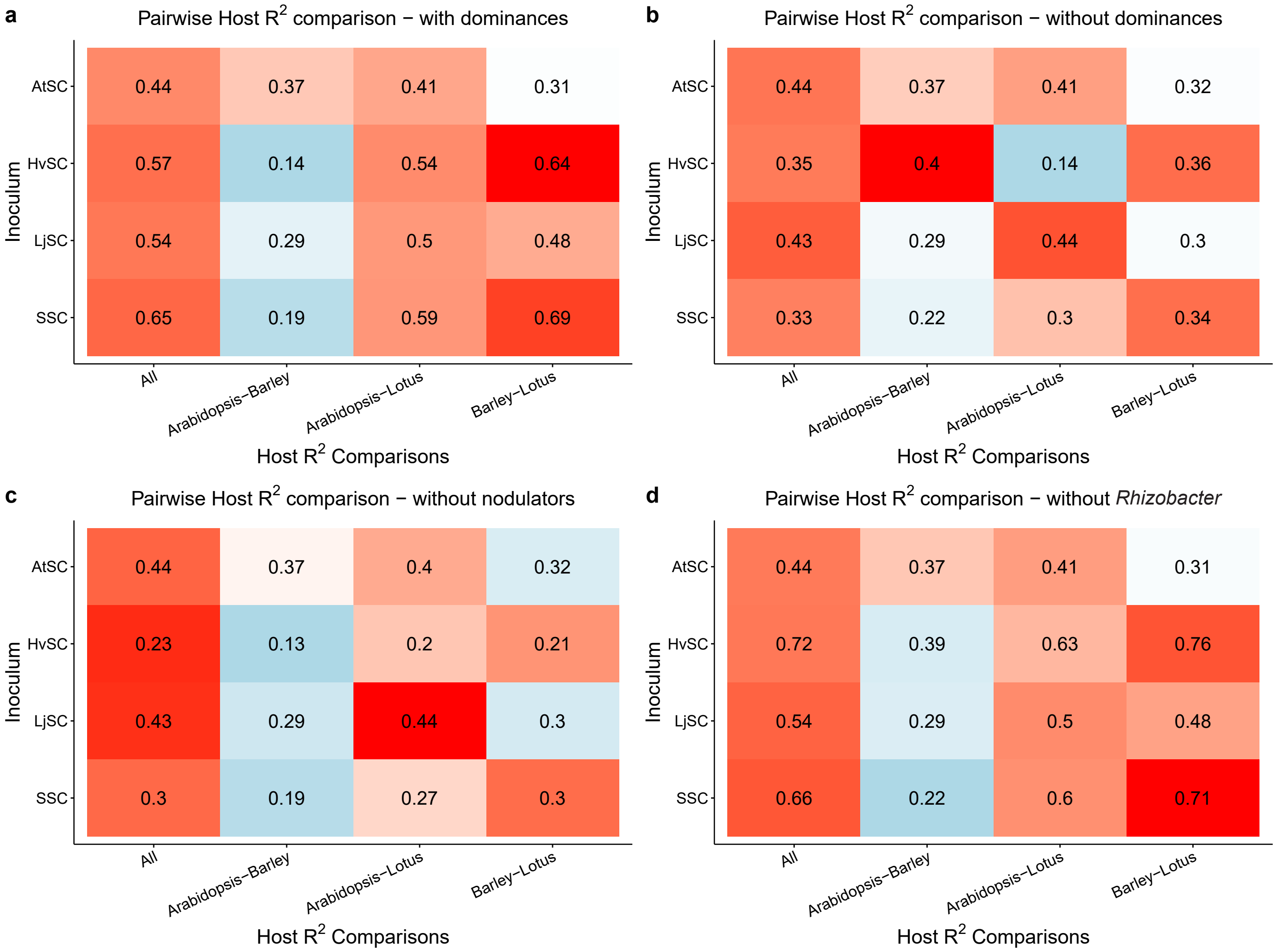

**Figure S16. Host effect on functional $R^2$ in each inoculum.** The effect of the hosts on the functional composition when subsetted for inocula (in the rows) in the original dataset (a), the dataset without the Lotus symbionts and HvSC *Rhizobacter* sp. P2_G4 (b), or the dataset without the Lotus symbionts (c) or *Rhizobacter* sp. P2_G4 (d). The values indicate the host $R^2$ with all hosts included (first column) or when a host is dropped out of the analysis (other columns). The changes in taxonomic host $R^2$ can be found in figures S17, S18, and S19 for the dataset without both the Lotus symbionts and *Rhizobacter* sp. P2_G4, without the Lotus symbionts and without the *Rhizobacter* sp. P2_G4 respectively.

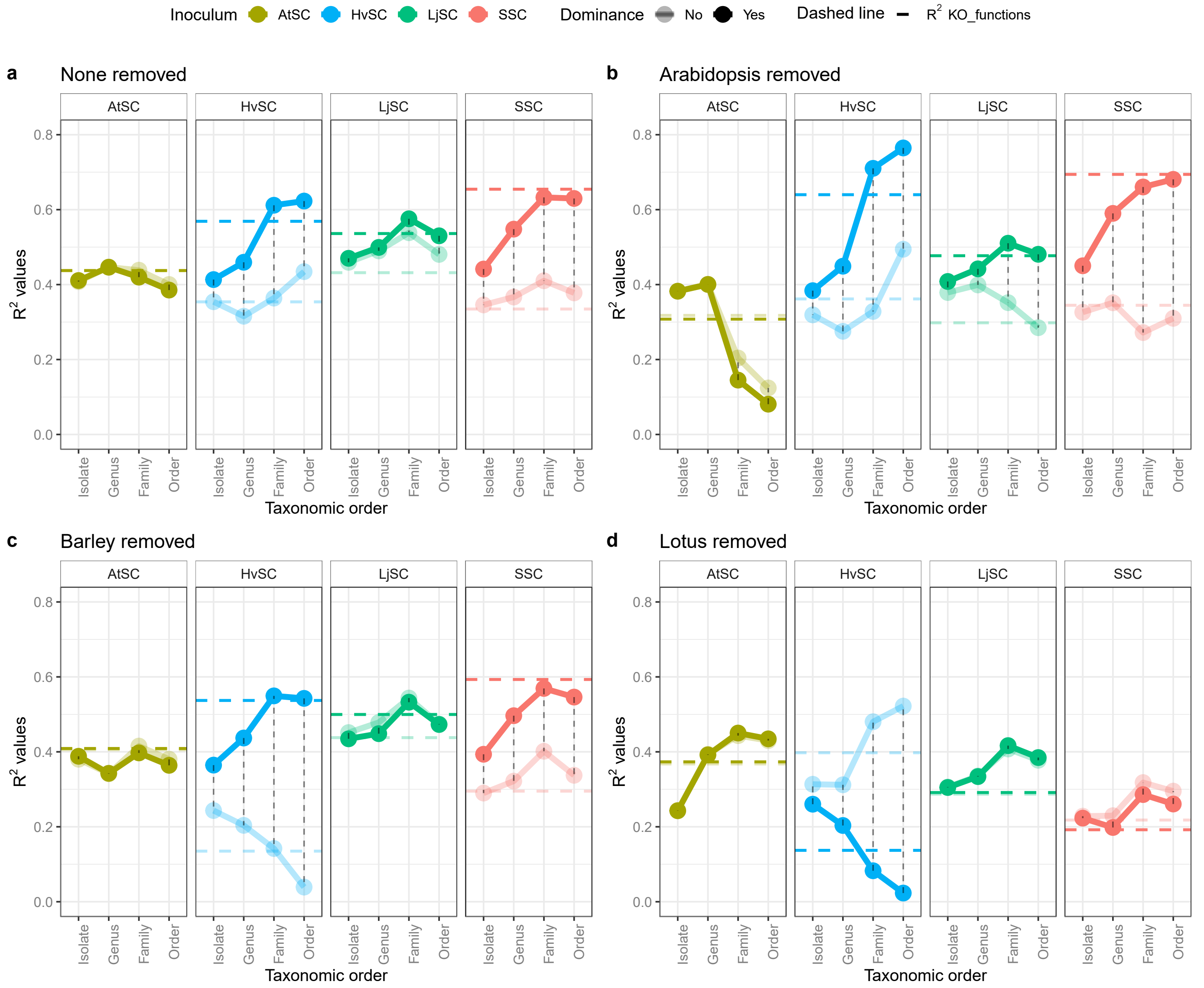

**Figure S17. Host effect on the taxonomic and functional composition with and without dominant isolates.** The effect of the host on the microbial composition on different taxonomic levels and the functional composition (a). The more transparent lines indicate the data where the Lotus symbionts and the dominant HvSC *Rhizobacter* sp. P2_G4 and their genes are excluded. The effect of each host is additionally assessed by removing it from the dataset computationally and investigate the $R^2$ for the two remaining hosts, where either Arabidopsis (b), Barley (c) or Lotus (d) was removed from the dataset. The $R^2$ values are calculated by an Adonis test.

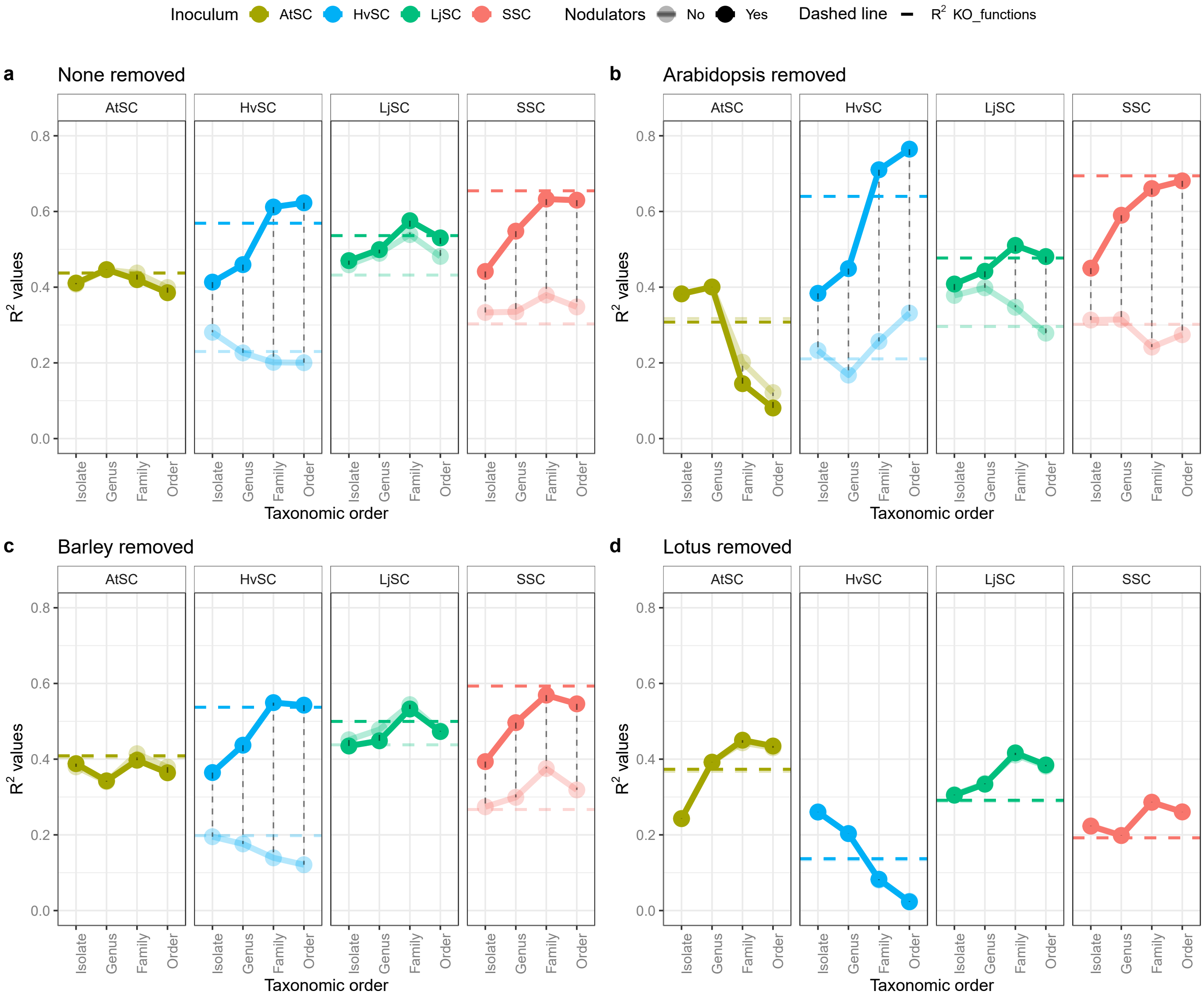

**Figure S18. Host effect on the taxonomic and functional composition with and without nodulator isolates.** The effect of the host on the microbial composition on different taxonomic levels and the functional composition (a). The more transparent lines indicate the data where the Lotus symbionts and their genes are excluded. The effect of each host is additionally assessed by removing it from the dataset computationally and investigate the $R^2$ for the two remaining hosts, where either Arabidopsis (b), Barley (c) or Lotus (d) was removed from the dataset. The $R^2$ values are calculated by an Adonis test.

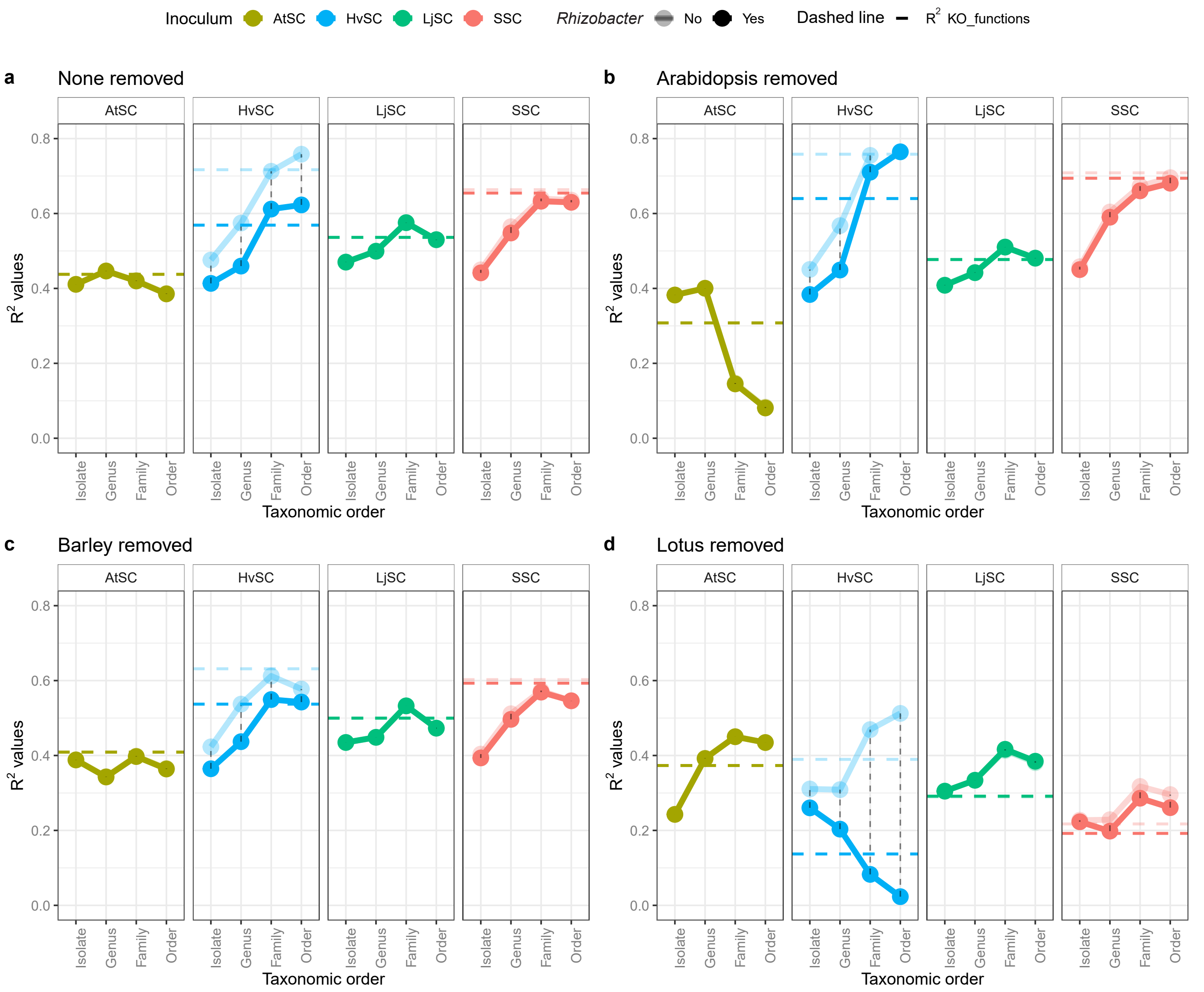

**Figure S19. Host effect on the taxonomic and functional composition with and without *Rhizobacter* sp. P2_G4.** The effect of the host on the microbial composition on different taxonomic levels and the functional composition (A). The more transparent lines indicate the data where the dominant HvSC *Rhizobacter* sp. P2_G4 and its genes are excluded. The effect of each host is additionally assessed by removing it from the dataset computationally and investigate the $R^2$ for the two remaining hosts, where either Arabidopsis (b), Barley (c) or Lotus (d) was removed from the dataset. The $R^2$ values are calculated by an Adonis test.

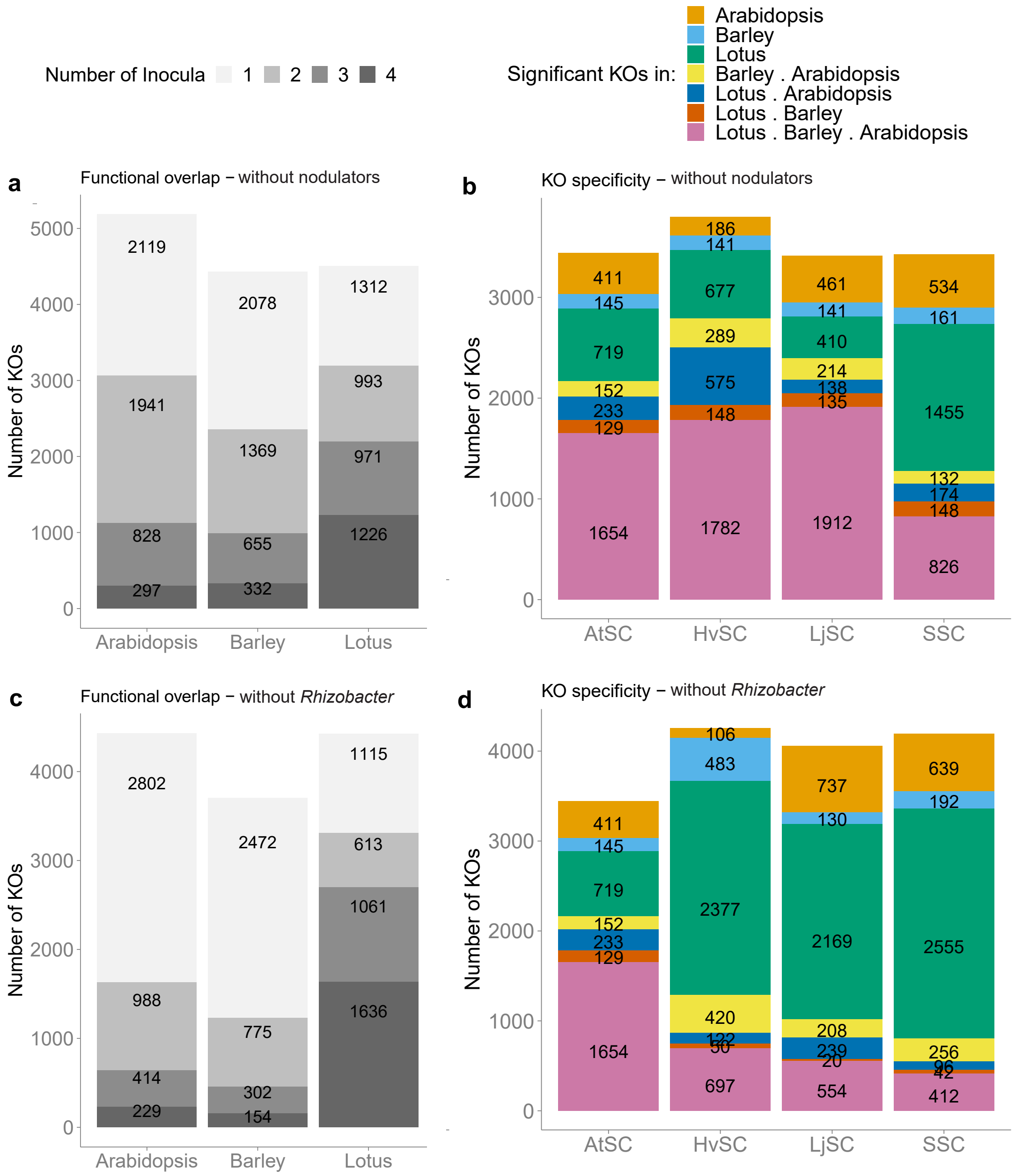

**Figure S20. Overlap and number of root microbiome-enriched KOs across hosts and inocula.** The overlap in differentially abundant bacterial KOs recruited by Arabidopsis, Barley or Lotus compared to the initial inoculum. Overlap between the four inocula and the three hosts shown in number of differentially abundant KOs in the dataset without the Lotus symbionts (a and b respectively), or the dataset without the *Rhizobacter* sp. P2_G4 (c and d respectively).

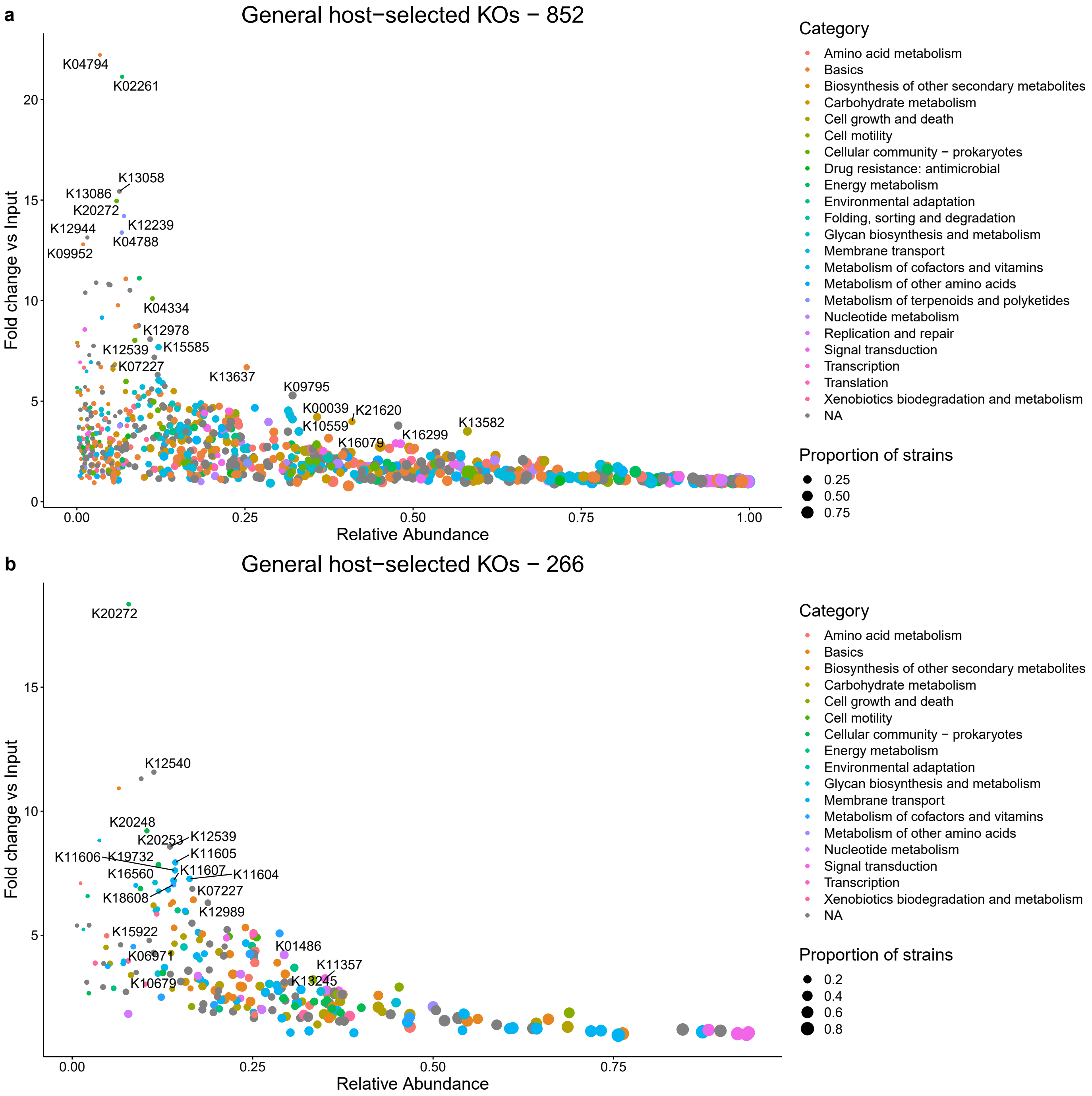

**Figure S21. General enriched KOs in the root microbiome.** KOs that are selected by the host from the DESeq2 analysis are plotted regarding the relative abundance of isolates that have the KO (X-axis) and how this relative abundance changed in comparison to the initial inoculum (Y-axis). The coordinate sizes indicates the proportion of isolates that have the KO in the SSC and the color indicates the category in which the KO functions. (a) The general KOs from the lenient DESeq2 selection (significant KOs in >6/12 host inoculum combinations). (b) The general KOs from the strict DESeq2 selection can be found (significant KOs in 12/12 host inoculum combinations in the dataset without the Lotus symbionts and *Rhizobacter* sp. P2_G4)

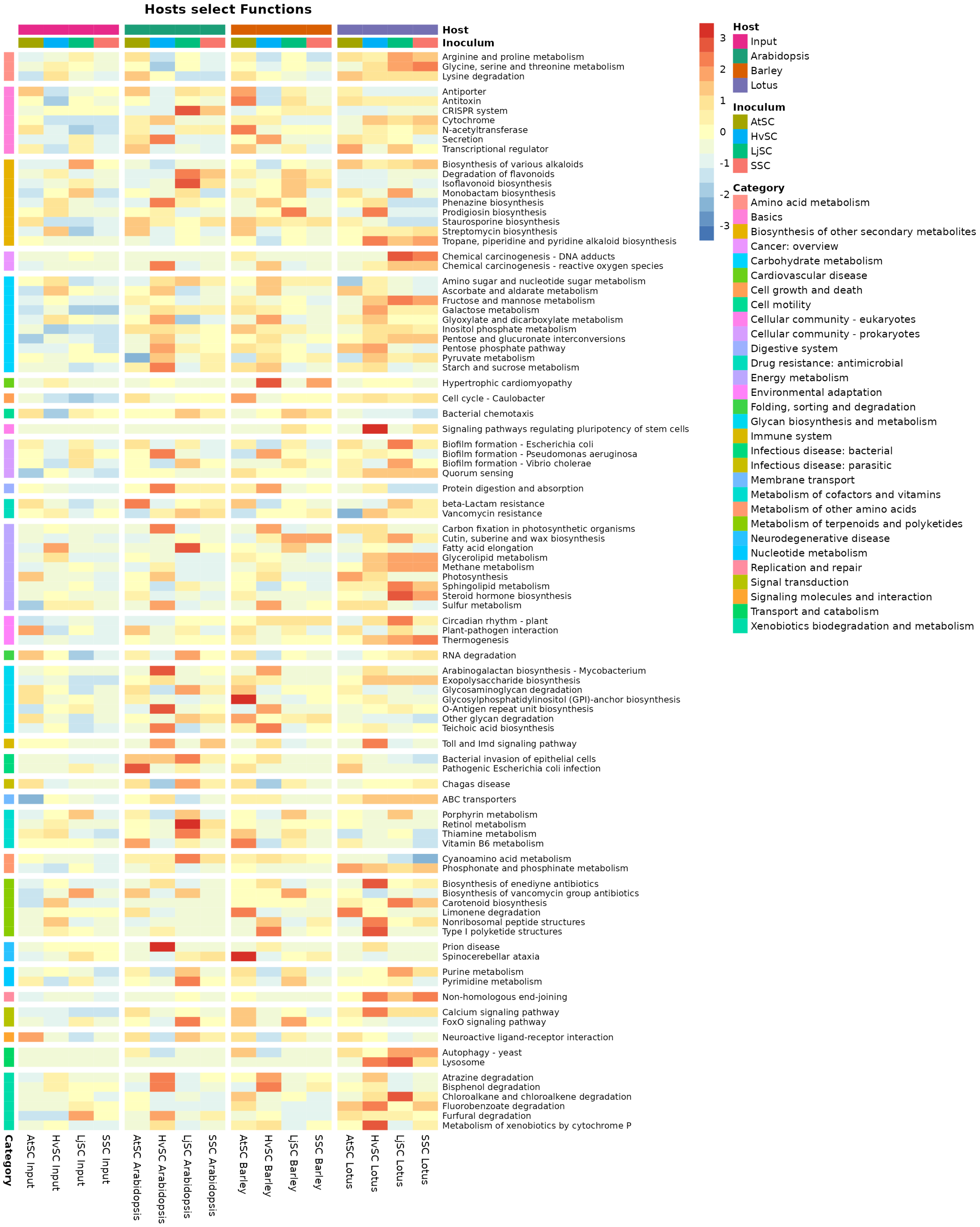

**Figure S22. Pathway abundances of KOs that are generally selected for by Arabidopsis, Barley, and Lotus.** A heatmap on pathway abundances that is scaled on the rows (pathways) for every combination of input/host and inoculum. The heatmap values were calculated by averaging the KO abundances of all significant KOs in a pathway from the lenient DESeq2 selection (significant in >6/12 host inoculum combinations). The annotation colors indicate the pathway category (rows) and the sample type (column).

### Supplemental Results 2 – General host-selected functions – the 266 KO set

Computational exclusion of the KOs from the Lotus symbionts and those from *Rhizobacter* sp. P2_G4 followed by analysis of the remaining significant KOs across all combinations of host and inoculum yielded 266 strictly enriched KOs (Table S5) (Figure S23c and d). The 57 pathways that these 266 KOs encode mainly relate to metabolism, involving carbohydrate, amino acid, energy, glycan, cofactors and vitamins, and xenobiotics metabolism (Figure S23e and f), but also quorum sensing and two-component system-related functions. Several bacteria in our dataset possess a large proportion of these 266 KOs (Figure S23g), with 37 bacterial strains possessing ≥ 200 KOs. Interestingly, several plant-associated bacteria described by Levy *et al.* (2018), possess +/- 150-160 of these KOs (Figure S23h), while nearly none of the non-plant-associated and soil bacteria possess these. Thus, it is conceivable that these bacteria are highly root competent.

Interestingly, the most significant pathway with regard to the fold change of bacterial isolates from the initial inoculum to the root microbiome relates to "secretion". This is mainly the result of one type VI secretion system-related gene, *impN*, which is thought to play a role in bacterial interspecies competition and host manipulation (Ma *et al.*, 2014; Bernal *et al.*, 2018). This finding corroborates recent observations where enrichment of mainly type III and type VI secretion systems was identified via computational meta-analysis of rhizosphere and soil metagenomes (Fourie *et al.*, 2024). In addition, five type IV secretion system proteins can be found among the set of 266 KOs, which are also involved in host manipulation strategies, both to induce symbiotic relationships (Hubber *et al.*, 2004 ; Hubber *et al.,* 2007; Wangthaisong *et al.*, 2023) as well as pathogenic relationships (Li & Christie, 2018). Interestingly, it has recently been shown that type IV secretion systems and corresponding type IV effectors can kill other bacterial cells, which could provide a competitive advantage in the root ecosystem (Purtschert-Montenegro *et al.*, 2022).

*References*

Bernal, P., Llamas, M. A., & Filloux, A. (2018). Type VI secretion systems in plant-associated bacteria. *Environmental Microbiology*, *20*(1), 1–15.

Fourie, A., López, J. L., Sánchez-Gil, J. J., Poppeliers, S. W. M., de Jonge, R., & Dutilh, B. E. (2024). Bacterial family-specific enrichment and functions of secretion systems in the rhizosphere. *BioRxiv*, 2024–2025.

Ma, L.-S., Hachani, A., Lin, J.-S., Filloux, A., & Lai, E.-M. (2014). Agrobacterium tumefaciens deploys a superfamily of type VI secretion DNase effectors as weapons for interbacterial competition *in planta*. *Cell Host & Microbe*, *16*(1), 94–104.

Hubber, A., Vergunst, A. C., Sullivan, J. T., Hooykaas, P. J., & Ronson, C. W. (2004). Symbiotic phenotypes and translocated effector proteins of the *Mesorhizobium loti* strain R7A VirB/D4 type IV secretion system. Molecular microbiology, 54(2), 561–574.

Hubber, A. M., Sullivan, J. T., & Ronson, C. W. (2007). Symbiosis-induced cascade regulation of the *Mesorhizobium loti* R7A VirB/D4 type IV secretion system. Molecular plant-microbe interactions, 20(3), 255–261.

Li, Y. G., & Christie, P. J. (2018). The *Agrobacterium* VirB/VirD4 T4SS: Mechanism and Architecture Defined Through In Vivo Mutagenesis and Chimeric Systems. Current topics in microbiology and immunology, 418, 233–260.

Purtschert-Montenegro, G., Cárcamo-Oyarce, G., Pinto-Carbó, M., Agnoli, K., Bailly, A., & Eberl, L. (2022). *Pseudomonas putida* mediates bacterial killing, biofilm invasion and biocontrol with a type IVB secretion system. *Nature Microbiology*, *7*(10), 1547–1557.

Wangthaisong, P., Piromyou, P., Songwattana, P., Wongdee, J., Teamtaisong, K., Tittabutr, P., Boonkerd, N., & Teaumroong, N. (2023). The type IV secretion system (T4SS) mediates symbiosis between *Bradyrhizobium* sp. SUTN9-2 and legumes. *Applied and Environmental Microbiology*, *89*(6).

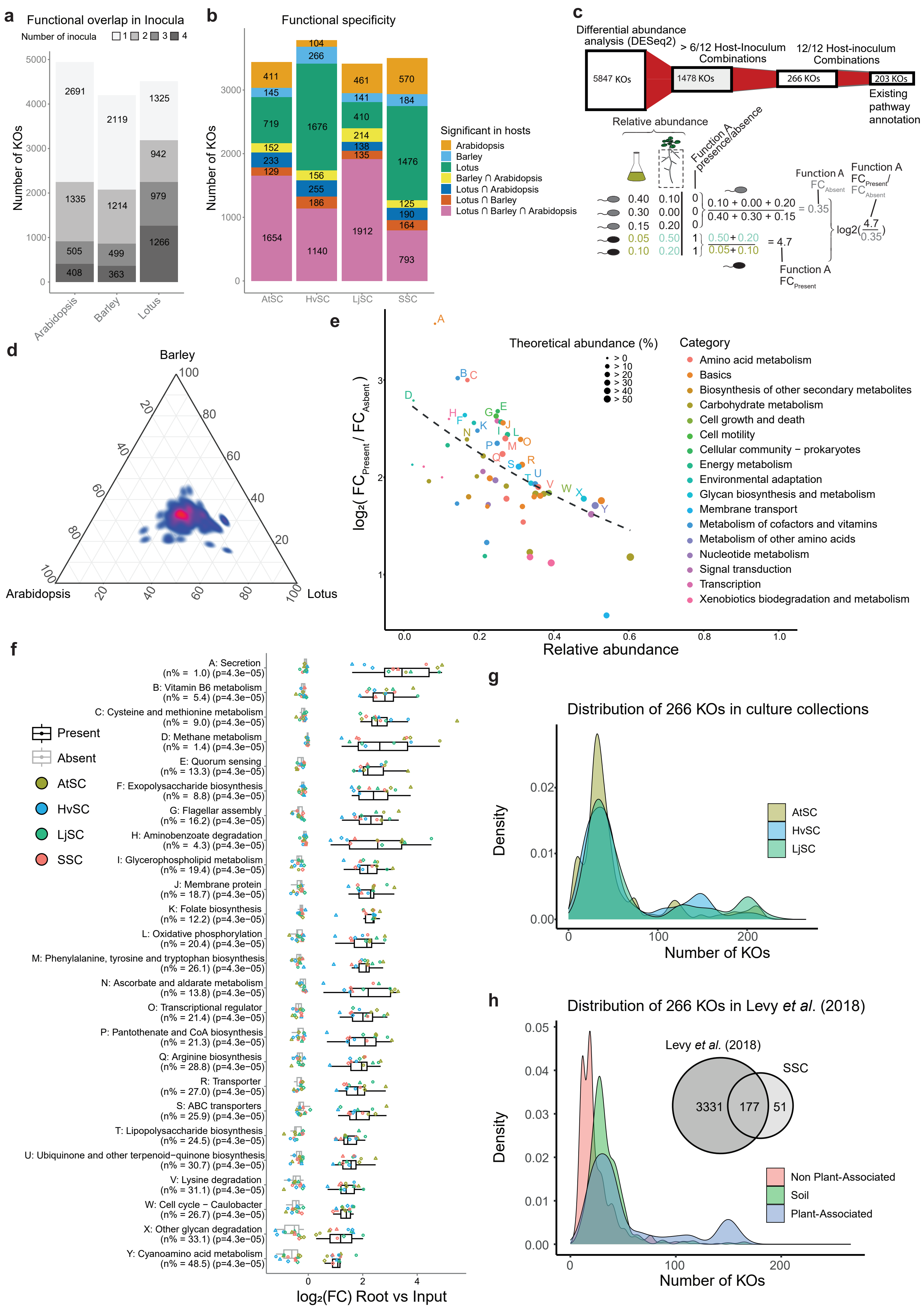
a
Functional overlap in Inocula
Number of inocula 1 2 3 4
Number of KOs
2691
1335
505
408
2119
1214
499
363
1325
942
979
1266
Arabidopsis
Barley
Lotus
b
Functional specificity
Number of KOs
411
145
719
152
233
129
1654
104
266
1676
156
255
186
1140
461
141
410
214
138
135
1912
570
184
1476
125
190
164
793
AtSC
HvSC
LjSC
SSC
Significant in hosts
Arabidopsis
Barley
Lotus
Barley ∩ Arabidopsis
Lotus ∩ Arabidopsis
Lotus ∩ Barley
Lotus ∩ Barley ∩ Arabidopsis
c
Differential abundance analysis (DESeq2)
5847 KOs
> 6/12 Host-Inoculum Combinations
1478 KOs
12/12 Host-inoculum Combinations
266 KOs
203 KOs
Existing pathway annotation
Relative abundance
Function A presence/absence
0.40 0.10 0
0.30 0.00 0
0.15 0.20 0
0.05 0.50 1
0.10 0.20 1
0.10 + 0.00 + 0.20 / 0.40 + 0.30 + 0.15 = 0.35
0.50+0.20 / 0.05+0.10 = 4.7
Function A FC Absent
Function A FC Present / FC Absent
log2(4.7/0.35)
Function A FC Present
d
Barley
Arabidopsis
Lotus
e
Theoretical abundance (%)
> 0
> 10
> 20
> 30
> 40
> 50
Category
Amino acid metabolism
Basics
Biosynthesis of other secondary metabolites
Carbohydrate metabolism
Cell growth and death
Cell motility
Cellular community − prokaryotes
Energy metabolism
Environmental adaptation
Glycan biosynthesis and metabolism
Membrane transport
Metabolism of cofactors and vitamins
Metabolism of other amino acids
Nucleotide metabolism
Signal transduction
Transcription
Xenobiotics biodegradation and metabolism
log2( FC Present / FC Absent )
Relative abundance
f
Present
Absent
AtSC
HvSC
LjSC
SSC
A: Secretion (n% = 1.0) (p=4.3e−05)
B: Vitamin B6 metabolism (n% = 5.4) (p=4.3e−05)
C: Cysteine and methionine metabolism (n% = 9.0) (p=4.3e−05)
D: Methane metabolism (n% = 1.4) (p=4.3e−05)
E: Quorum sensing (n% = 13.3) (p=4.3e−05)
F: Exopolysaccharide biosynthesis (n% = 8.8) (p=4.3e−05)
G: Flagellar assembly (n% = 16.2) (p=4.3e−05)
H: Aminobenzoate degradation (n% = 4.3) (p=4.3e−05)
I: Glycerophospholipid metabolism (n% = 19.4) (p=4.3e−05)
J: Membrane protein (n% = 18.7) (p=4.3e−05)
K: Folate biosynthesis (n% = 12.2) (p=4.3e−05)
L: Oxidative phosphorylation (n% = 20.4) (p=4.3e−05)
M: Phenylalanine, tyrosine and tryptophan biosynthesis (n% = 26.1) (p=4.3e−05)
N: Ascorbate and aldarate metabolism (n% = 13.8) (p=4.3e−05)
O: Transcriptional regulator (n% = 21.4) (p=4.3e−05)
P: Pantothenate and CoA biosynthesis (n% = 21.3) (p=4.3e−05)
Q: Arginine biosynthesis (n% = 28.8) (p=4.3e−05)
R: Transporter (n% = 27.0) (p=4.3e−05)
S: ABC transporters (n% = 25.9) (p=4.3e−05)
T: Lipopolysaccharide biosynthesis (n% = 24.5) (p=4.3e−05)
U: Ubiquinone and other terpenoid−quinone biosynthesis (n% = 30.7) (p=4.3e−05)
V: Lysine degradation (n% = 31.1) (p=4.3e−05)
W: Cell cycle − Caulobacter (n% = 26.7) (p=4.3e−05)
X: Other glycan degradation (n% = 33.1) (p=4.3e−05)
Y: Cyanoamino acid metabolism (n% = 48.5) (p=4.3e−05)
log₂(FC) Root vs Input
g
Distribution of 266 KOs in culture collections
Density
AtSC
HvSC
LjSC
Number of KOs
h
Distribution of 266 KOs in Levy et al. (2018)
Levy et al. (2018)
SSC
3331
177
51
Density
Non Plant-Associated
Soil
Plant-Associated
Number of KOs

**Figure S23. Core functions enriched in root microbiota in the absence of dominant strains.** (a) Overlap in differentially abundant bacterial KEGG Orthologs (KOs) enriched in the roots of Arabidopsis, Barley, or Lotus compared to the initial inoculum across four different inocula in the dataset in which the Lotus symbionts and *Rhizobacter* sp. P2_G4 were computationally excluded. (b) Functional overlap between hosts per inoculum, shown by the number of differentially abundant KOs in the dataset in which the Lotus symbionts and *Rhizobacter* sp. P2_G4 were computationally excluded. (c) Sankey diagram (top) of KO enrichment and filtering calculation with a stricter selection as compared to figure 4, from 5847 root enriched bacterial KOs, to 266 common host-enriched KOs, and to 203 pathway-annotated functions. Function enrichment index method (bottom), calculated as a fold change ratio based on change in cumulative relative abundance of bacterial populations with or without the pathway between root and initial inoculum. (d) Density distribution of common host-enriched KOs (n=266) across the three hosts based on the functional enrichment index (c). (e) Pathways selected by the host based on significant KOs (from the DESeq2 analyses) reflected in the cumulative relative abundance of the isolates encoding these (X-axis) and the function enrichment index combined across the three hosts (Y-axis). Best pathways have been selected using an exponential decay function maximizing function enrichment index and the cumulative relative abundance of the pathways. Pinpointed pathways are tagged by letters from highest to lowest function enrichment index. The coordinate sizes indicate the proportion of isolates that have the respective pathway in the SSC condition and the color indicates the category in which the KOs are included. (f) Boxplots of Log2(FC) values of root versus initial inoculum reference for bacteria community members with or without the given pathway. (g) Density plots showing the distribution of the 266 common host-enriched KOs in the three culture collections used in this study. (h) Density plots showing the distribution of the 266 common host-enriched KOs in the bacterial genome database categorized as soil, non-plant associated or plant-associated bacteria (Levy et al., 2018). The Venn diagram displays the overlap between the identified plant-associated KO functions in this study and Levy et al. (2018).

#### Supplemental Results 3 – Host-specific functions

When examining host-dependent bacterial functions, we found that the number of Lotus-specific bacterial genes (355) far exceeds that of Arabidopsis-specific (48) and Barley-specific (40) bacterial genes (Figures 4b, 4c, 5a, Tables S8, S9, and S10). This suggests that Lotus exerts a stronger selection pressure compared to the other two hosts. To identify the most relevant host-specific bacterial functions, we focused on functions from gene cassettes where most, if not all, genes consistently displayed host-specificity in our analysis.

*Arabidopsis-specific bacterial genetic features - bacteriochlorophyllides*

We observed that genes within the photosynthetic bacteriochlorophyllide biosynthesis gene cluster (*bch*) were enriched specifically in Arabidopsis-associated communities (Figure S24 and Table S9). Notably, 11 out of 18 *bch* genes showed Arabidopsis-specific enrichment (Figure S24a), especially when inoculated with the SSC (Figure S24b). This finding is surprising because bacteriochlorophyllide is involved in harnessing light energy for photosynthesis, yet light penetrates only about 1 cm into the soil, reaching only a small fraction of the Arabidopsis root (Wu *et al.*, 2014). However, a positive correlation between soil moisture and light penetration depth has been observed (Williams *et al.*, 2016). This correlation is consistent with the fact that the Arabidopsis plants were kept covered during the experiment, likely affecting soil moisture. Bacteria encoding this photosynthetic gene cluster mostly derive from the *Flavobacteriaceae, Pseudomonadaceae*, and *Burkholderiaceae* families (Figure S24c), representing common colonizers of Arabidopsis roots (Pieterse *et al.*, 2021; Thiergart *et al.*, 2019; Wang *et al.*, 2021). We observed that the abundance of two *bch* genes, *bchE* and *bchU*, was markedly larger for Lotus. BchE has been reported to be mainly active in strictly anaerobic photosynthetic bacteria (Ouchane *et al.*, 2004), while bchU converts chlorophyllide *a* to chlorophyll, a more efficient chlorophyllide due to its ability to harvest energy from a greater bandwidth of light (Malina *et al.*, 2021). This suggests that photosynthetic traits are Arabidopsis-specific, though photosynthetic isolates that are strictly anaerobic and have more efficient light-harvesting abilities could be Lotus-specific. Alternatively, imprecise, or wrongful annotation of *bchE* homologues could be responsible for this observation, and its high abundance likely underlies the unexpected Lotus, as opposed to Arabidopsis, specificity when isolate abundances are taken into consideration (Figure S24d).

*Barley-specific bacterial genetic features – exopolysaccharide succinoglycans*

Among the Barley-specific genes, multiple genes can be found that constitute thiamine and inositol transporters, as well as two succinoglycan biosynthesis proteins *exoV* and *exoZ* mainly when inoculated with the AtSC (Figures S25a, b, and c, and Table S10). These two *exo* genes derive mainly from *Rhizobiaceae* isolates (Figure S25d). Succinoglycans are extracellular structures associated with the bacterial cell wall, where low molecular weight succinoglycans, in particular, play a role in nodule infection (Maillet *et al.*, 2020; Mendis *et al.*, 2016; Skorupska *et al.*, 2006). ExoV and exoZ add pyruvate and acetyl groups to the end of the polysaccharide chain (Figure S25a), therewith stabilizing the succinoglycans, and leading to a higher proportion of high molecular weight succinoglycans (Jeong *et al.*, 2022; York & Walker, 1998). The high relative abundance of both genes in Barley-associated communities indicates that these high molecular weight succinoglycans could be important for Barley root colonization (Figure S25a).

*Lotus-specific bacterial genetic features – symbiosis-related*

Lotus shows a substantially higher amount of enriched host-specific genes as compared to Arabidopsis or Barley (Table S11). Nonetheless, in the absence of symbionts, only a fraction of those remains (Table S12) (61 out of the 355). In the dataset including symbionts, Lotus shows, as predicted, an enrichment of *nif* and *nod* genes in its root microbiome, which are involved in nitrogen fixation and nodulation, respectively (Göttfert, 1993; Spaink *et al.*, 2012) (Figure S26, S27 and Table S11). The *nif* and *nod* gene clusters are entirely Lotus-specific (Koirala & Brözel, 2021), except for *nifV* (Arabidopsis-specific), *nifH* in the AtSC (Barley-specific), and *nodU* (not host-specific) (Figure S26 and S27). The *nifV* gene encodes an homocitrate synthase required for nitrogen fixation. However, most legume nitrogen-fixing symbionts such as *Mesorhizobium* spp. lack this gene, and homocitrate is provided by the host legume via FEN1 (Hakoyama *et al.*, 2009). Enrichment of *nifV* in Arabidopsis roots indicates ta possible selection for plant growth-promoting diazotrophic rhizobacteria that are capable of fixing atmospheric nitrogen to the benefit of the plant, a phenomenon that was previously described by Santos *et al.* (2017). It is notable, however, that Barley most strongly selects for nitrogen-fixation genes even if it does not have the natural capacity for symbiosis with nitrogen-fixing bacteria. This also includes Barley inoculated with the AtSC, where we also observed that nodulation genes are highly Barley-specific (Figure S26c), suggesting that Barley might be a strong selector of bacteria that make nitrogen available to the plant.

One of the most enriched Lotus-specific gene clusters aside from the *nif* and *nod* genes are the *ery* genes that are involved in the transport and catabolism of the sugar erythritol (Table S11) (Figure S28) (Geddes *et al.*, 2010). While erythritol is exuded by many plants (Carvalhais *et al.*, 2015; Vranova *et al.*, 2013), its exudation by Lotus nodules (Colebatch *et al.*, 2004; Ranner *et al.*, 2023) suggests that bacterial erythritol catabolism might be uniquely associated with Lotus nodules. This process could serve as another signal for symbionts to be enriched in the Lotus root microbiome.

#### *References*

Carvalhais, L. C., Dennis, P. G., Badri, D. V., Kidd, B. N., Vivanco, J. M., & Schenk, P. M. (2015). Linking jasmonic acid signaling, root exudates, and rhizosphere microbiomes. *Molecular Plant-Microbe Interactions*, *28*(9), 1049–1058.

Colebatch, G., Desbrosses, G., Ott, T., Krusell, L., Montanari, O., Kloska, S., Kopka, J., & Udvardi, M. K. (2004). Global changes in transcription orchestrate metabolic differentiation during symbiotic nitrogen fixation in *Lotus japonicus*. *The Plant Journal*, *39*(4), 487–512.

Geddes, B. A., Pickering, B. S., Poysti, N. J., Collins, H., Yudistira, H., & Oresnik, I. J. (2010). A locus necessary for the transport and catabolism of erythritol in *Sinorhizobium meliloti*. *Microbiology*, *156*(10), 2970–2981.

Göttfert, M. (1993). Regulation and function of rhizobial nodulation genes. *FEMS Microbiology Letters*, *104*(1–2), 39–63.

Hakoyama, T., Niimi, K., Watanabe, H., Tabata, R., Matsubara, J., Sato, S., Nakamura, Y., Tabata, S., Jichun, L., Matsumoto, T., Tatsumi, K., Nomura, M., Tajima, S., Ishizaka, M., Yano, K., Imaizumi-Anraku, H., Kawaguchi, M., Kouchi, H., & Suganuma, N. (2009). Host plant genome overcomes the lack of a bacterial gene for symbiotic nitrogen fixation. Nature, 462(7272), 514–517.

Jeong, J., Kim, Y., Hu, Y., & Jung, S. (2022). Bacterial succinoglycans: structure, physical properties, and applications. *Polymers*, *14*(2), 276.

Koirala, A., & Brözel, V. S. (2021). Phylogeny of nitrogenase structural and assembly components reveals new insights into the origin and distribution of nitrogen fixation across bacteria and archaea. *Microorganisms*, *9*(8), 1662.

Maillet, F., Fournier, J., Mendis, H. C., Tadege, M., Wen, J., Ratet, P., Mysore, K. S., Gough, C., & Jones, K. M. (2020). *Sinorhizobium meliloti* succinylated high-molecular-weight succinoglycan and the *Medicago truncatula* LysM receptor-like kinase MtLYK10 participate independently in symbiotic infection. *The Plant Journal*, *102*(2), 311–326.

Malina, T., Koehorst, R., Bína, D., Pšenčík, J., & van Amerongen, H. (2021). Superradiance of bacteriochlorophyll c aggregates in chlorosomes of green photosynthetic bacteria. *Scientific Reports*, *11*(1), 8354.

Mendis, H. C., Madzima, T. F., Queiroux, C., & Jones, K. M. (2016). Function of succinoglycan polysaccharide in *Sinorhizobium meliloti* host plant invasion depends on succinylation, not molecular weight. *MBio*, *7*(3).

Ouchane, S., Steunou, A.-S., Picaud, M., & Astier, C. (2004). Aerobic and anaerobic Mg-protoporphyrin monomethyl ester cyclases in purple bacteria. *Journal of Biological Chemistry*, *279*(8), 6385–6394.

Pieterse, C. M. J., Berendsen, R. L., de Jonge, R., Stringlis, I. A., Van Dijken, A. J. H., Van Pelt, J. A., Van Wees, S. C. M., Yu, K., Zamioudis, C., & Bakker, P. A. H. M. (2021). *Pseudomonas simiae* WCS417: star track of a model beneficial rhizobacterium. *Plant and Soil*, *461*(1–2), 245–263.

Ranner, J. L., Schalk, S., Martyniak, C., Parniske, M., Gutjahr, C., Stark, T. D., & Dawid, C. (2023). Primary and secondary metabolites in *Lotus japonicus*. *Journal of Agricultural and Food Chemistry*, *71*(30), 11277–11303.

Santos, A. R. S., Etto, R. M., Furmam, R. W., Freitas, D. L. de, Santos, K. F. d'Eça N., Souza, E. M. de, Pedrosa, F. de O., Ayub, R. A., Steffens, M. B. R., & Galvão, C. W. (2017). Labeled *Azospirillum brasilense* wild type and excretion-ammonium strains in association with barley roots. *Plant Physiology and Biochemistry*, *118*, 422–426.

Skorupska, A., Janczarek, M., Marczak, M., Mazur, A., & Król, J. (2006). Rhizobial exopolysaccharides: genetic control and symbiotic functions. *Microbial Cell Factories*, *5*(1), 7.

Spaink, H. P., Kondorosi, A., & Hooykaas, P. J. J. (2012). *The Rhizobiaceae: molecular biology of model plant-associated bacteria*. Springer Science & Business Media.

Thiergart, T., Durán, P., Ellis, T., Vannier, N., Garrido-Oter, R., Kemen, E., Roux, F., Alonso-Blanco, C., Ågren, J., Schulze-Lefert, P., & Hacquard, S. (2019). Root microbiota assembly and adaptive differentiation among European *Arabidopsis* populations. *Nature Ecology & Evolution*, *4*(1), 122–131.

Vranova, V., Rejsek, K., Skene, K. R., Janous, D., & Formanek, P. (2013). Methods of collection of plant root exudates in relation to plant metabolism and purpose: a review. *Journal of Plant Nutrition and Soil Science*, *176*(2), 175–199.

Wang, N. R., Melnyk, R. A., Wiesmann, C. L., Hossain, S. S., Chi, M.-H., Martens, K., & Haney, C. H. (2021). Commensal *Pseudomonas fluorescens* protect *Arabidopsis* from closely-related *Pseudomonas* pathogens in a colonization-dependent manner. *BioRxiv*, 2021–2029.

Williams, E. A., Cobbina, S. M., & Bentil, J. (2016). The effects of light intensity on soil depth of different moisture contents using laser sensor. *International Journal of Scientific and Research Publications*, *6*(5), 488–493.

Wu, X., Ge, T., Yuan, H., Li, B., Zhu, H., Zhou, P., Sui, F., O'Donnell, A. G., & Wu, J. (2014). Changes in bacterial CO2 fixation with depth in agricultural soils. *Applied Microbiology and Biotechnology*, *98*(5), 2309–2319.

York, G. M., & Walker, G. C. (1998). The succinyl and acetyl modifications of succinoglycan influence susceptibility of succinoglycan to cleavage by the *Rhizobium meliloti* glycanases ExoK and ExsH. *Journal of Bacteriology*, *180*(16), 4184–4191.

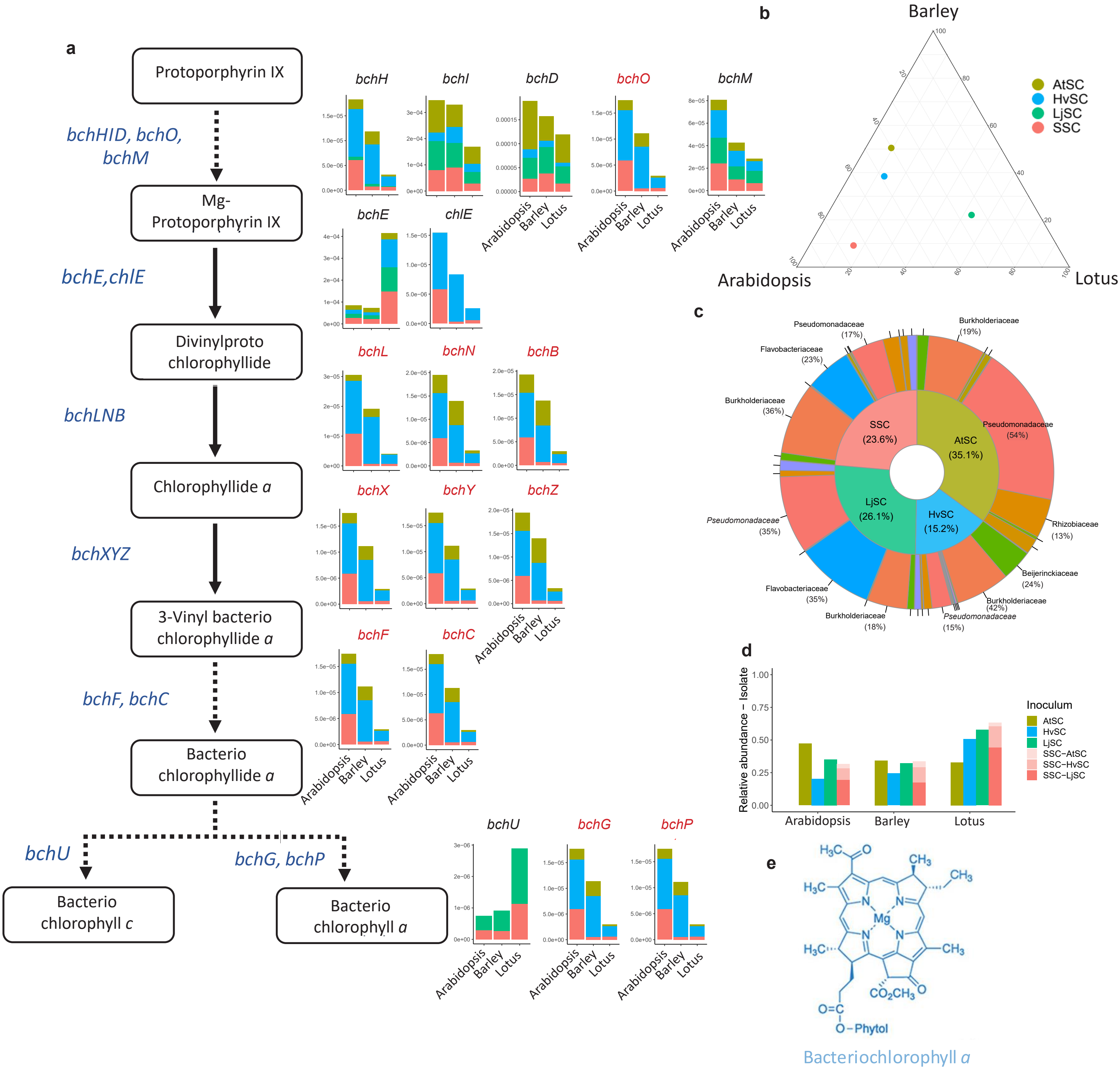

**Figure S24. Arabidopsis-specific genes *bch/chl*.** (a) Bacterial *bch/chl* genes in the bacteriochlorophyllide biosynthesis pathway with gene abundances in every host and inoculum. (b) Host-specificity of the *bch/chl* genes in each inoculum regarding the fold change of isolates from initial inoculum to root. (c) Piedonut plot illustrating which bacterial families are most responsible for the cumulative relative abundance of bacteria with *bch/chl* genes in each inoculum and how these cumulative relative abundances compare to each other across inocula. (d) Abundances of isolates in each host and inoculum that have at least one of the displayed *bch/chl* genes. In the SSC it shows the origin of the isolates that contribute to this cumulative relative abundance. (e) Structure formula of bacteriochlorophyll.

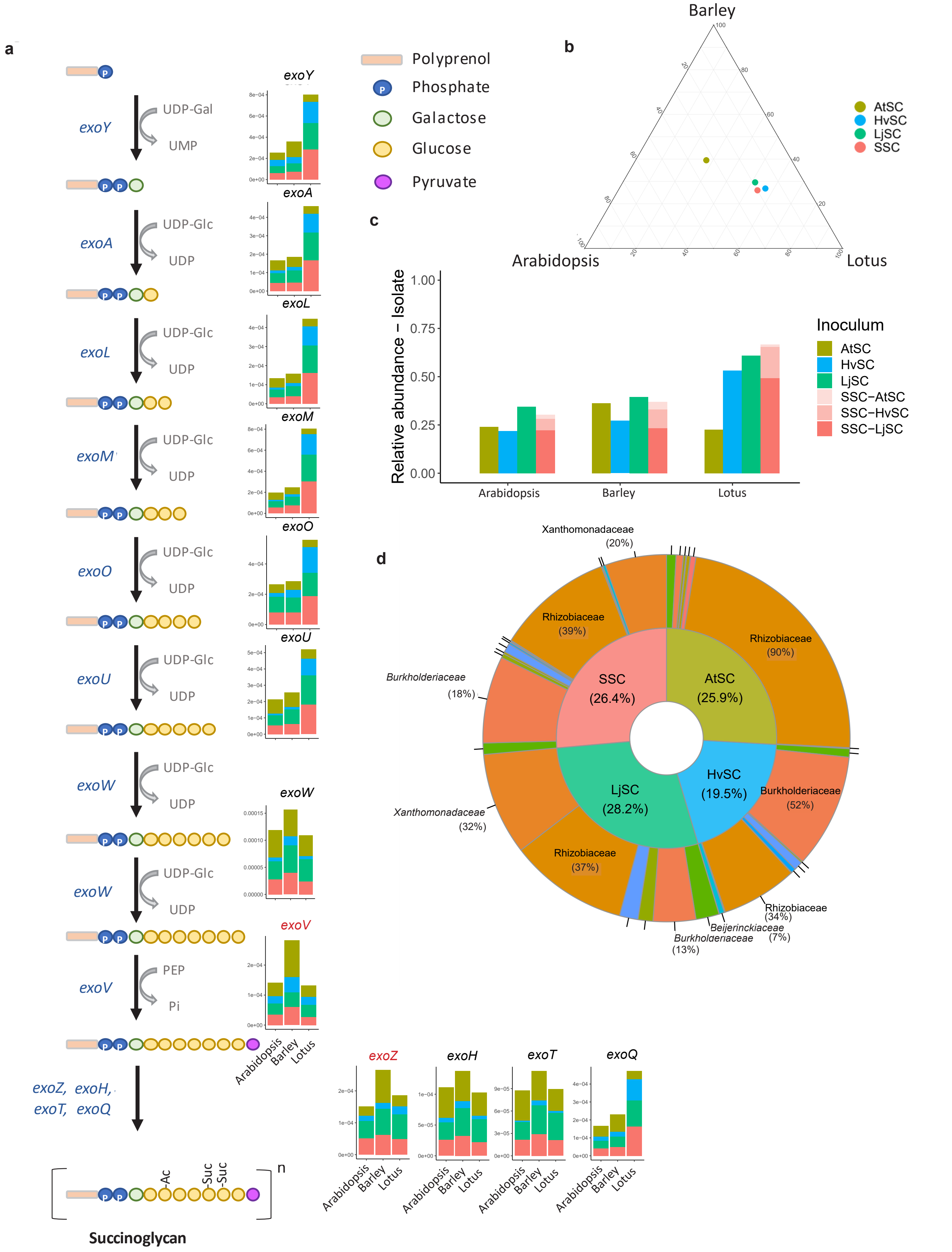

**Figure S25. Barley-specific genes *exoV* and *exoZ*.** (a) Bacterial *exo* genes in the succinoglycan biosynthesis pathway with gene abundances in every host and inoculum. (b) Host-specificity of the *exo* genes in each inoculum regarding the fold change of isolates from initial inoculum to root. (c) Abundances of isolates in each host and inoculum that have *exo* genes. In the SSC it shows the origin of the isolates that contribute to this cumulative relative abundance (d) Piedonut plot illustrating which bacterial families are most responsible for the cumulative relative abundance of bacteria with *exo* genes in each inoculum and how these cumulative relative abundances compare to each other across inocula.

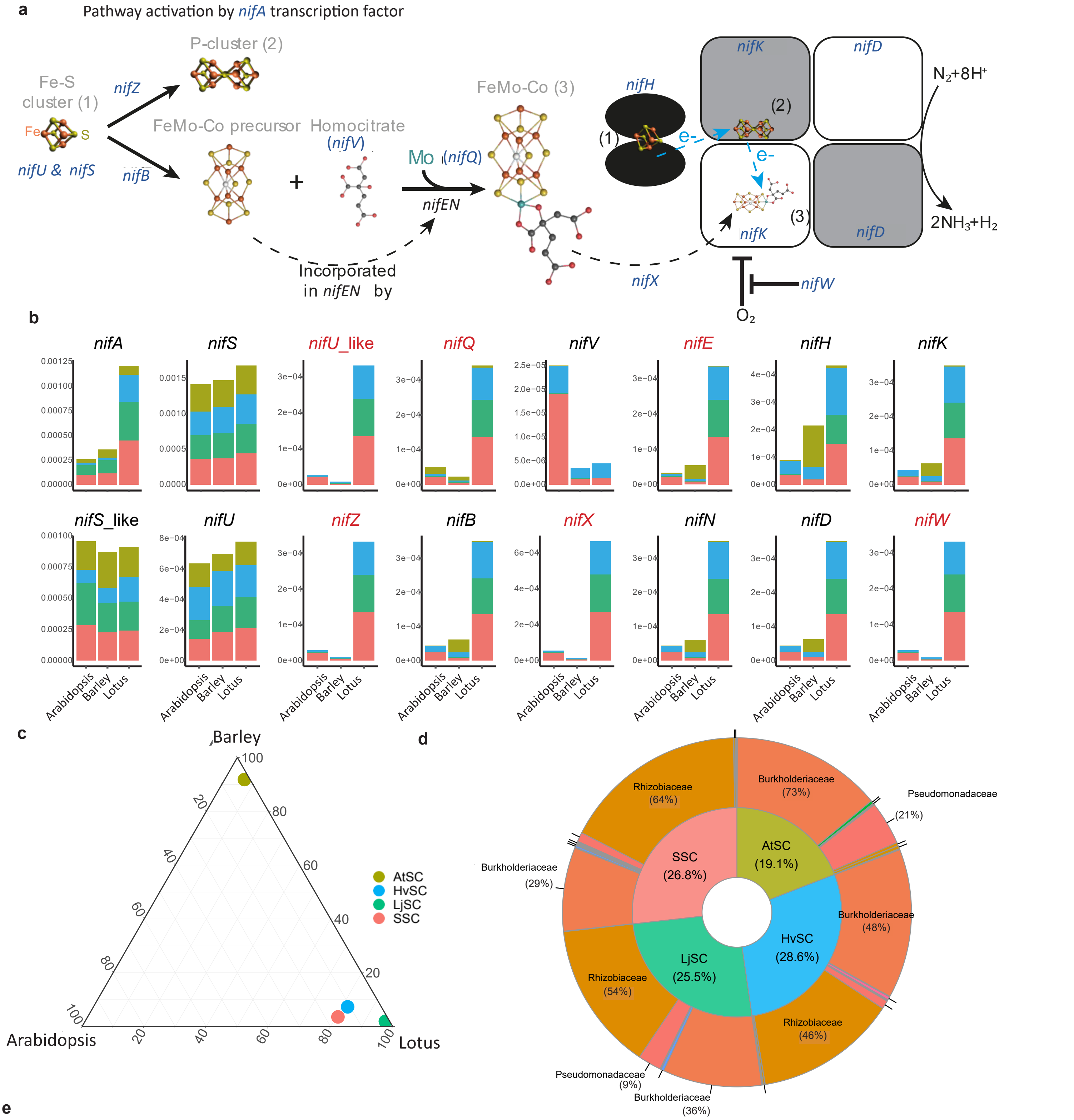

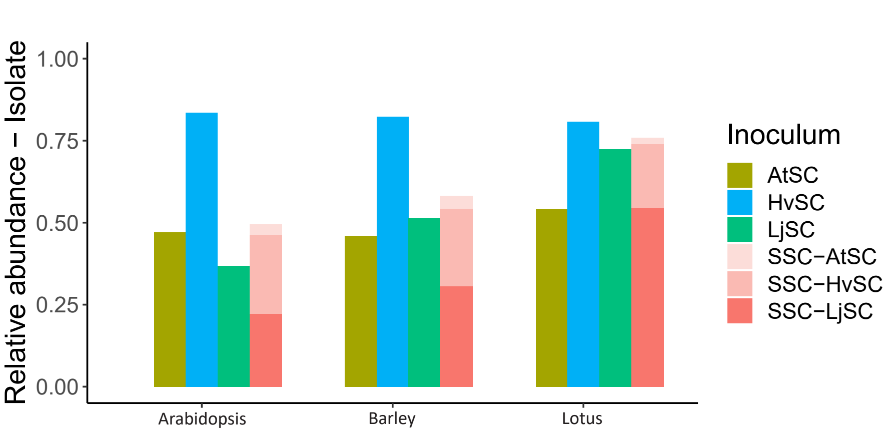

**Figure S26. Lotus-specific nitrogen fixation genes *nif*.** (a) Bacterial *nif* genes in the nitrogen fixation pathway. (b) *nif* gene abundances in every host and inoculum. (c) Host-specificity of the *nif* genes in each inoculum regarding the fold change of isolates from initial inoculum to root. (d) Piedonut plot illustrating which bacterial families are most responsible for the cumulative relative abundance of bacteria with *nif* genes in each inoculum and how these cumulative relative abundances compare to each other across inocula. (e) Abundances of isolates in each host and inoculum that have *nif* genes. In the SSC it shows the origin of the isolates that contribute to this cumulative relative abundance.

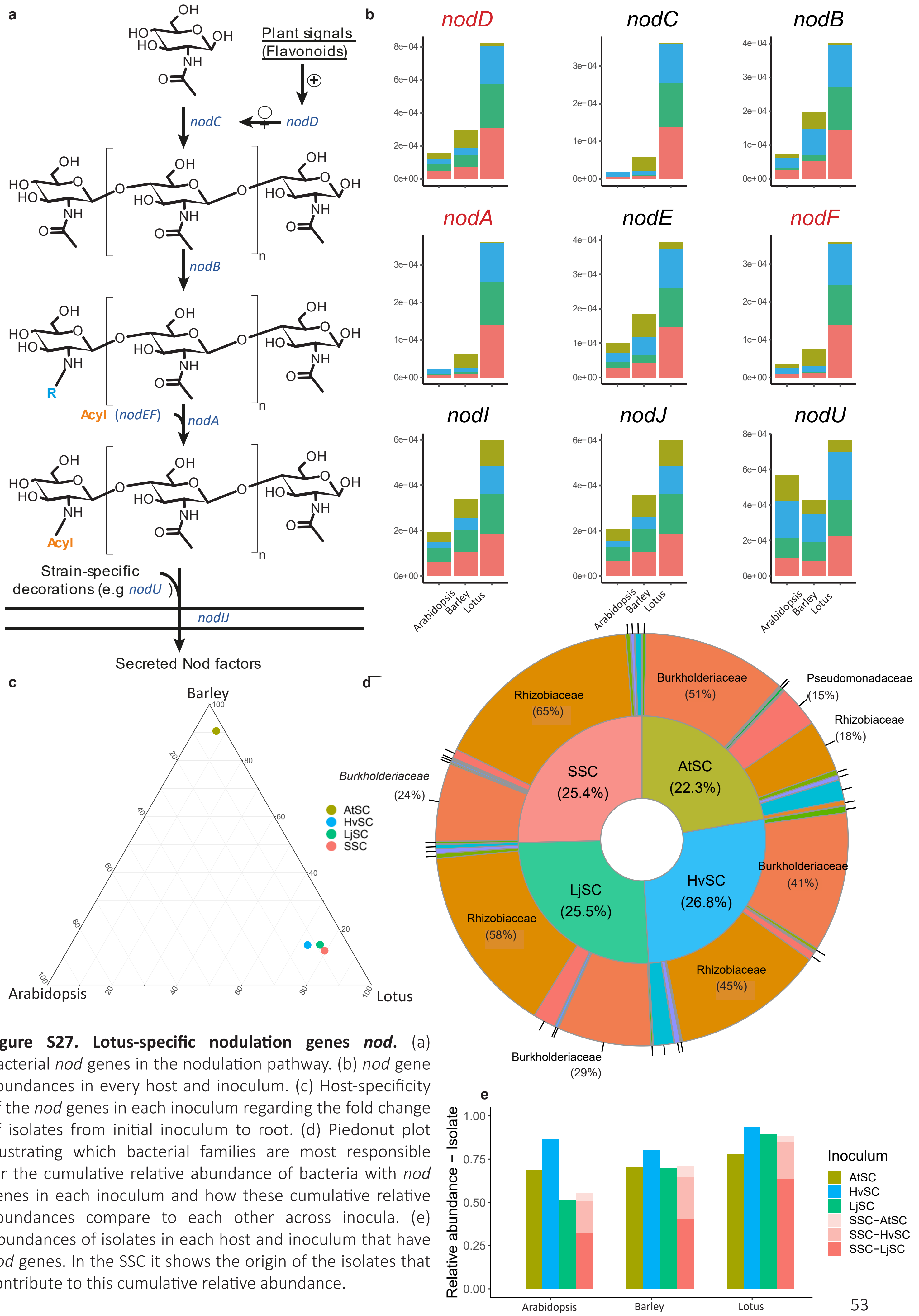

**Figure S27. Lotus-specific nodulation genes *nod*.** (a) Bacterial *nod* genes in the nodulation pathway. (b) *nod* gene abundances in every host and inoculum. (c) Host-specificity of the *nod* genes in each inoculum regarding the fold change of isolates from initial inoculum to root. (d) Piedonut plot illustrating which bacterial families are most responsible for the cumulative relative abundance of bacteria with *nod* genes in each inoculum and how these cumulative relative abundances compare to each other across inocula. (e) Abundances of isolates in each host and inoculum that have *nod* genes. In the SSC it shows the origin of the isolates that contribute to this cumulative relative abundance.

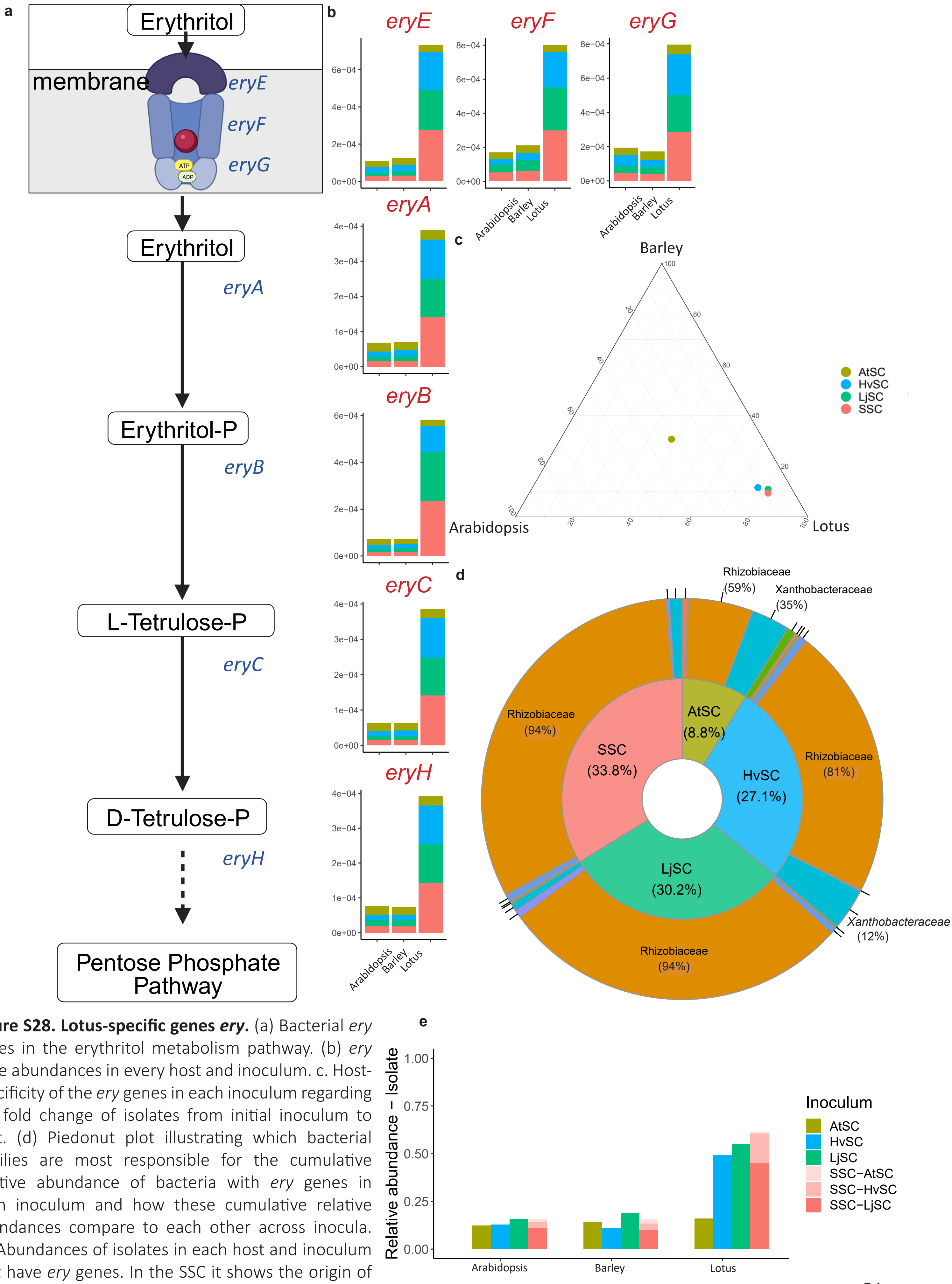

**Figure S28. Lotus-specific genes *ery*.** (a) Bacterial *ery* genes in the erythritol metabolism pathway. (b) *ery* gene abundances in every host and inoculum. c. Host-specificity of the *ery* genes in each inoculum regarding the fold change of isolates from initial inoculum to root. (d) Piedonut plot illustrating which bacterial families are most responsible for the cumulative relative abundance of bacteria with *ery* genes in each inoculum and how these cumulative relative abundances compare to each other across inocula. (e) Abundances of isolates in each host and inoculum that have *ery* genes. In the SSC it shows the origin of the isolates that contribute to this cumulative relative abundance.

#### Supplemental results 4 – Host-specific functions - ABC transporters

To investigate general trends in host-specificity, we grouped KOs into genes by compiling all KO gene cassettes into gene clusters based on their shared gene-level KEGG annotation (Supplemental Data S21) and assessed the host-specificity of these clusters in the nine pathways that are most frequently host-specific (Figures 5c and S29). This analysis could potentially provide additional information about host-specificity at the gene/gene cluster level beyond the broader pathway level. We observe that host-specificity can differ substantially at the gene/gene cluster level depending on the respective inoculum (Figure S29 and Table S13). One such example is represented by the host-specificity of *ssp* genes, an operon that plays a role in quorum sensing (Rice *et al.*, 2001). We identified this operon as Arabidopsis-specific with LjSC and SSC, Barley-specific with AtSC and Lotus-specific when inoculated with HvSC (Figure S29). The nonconformist pattern of specificity displayed by these genes may suggest a complex interaction between host and bacterium. Alternatively, these genes could be genetic hitchhikers that are coincidentally present in the same bacterial genome as genes that enhance a bacterium's root competence.

*Diversity of ABC transporters and root competency*

The analysis of differential abundance of KOs between inocula and root communities revealed that ABC transporters emerged as a category with the largest number of host-specific bacterial functions (Figures 5c, S29 and Table S13). To better understand the role and contribution of ABC transporters in root microbiota assembly, we first investigated the number and diversity of ABC transporters in the three host-specific inocula (Figure S30). Analysis at the bacterial isolate level revealed that LjSC encompassed isolates with the largest number of ABC transporters, followed by HvSC and AtSC (Figure S30, solid lines). All three hosts enriched their root microbiota for isolates that encode a broad spectrum of ABC transporter illustrated by the many top colonizers encoding a relatively large number of ABC transporters. This included the so-called dominator strains, i.e., the Lotus symbionts and the *Rhizobacter* sp. P2_G4, indicating that ABC transporter diversity might contribute to their observed dominance (Figure S30). Root communities assembled by the three hosts in the presence of LjSC, however, were dissimilar, revealing little to no correlation—or even a negative correlation—between the number of ABC transporters and the relative abundance of isolates (Figure S30). We suggest that this is likely due to LjSC containing a higher overall number of ABC transporters compared to the other SynComs, which raises the baseline and diminishes the impact of additional transporters.

Next, we investigated if the number and the diversity of the ABC transporters encoded by specific isolates, when compared to the remaining isolates within the same family, can explain the abundance in the root communities. We followed a similar strategy used for analyzing functional diversity in bacterial families as illustrated in Figures 3d and 3e (Figure S31). This analysis revealed a complex pattern. We observed an overall positive correlation between an isolates' number of ABC transporters and its relative abundance for AtSC or HvSC (Figure S31), despite not being always significant. For LjSC and SSC, in turn, we observed opposite patterns. A negative correlation in the LjSC (Figure S31) could relate to the high number of ABC transporters in LjSC isolates (Figure S30), suggesting that competition, in terms of relative abundance in the root microbiome, does not depend on ABC transporter diversity when the average ABC transporter diversity in the community is already high. Bacterial isolates belonging to the *Rhizobiaceae*, *Pseudomonadaceae*, and *Burkholderiaceae* families display the highest diversity in ABC transporters (Figure S32). These are also the largest families in the SSC, both in terms of numbers and abundances in the root communities (Figure 3d). Altogether, this suggests that bacteria that can import/export a larger diversity of compounds are more likely to successfully colonize the plant root. Additionally, these bacteria could play a crucial role in the microbial community by fostering cooperation, such as producing and exchanging amino acids, vitamins, or other compounds with other community members (Mataigne *et al.*, 2021). Interestingly, the *Flavobacteriaceae* display a negative correlation with ABC transporter diversity (Figure S31), suggesting that these bacterial isolates have a different strategy to colonize the roots. Multiple studies indicated how *Flavobacteriaceae* are used as biocontrol agents by negatively affecting fungal and oomycetal pathogens (Carrión *et al.*, 2019; Sang *et al.*, 2008), indicating that other traits in the *Flavobacteriaceae,* possibly related to microbe-microbe warfare, might enhance their root competence.

*References*

Carrión, V. J., Perez-Jaramillo, J., Cordovez, V., Tracanna, V., de Hollander, M., Ruiz-Buck, D., Mendes, L. W., van Ijcken, W. F. J., Gomez-Exposito, R., Elsayed, S. S., Mohanraju, P., Arifah, A., van der Oost, J., Paulson, J. N., Mendes, R., van Wezel, G. P., Medema, M. H., & Raaijmakers, J. M. (2019). Pathogen-induced activation of disease-suppressive functions in the endophytic root microbiome. *Science*, *366*(6465), 606–612.

Mataigne, V., Vannier, N., Vandenkoornhuyse, P., & Hacquard, S. (2021). Microbial systems ecology to understand cross-feeding in microbiomes. *Frontiers in Microbiology*, *12*.

Rice, K., Peralta, R., Bast, D., de Azavedo, J., & McGavin, M. J. (2001). Description of *Staphylococcus* serine protease (ssp) operon in *Staphylococcus aureus* and nonpolar inactivation of *sspA* -encoded serine protease. *Infection and Immunity*, *69*(1), 159–169.

Sang, M. K., Chun, S.-C., & Kim, K. D. (2008). Biological control of Phytophthora blight of pepper by antagonistic rhizobacteria selected from a sequential screening procedure. *Biological Control*, *46*(3), 424–433.

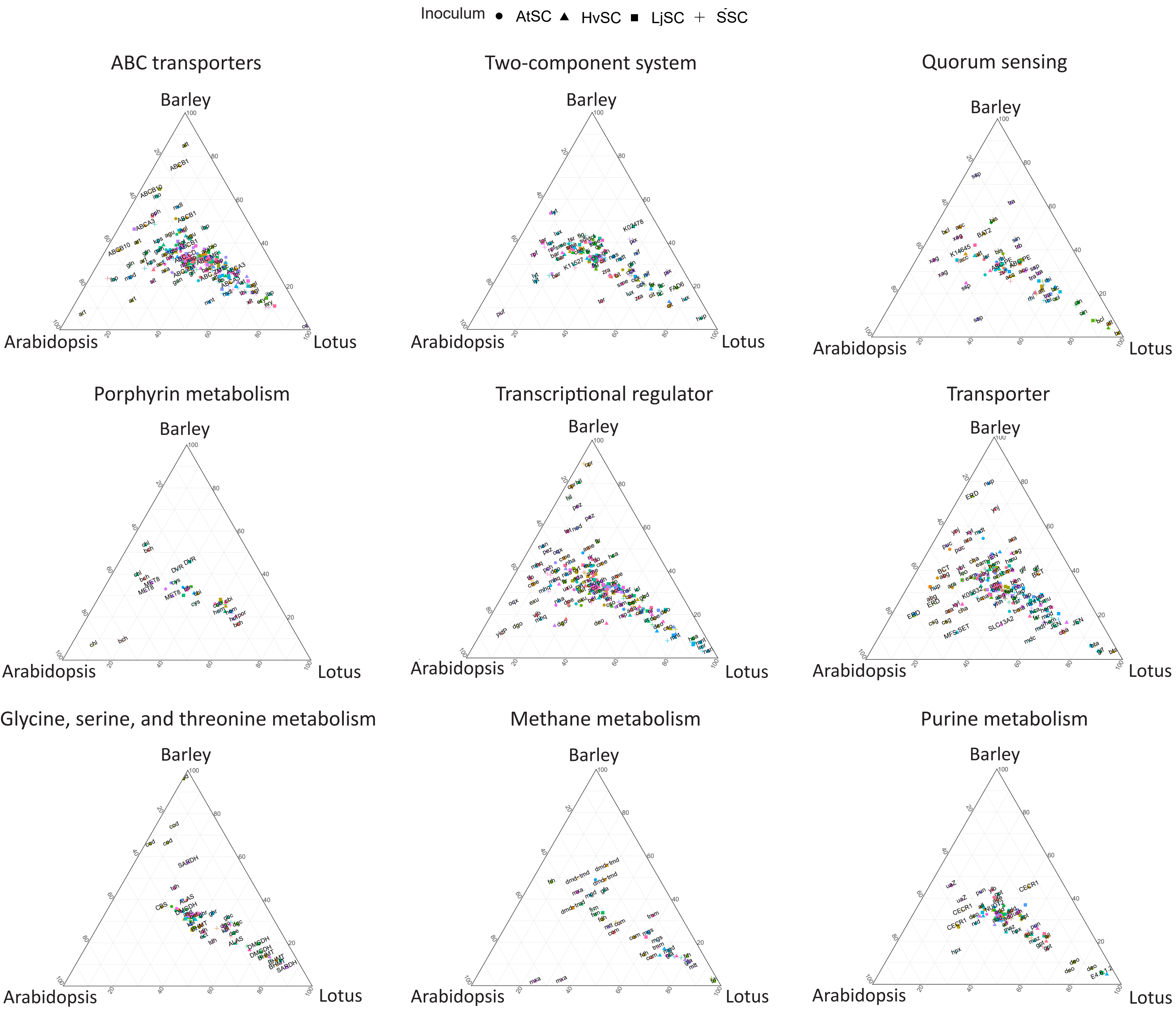

**Figure S29. Host-specificity of genes from the top nine most frequent host-specific pathways.** Ternary plots that indicate host-specificity of genes for the nine pathways with the most host-specific genes. The coordinates position is the result of three values, one per host, in which the value indicates the fold change of isolates with the gene from the initial inoculum to the root microbiome. Each gene has four coordinates that represent the inocula (indicated by shapes).

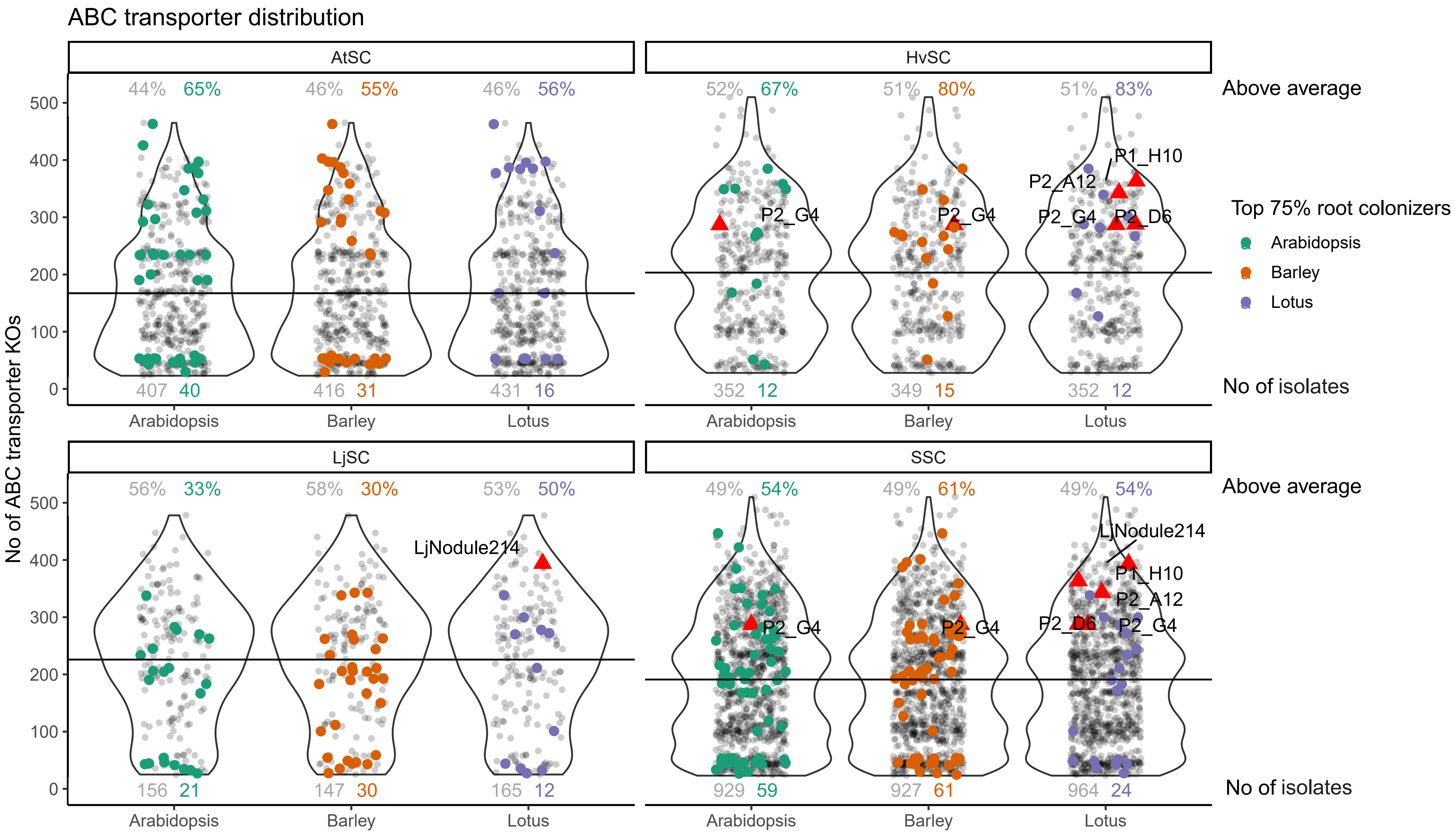

**Figure S30. Number of unique ABC transporters in top root colonizers.** For each inoculum and host, the number of ABC transporters of the bacterial isolates is shown on the Y-axis. The colored coordinates indicate the bacterial isolates that are among the most abundant isolates (top 75%) on the roots of each host and inoculum. The solid line indicates the inoculum's average number of ABC transporters, while the colored numbers on the top indicate the percentage of the most abundant isolates that are above average. The colored numbers at the bottom indicate the number of isolates that are in the top 75% most abundant isolates. The grey-colored numbers indicate those percentages and numbers of isolates that are not among the top 75%. The red triangles indicate the dominant isolates in the dataset; the Lotus symbionts (LjNodule214, P1_H10, P2_A12, and P2_D6) and the dominant HvSC *Rhizobacter* sp. P2_G4.

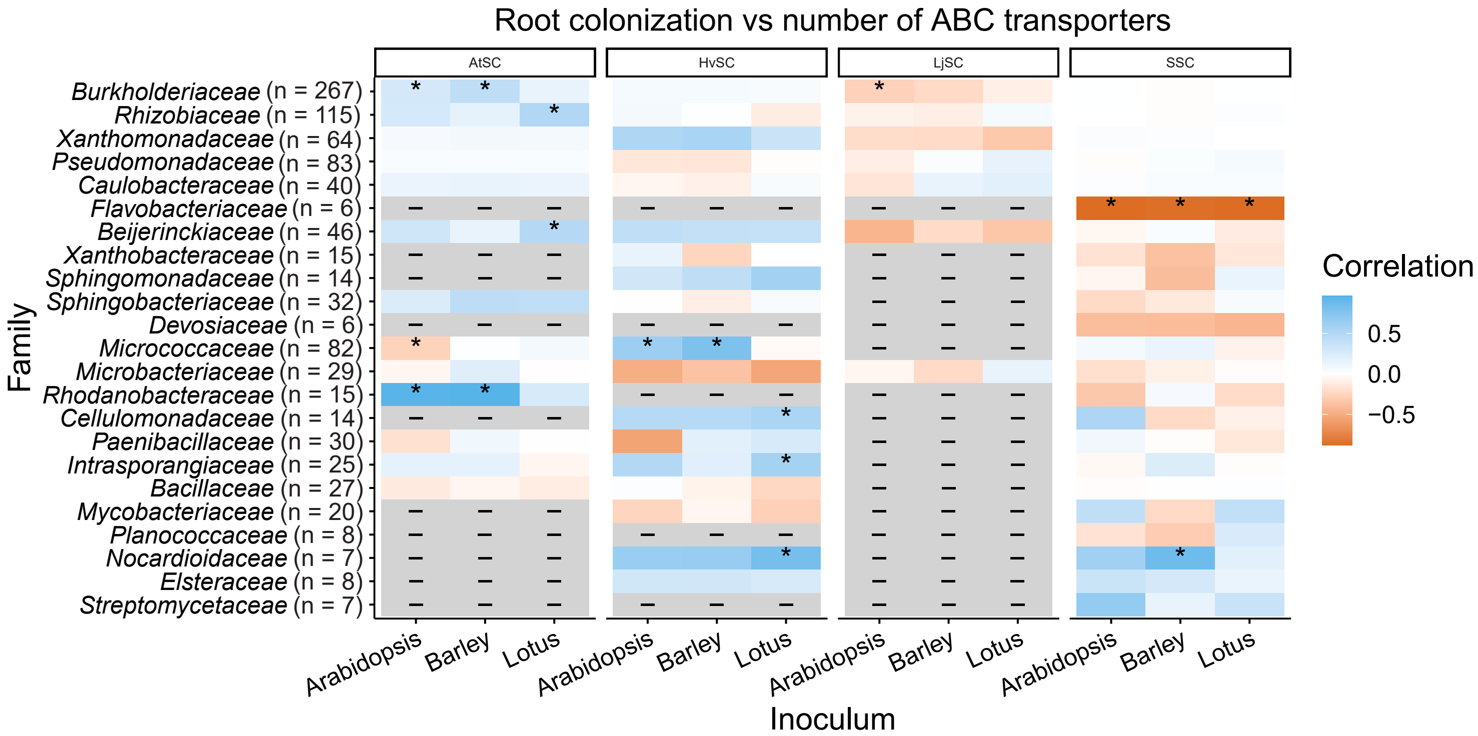

**Figure S31. Correlation heatmap between ABC transporter diversity and abundance.** For each isolate, the number of ABC transporters is correlated with its abundance on the root of different hosts. The resulting $R^2$ correlation value is indicated in the heatmap. A significant correlation is indicated by '*' (p-value < 0.05). Grey-colored cells indicate that there were not enough root colonizers in that family to create a correlation.

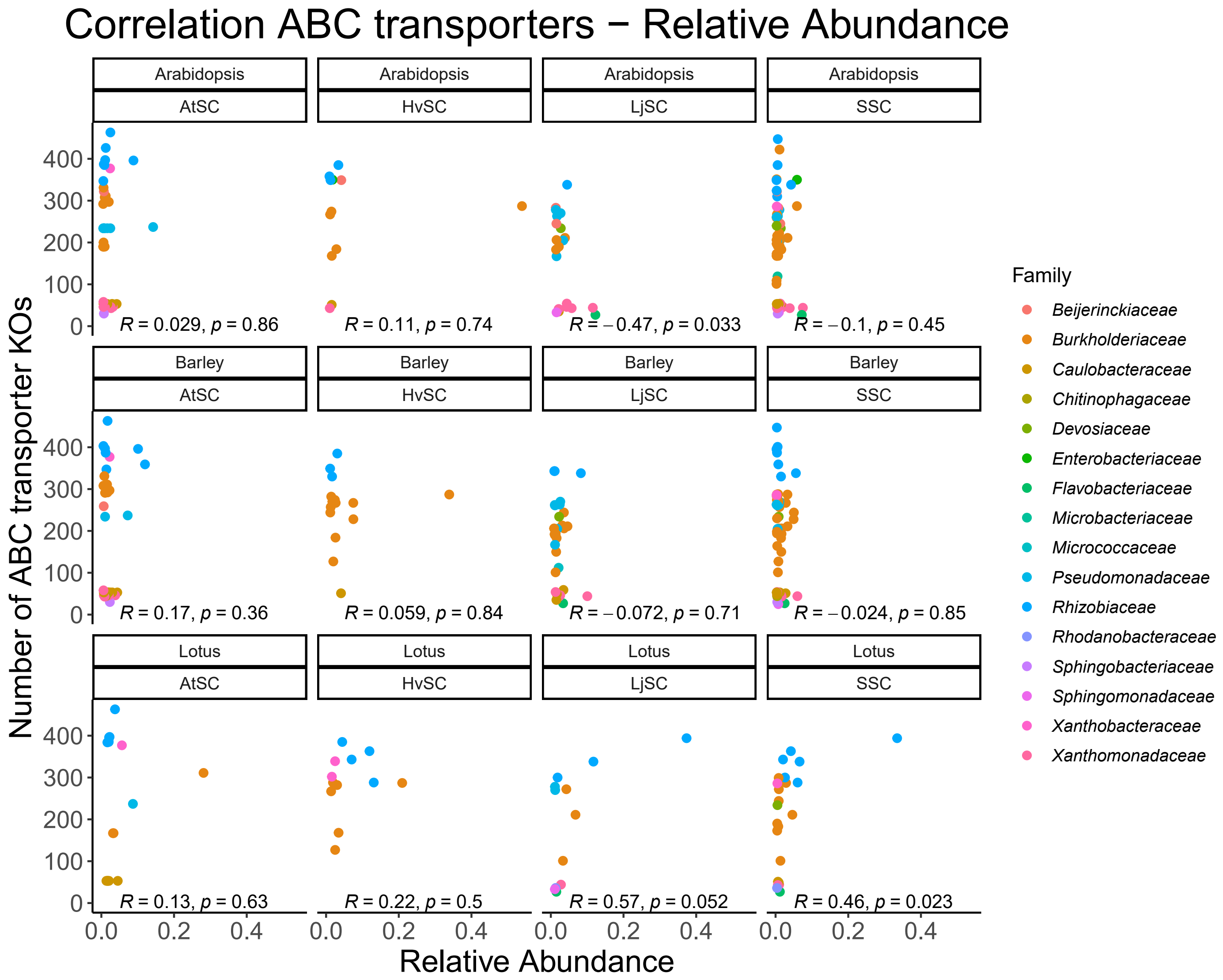

**Figure S32. Correlation ABC transporter diversity and relative abundance colored by family and subsetted per host-inoculum combination.** In each host inoculated with microbial communities, the abundance of the inoculum isolates (X-axis) is compared to the number of ABC transporters in the genome (Y-axis). Correlation coefficients (R) and significance of these correlations (p) are shown in the bottom of the plot.
